# The Dayhoff Atlas: scaling sequence diversity for improved protein generation

**DOI:** 10.1101/2025.07.21.665991

**Authors:** Kevin K. Yang, Sarah Alamdari, Alex J. Lee, Kaeli Kaymak-Loveless, Samir Char, Garyk Brixi, Carles Domingo-Enrich, Chentong Wang, Suyue Lyu, Nicolo Fusi, Neil Tenenholtz, Ava P. Amini

**Author notes:** Corresponding authors. *Email addresses:* (Neil Tenenholtz), (Ava P. Amini). These authors contributed equally (Kevin K. Yang, Sarah Alamdari, Alex J. Lee). Work done while interning at Microsoft Research. Current affiliations: A.J.L: Weill Institute for Neurosciences, UCSF, San Francisco, CA, 94158, USA, and Department of Neurology, UCSF, San Francisco, CA, 94158, USA; K.K.L: Liquid AI, Cambridge, MA, 02142, USA; G.B.: Arc Institute, Palo Alto, CA, 94304, USA, and Department of Genetics, Stanford University, Palo Alto, CA, 94305, USA; C.W.: School of Life Sciences, Westlake University, Hangzhou, Zhejiang, China; S.L.: Institute of Biomedical Engineering, University of Toronto, Toronto, CA.

## Abstract

Organized information powers modern biology, a framework pioneered by Margaret Dayhoff’s Atlas of Protein Sequence and Structure and advanced by today’s databases and computational methods. Here, we extend this paradigm for the AI era, presenting the Dayhoff Atlas of protein sequence data and generative models to accelerate protein biology and design. The Atlas introduces GigaRef, the largest open dataset of natural proteins, spanning 3.34B genomic and metagenomic sequences across 1.70B clusters, and BackboneRef, which distills structural information from 240,811 synthetic backbones into 46M synthetic sequences. Leveraging these datasets, we trained the Dayhoff protein language models, which can predict mutation effects, scaffold structural motifs, and generate novel proteins within families. Training on metagenomic and structure-based synthetic sequences increased the expression rates of generated proteins, demonstrating the value of data diversity and scale. We release the Dayhoff Atlas code, datasets, and models under a permissive license to empower computation in protein design.

## Introduction

In 1965, Margaret O. Dayhoff published the Atlas of Protein Sequence and Structure, which collated the 65 proteins whose amino acid sequences were then known [1]. Today’s protein databases – including UniProt [2] for genomic-derived sequences; metagenomic databases [3, 4, 5, 6, 7]; the Protein Data Bank (PDB) of protein structures [8, 9]; and datasets of multiple sequence alignments (MSAs) of homologous proteins [10] – all descend from this common ancestor.

The creation and public availability of these databases have directly enabled new understanding and engineering of proteins, catalyzed by a parallel revolution in computational methods for biology. Only because of these databases is it now possible to train large-scale deep learning models for proteins, including protein language models (PLMs) that learn the amino-acid language to generate new sequences [11, 12]. PLMs can generalize to new structural topologies [13], predict the effects of mutations [14], generate novel versions of gene editors [15], and directly leverage the evolutionary information in MSAs to generate functional proteins within a homologous family [16, 17].

The performance and downstream utility of PLMs depends directly on their training data [18, 19, 20]. Existing protein databases overlap but have distinct and complementary advantages for developing generative capabilities. Moreover, they have not been unified into a single centralized resource for building foundational models for protein design. UniProt contains sequences reliably inferred from genomes, while metagenomics has the potential to expand the diversity of available sequences because it removes the need to culture organisms before sequencing. Though PLMs have been trained on metagenomic sequences [7, 21], the effects of data composition on performance have not been systematically studied, in part because metagenomic databases have historically not been combined and deduplicated consistently with UniRef. Multiple sequence alignments (MSAs) provide another rich source of protein sequence data. Despite their potential to convey explicit evolutionary information and implicit structural information, the large size of MSAs has led to bespoke and specialized architectures separate from traditional PLMs [22, 23, 17, 24, 25], and has been limited by context length in autoregressive settings [26], preventing modeling of large or longer sets of homologs.

Beyond these abundant sources of protein sequence data, language models for protein design would ideally be complemented by the richness and resolution of protein structure information [27]. Previous approaches for combining information from sequences and structures include repurposing a structure prediction module [28, 29], designing a model architecture that can use both sequence and structure [30], introducing structure-aware tokens [31, 32], or predicting structures from sequence [33], but none of these allow training a simple protein language model on sequences only. Despite the complementarity of protein sequence and structure information, they have not been used together in concert to build highly-capable, general-purpose protein language models that can be deployed across a range of protein design tasks.

Here we present the Dayhoff Atlas of both protein sequence data and generative language models as a centralized resource for protein design (**Fig. 1**). The Dayhoff Atlas provides the largest open datasets of natural proteins and synthetic proteins to date. We first created a giga-scale natural protein dataset, GigaRef, that combines sequences from diverse metagenomic databases with UniRef100 to span both genomic- and metagenomic-derived proteins. To infuse the benefits of structure information into sequence space, we generated a structure-based synthetic dataset, BackboneRef, by using a generative model to sample 240,811 backbone structures and then performing fixed-backbone sequence design to produce 46 million synthetic sequences. Using these datasets, we trained the Dayhoff series of protein language models, including the first trained with structure-based synthetic data augmentation. By leveraging a model architecture that scales efficiently and accommodates long input lengths, we built a PLM which combines single sequences and sets of evolutionarily-related sequences at scale, achieving unified modeling of single sequences, aligned homologs, and unaligned homologs. We then use the trained Dayhoff models to generate DayhoffRef, a dataset of 16 million synthetic protein sequences, demonstrating the potential for generative models to expand the known sequence space. The Dayhoff models make accurate zero-shot predictions of mutation effects, perform guided generation based on evolutionarily-related proteins, and generate short Cas9s that preserve the functional domain architecture, supporting the models’ general-purpose utility. Finally, we performed the first systematic comparison of the effects of model scale and training data on the *in vitro* expressibility of PLM-designed proteins, finding that larger models, metagenomic sequences, and structure-based augmentation all increased expression rates in *E. coli*, while commonly-used *in silico* quality metrics did not discriminate between expressed and unexpressed sequences. We open source the Dayhoff Atlas datasets, model weights, and code to accelerate innovation at the interface of protein biology and computation.

**Figure 1:**
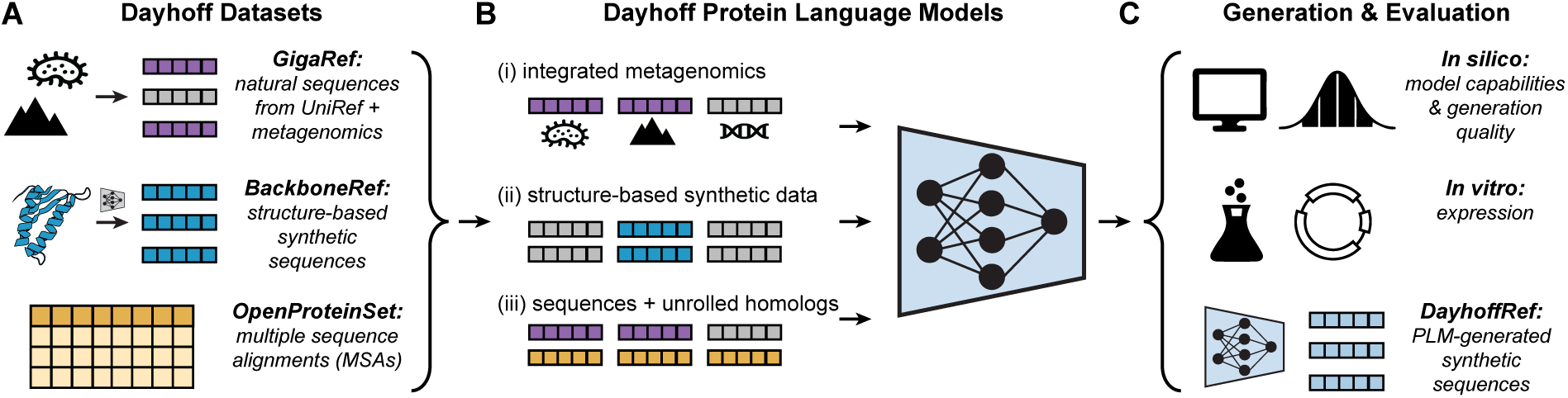
The Dayhoff Atlas of datasets and models for protein sequence generation. **(A)** The Dayhoff datasets: GigaRef (top, grey and purple), a natural protein dataset of 3.34 billion sequences of genomic- (grey) and metagenomic-derived (purple) proteins; BackboneRef (middle, blue), a synthetic dataset created by using a structure-based generative model to sample backbones and then performing fixed-backbone sequence design; and the OpenProteinSet (bottom, yellow) dataset of multiple sequence alignments (MSAs). **(B)** The Dayhoff family of protein language models (PLMs) integrate metagenomics data (i), leverage structure-based synthetic data augmentation (ii), and combine single sequences and sets of homologs at scale (iii), via a long-context hybrid architecture. **(C)** Dayhoff-generated sequences are evaluated both *in silico* and *in vitro*. Scaling generation produces the DayhoffRef dataset of synthetic sequences.

### The Dayhoff Atlas centralizes genomic, metagenomic, and synthetic protein sequences

In creating the Dayhoff Atlas, we aimed to create a comprehensive resource integrating the diversity and scale of genomic and metagenomic sources with the richness of structural information. To this end, we establish the largest open dataset of natural proteins to date, GigaRef, and the first set of synthetic protein sequences derived from *de novo* generated structural backbones, BackboneRef, to centralize information from protein sequences and synthetic structures into a single resource.

#### GigaRef: a natural protein dataset of expanded scale and diversity

We first sought to expand the scale and diversity of natural protein sequences for protein language model training. Most PLMs are trained on protein sequences derived from genomic sequences in GenBank and compiled in UniProt or UniRef. However, this restricts training to organisms from which we can derive whole genomes and excludes a vast diversity of unculturable organisms across diverse evolutionary lineages. Through the recovery of genetic material directly from samples, metagenomics removes the need to culture organisms to obtain genomic information, thus enabling the characterization of the genomes and proteomes of organisms from diverse clinical and environmental contexts. Drawing inspiration from recent work in protein structure prediction [34] and biological language models [35, 18, 7, 21], we reasoned that proteins inferred from metagenomics could enhance dataset scale and diversity. While metagenomic sequences have previously been used to train large-scale biological language models [7] by incorporating their genomic contexts, the effects of systematically combining metagenomic and genomic protein sequences alone have not been studied.

To this end, we systematically integrated metagenomic- and genomic-derived sequences by combining, deduplicating, and reclustering the following 9 databases – MGnify [3], soil metagenome-assembled genomes (SMAG) [36], MetaEuk [37], metagenomic gut virus catalog (MGV) [38], Gut Phage Database (GPD) [39], Soil Reference Catalog (SRC) [6], Marine Eukaryotic Reference Catalog (MERC) [6], Tara Oceans Particle-Associated MAGs (TOPAZ) [40], and UniRef100 – into one dataset suitable for model training: GigaRef (**Fig. 2A**; Table S1, S2). Because the total number of sequences across these datasets (5.7 billion) exceeded the allowable limit of the MMseqs2 clustering software [41] (2^32^ or 4.29 billion), we performed a multi-step clustering process to deduplicate the raw sequences. Starting with the largest datasets, we combined and preclustered MERC and SRC at 70% identity, resulting in 1.81 billion representative sequences, and preclustered MGnify at 70% identity, resulting in 1.67 billion representative sequences (see Methods). We then combined these representatives with sequences from SMAG, MetaEuk, MGV, GPD, TOPAZ, and UniRef100, yielding a total of 3.94 billion sequences. Finally, we clustered these sequences at 90% sequence identity, then clustered the resulting 3.34 billion cluster representatives at 50% identity to produce the GigaRef dataset (**Fig. 2A**).

**Figure 2:**
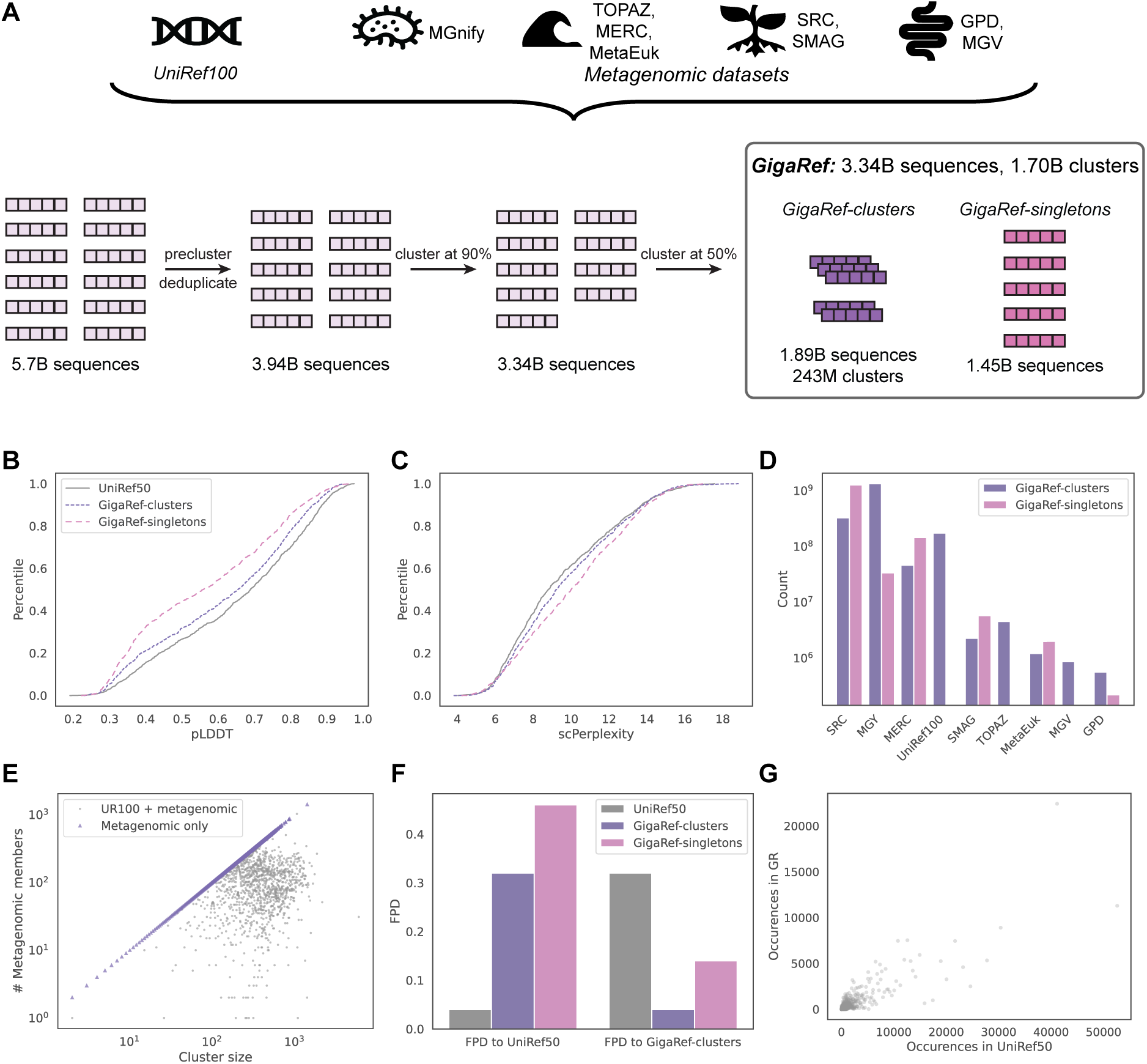
GigaRef is a giga-scale natural protein sequence dataset unifying genomic- and metagenomic-derived sequences. **(A)** GigaRef is created by combining, deduplicating, and reclustering sequences from nine distinct source databases, including UniRef100 and metagenomics datasets covering microbiome, oceanic, soil, and clinical samples. The final GigaRef dataset is comprised of 3.34B sequences spanning 1.70B clusters, with 243M clusters containing multiple sequences (1.89B clustered sequences) in GigaRef-clusters and 1.45B singleton sequences in GigaRef-singletons. **(B-C)** Empirical CDF (ECDF) of pLDDT (B) and scPerplexity (C) values over predicted structures and sequences from UniRef50 (grey), GigaRef-clusters (purple), and GigaRef-singletons (pink). **(D)** Composition of GigaRef clusters (purple) and singletons (pink), as measured by the number of sequences from each source dataset. The absence of a bar represents a count of zero (singletons from UniRef100, TOPAZ, and MGV). **(E)** Scatterplot of the distribution of cluster sizes in GigaRef-clusters (x-axis) versus the number of metagenomic sequences in each cluster (y-axis) over a random sample of GigaRef-clusters (*n*=233). Each cluster is colored by whether it contains both metagenomic and UniRef100 sequences (grey, *n*=123 clusters) or all metagenomic sequences (purple, *n*=110 clusters). **(F)** Distributional distances, as measured by FPD, between UniRef50 and the GigaRef datasets relative to each of the UniRef50 and GigaRef-clusters test sets. **(G)** Number of occurrences of individual Pfam domains in a random subsample (*n*=2048 sequences) of UniRef50 (x-axis) and GigaRef (y-axis) for domains that appear at least once in either dataset.

GigaRef contains c.a. 3.34 billion members across 1.70 billion clusters, 1.45 billion of which contain only one sequence (“singletons”). Following the convention in metagenomics, we focus on clusters with more than one sequence, as these are more likely to contain real proteins [3, 35, 7]. Removing singletons yields 1.89 billion sequences in 243 million clusters – hereafter, “GigaRef” (GR) refers to this dataset of non-singleton sequences (**Fig. 2A**).

GigaRef provides a c.a. 16x increase in the total number of sequences relative to the March 2025 release of UniRef90 and a c.a. 24x increase in the number of clusters relative to the March 2025 release of UniRef50. It has slightly more sequences (3.34 billion vs. 3.3 billion) and clusters with at least 2 sequences (243 million vs. 207 million) than the recent metagenomics database OMG [7], but differs more substantially in composition and construction. Whereas OMG is assembled from two repositories, JGI IMG (contributing both isolate genomes and metagenome assemblies) and MGnify, GigaRef integrates nine metagenomic, environmental, viral, eukaryotic, and curated databases (Table S1), most notably adding UniRef100, a set of curated UniProt-derived proteins absent from OMG. The two corpora are additionally distinct in their clustering approaches (Fig. 2F, S1) Given its scale and the diversity of source datasets used to construct GigaRef, we first sought to characterize the structural plausibility and quality of individual sequences within GigaRef. We evaluated the foldability of individual GigaRef sequences by predicting their corresponding structures using ESMFold [42] and computing the average predicted local distance difference test (pLDDT) across the whole structure. As an additional metric of structural plausibility, we computed a self-consistency perplexity (scPerplexity) by redesigning each predicted structure with the fixed-backbone sequence design algorithm ProteinMPNN [43]. These metrics for randomly-sampled sequences from GigaRef (cluster representatives; no singletons) and GigaRef-singletons revealed that the singletons have significantly lower pLDDT and higher scPerplexity than the cluster representatives (**Fig. 2B-C**, Table S19). As a baseline, we computed the same metrics of structural plausibility for randomly-selected, held-out sequences from UniRef50; these genome-derived sequences exhibit higher pLDDT and lower scPerplexity than GigaRef sequences. We next characterized GigaRef’s composition in terms of the relative representation of its various source datasets (**Fig. 2D-E**). The MERC, SRC, SMAG, and MetaEuk databases are overrepresented amongst the singleton sequences, while there are no singleton sequences from UniRef100, TOPAZ, and MGV (**Fig. 2D**). Out of the 243 million non-singleton clusters, which range in size from 2 to 25,535 members, approximately 231 million clusters do not contain any UniRef100 sequences – that is, they are entirely metagenomic – and c.a. 13 million clusters are a mix of UniRef100 and metagenomic sequences, while no clusters consist entirely of UniRef100 sequences (**Fig. 2E**). This indicates that metagenomic sequences increase diversity by both forming new clusters and augmenting existing ones. Because we did not privilege genomic sequences during the two-stage clustering procedure, 57.5% of UniRef100 sequences were dropped in favor of metagenomic cluster representatives during the 90% clustering stage (Table S2).

Subsequently, we investigated the extent to which GigaRef expands the scope and diversity of natural protein space relative to UniRef. Ideally, GigaRef should shift and expand the distributional coverage of protein sequence, structure, and function space. We thus reasoned that distributional distances over protein embeddings and function annotation predictions could provide proxy metrics to assess GigaRef’s divergence from UniRef. As metrics we used the Fréchet distance (FPD) over ProtBert [35] sequence embeddings, which reflect sequence, structure, and functional information, and the maximum mean discrepancy over function annotations predicted by ProtNote [44] (PNMMD). The FPD and PNMMD between the GigaRef and UniRef50 train sequences were higher than the train-test FPDs and PNMMDs within each of GigaRef and UniRef50, indicating a distribution shift between the two datasets (**Fig. 2F**, Table S3-S4, Fig. S2-S3). The FPD between GigaRef-singletons and UniRef50 was higher than that between GigaRef-clusters and UniRef50, suggesting that singletons are distributionally further from genomic-derived sequences. Inference of protein families present in GigaRef revealed a lower density of Pfam domains relative to UniRef50, though the frequency of occurrences per domain was similar across both datasets (**Fig. 2G**, Table S5). While sequences in UniRef50 averaged 1.58 domains per sequence, GigaRef cluster representatives and singletons averaged 1.1 and 1.0 domains per sequence, respectively. Together, these results indicate that GigaRef provides a unified natural protein sequence database of expanded scale and diversity.

#### BackboneRef: a structure-based synthetic protein dataset

Beyond expanding the number of natural protein sequences, we hypothesized that structure-based synthetic data could implicitly distill the benefits captured in protein structure information – such as stability and biophysical plausibility – into sequence space (**Fig. 1B**). We thus developed a pipeline to generate structure-based synthetic protein data (**Fig. 3A**). We used this pipeline to create the BackboneRef dataset, with the aim of leveraging its synthetic sequences in model training to encourage generation of structurally stable, biophysically plausible new proteins.

**Figure 3:**
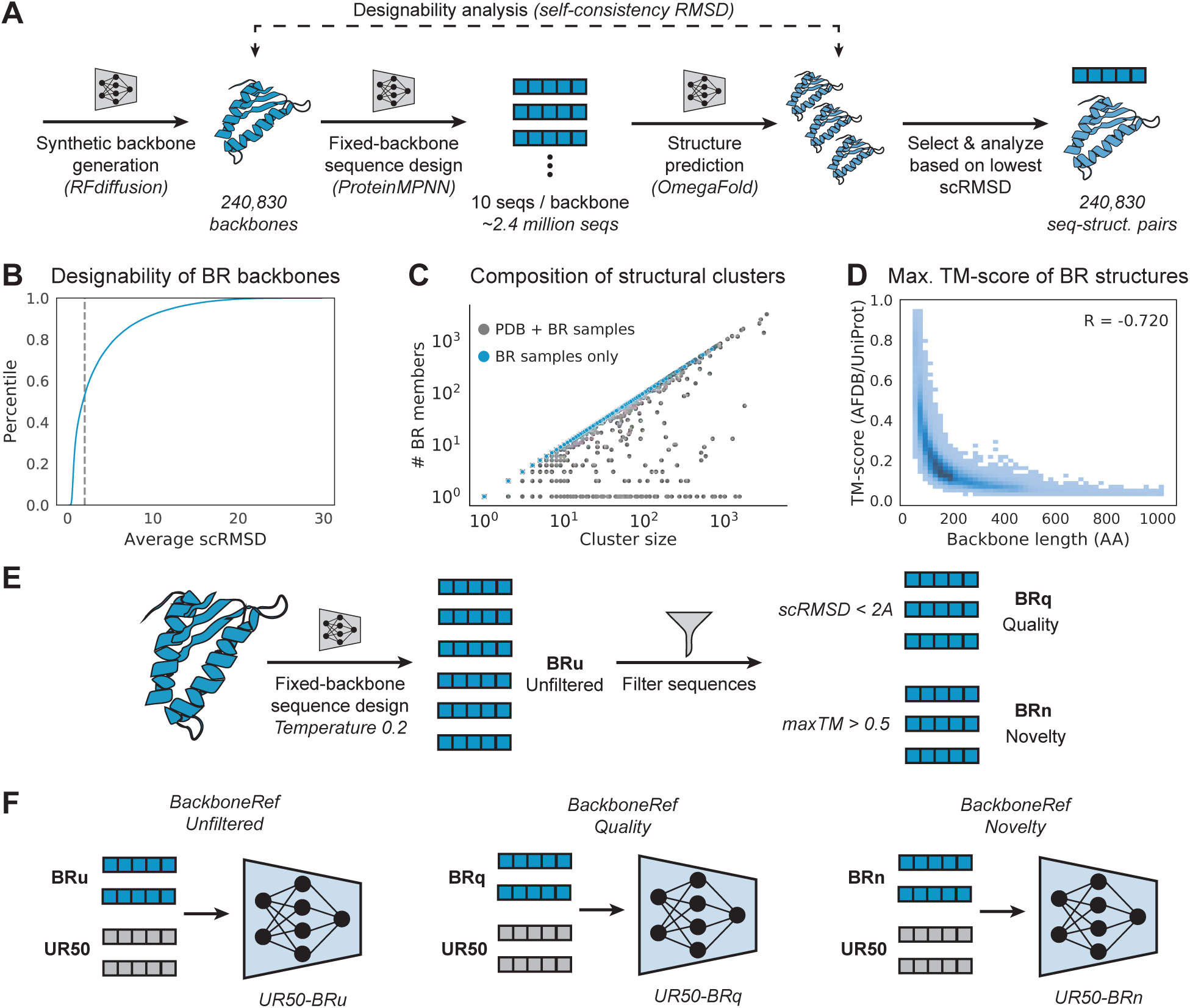
BackboneRef provides designable, novel, and diverse synthetic proteins for structure-based data augmen-tation. **(A)** Workflow for generation and analysis of BackboneRef synthetic structures. **(B)** Empirical CDF (ECDF) of scRMSD for each backbone, averaged over OmegaFold-predicted structures for all 10 fixed-backbone designed sequences. scRMSD is computed between the original backbone and each predicted structure. The dotted line indicates scRMSD = 2. **(C)** Scatterplot of cluster size (x-axis) vs. number of BackboneRef members (y-axis) for each inferred structural cluster in the “PDB + BR” dataset (*n*=105,506 clusters). Each cluster is colored by whether it contains both natural PDB and synthetic BackboneRef samples (grey) or all synthetic BackboneRef samples (blue). **(D)** Distribution of backbone length (x-axis) versus maximum TM-score (y-axis), computed between each backbone’s lowest scRMSD predicted structure searched using Foldseek against predicted structures for all natural proteins in AFDB/UniProt. **(E)** Augmenting PLM training datasets with structure-based synthetic data. Following fixed-backbone sequence design from synthetic backbones (BRu), sequences are filtered on the basis of quality (BRq) or novelty (BRn). **(F)** The resulting sets of sequences are used for data augmentation in PLM training, resulting in Dayhoff-UR50-BRu (left), Dayhoff-UR50-BRq (middle), and Dayhoff-UR50-BRn (right).

We first created a large-scale dataset of 240,811 synthetic backbone structures – to our knowledge, the largest set of generated structures to date – by sampling backbones unconditionally from RFdiffusion [45] (**Fig. 3A**). Secondary structure characterization revealed an enrichment of helical elements, versus disordered and *β*-stranded regions, in the synthetic backbones relative to natural structures (Fig. S4), consistent with prior reports [28, 17, 46]. To evaluate the quality and designability of these backbones, we performed fixed-backbone sequence design for all 240,811 synthetic backbones, sampling 10 sequences per backbone at temperature 0.1 using ProteinMPNN [43] to produce c.a. 2.4 million sequences, which we used to characterize the dataset (**Fig. 3A**). Using OmegaFold, structures were predicted for all resulting sequences to evaluate individual backbone quality via the self-consistency Root Mean Square Deviation (scRMSD) between the original synthetic backbone and the predicted structure (**Fig. 3A-B**). 53.0% of the backbones exhibited an average scRMSD below 2.0, supporting their designability (**Fig. 3B**). We then chose the sequence and predicted structure with the lowest scRMSD for each backbone to characterize the novelty and diversity of BackboneRef proteins.

To characterize the novelty of BackboneRef, we first assessed the extent to which BackboneRef structures include folds not present in natural structures from the PDB [8]. We clustered the lowest scRMSD predicted structures with the PDB using Foldseek [47] (**Fig. 3C**; Table S6). While 99.0% of clusters contained exclusively synthetic or exclusively natural structures, the 1.0% of mixed-membership clusters included 27.8% of synthetic structures. When clustered with the PDB, the 240,811 synthetic backbones yielded 83,121 novel clusters without any natural members (Table S6). Novel clusters had an average of 2.1 (s.d. 10.8) members each, indicating that generative models trained on the PDB can produce many distinct novel structures. We also evaluated the novelty of each individual BackboneRef structure by finding its closest structural match across all natural proteins in UniProt [2] (**Fig. 3D**). 57.3% of synthetic BackboneRef structures had a maximum TM-score less than 0.5 to any natural structure (**Fig. 3D**), consistent with the observation that 72.3% of synthetic structures were in BackboneRef-only clusters (**Fig. 3C**). A rarefaction analysis over the combined BackboneRef and PDB dataset revealed that cluster diversity does not saturate when sub-sampling either across the whole dataset or only BackboneRef (**Fig. S5**), suggesting that scaling could lead to additional novel structures. Together, these results showcase the novelty and diversity of BackboneRef structures both as a whole and individually.

Having characterized the quality and novelty of BackboneRef’s synthetic backbones, we used fixed-backbone sequence design to produce synthetic sequences for downstream augmentation of protein language model training (**Fig. 3E**). To balance diversity and quality of these synthetic sequences, we optimized sequence sampling temperature by performing fixed-backbone sequence design on a random subset of BackboneRef backbones and evaluating the self-consistency of the structures predicted from those sequences (Fig. S6). This analysis identified a steep increase in scRMSD between sequence design temperatures of *T* = 0.2 and *T* = 0.3; thus, we selected *T* = 0.2 as the sampling temperature, sampling 300 sequences per backbone for a total of 72,243,300 sequences. We then down-sampled this dataset for PLM training by randomly selecting 42 sequences per backbone, removing exact duplicates, and then randomly subsampling down to a total of 10M sequences (see Methods); we titled this dataset of backbones and associated sequences ‘BRu’ (BackboneRef, unfiltered).

We next generated filtered subsets of BRu backbones using quality and novelty metrics in order to assess how these attributes impact downstream model performance (**Fig. 3E**). When filtering by quality, we selected the 127,633 backbones with average scRMSD below 2Å and again applied the above sampling procedure (this time initially selecting 80 sequences per backbone) to create another 10M sequence subset; we titled this subset ‘BRq’. When filtering by novelty, we included backbones from BRu with a maximum TM-score to natural proteins in AFDB/UniProt [48] of at most 0.5 and used the resulting 138,044 backbones and above sampling procedure to generate a third 10M sequence subset, which we titled ‘BRn’.

To evaluate BackboneRef’s coverage at the sequence level, we quantified the distribution-level divergence from natural proteins for each set of BackboneRef synthetic sequences. All three BackboneRef sets of synthetic sequences yielded distribution shifts away from natural proteins, as evidenced by FPD and PNMMD metrics being much higher than those between sets of natural sequences (Table S3-S4). Finally, we assessed the protein families present in BackboneRef by computing Pfam annotations for BRu, BRq, and BRn and found that the Pfam domain distributions differed substantially between BR datasets and natural sequences, with Pearson correlations of domain frequencies *<* 0.2 (Table S5). Together, these characterizations suggest that BackboneRef augments protein sequence space with novel, diverse, and structurally plausible structure-based synthetic data.

#### Multiple sequence alignments from OpenProteinSet provide evolutionary information

Multiple sequence alignments (MSAs) are sets of evolutionarily-related (homologous) protein sequences with gaps inserted and unaligned residues deleted such that corresponding residues appear in the same positions. The search and alignment algorithms for constructing MSAs encode rich information about structure and function that is difficult to determine from single sequences. Protein language models trained on MSAs can leverage this evolutionary information to predict the effects of mutations, to sample related sequences, or to perform conditional design [22, 23, 17, 25, 24].

While there have been previous attempts to compute millions of MSAs for training PLMs [22, 23] or structure prediction models [34, 48], these datasets are not publicly available. OpenProteinSet is currently the largest set of pre-computed, publicly-available MSAs, consisting of MSAs for 140,000 unique chains in the PDB and 16M sequences in UniClust30 [10]. We reasoned that MSAs would provide further implicit structural information and explicit evolutionary information, complementing the diversity of GigaRef and novel structure-to-sequence translation of BackboneRef. We thus processed OpenProteinSet to source homologous protein sequences for training the Dayhoff models.

#### The Dayhoff models unify protein sequence datasets and modeling tasks

Having centralized protein sequence datasets within the Dayhoff Atlas, we next sought to unify learning over single sequences and evolutionarily-related homologs at scale in a single generative model (**Fig. 4A-B**). We approached this by “unrolling” each MSA into a series of sequences divided by separator tokens (**Fig. 4B**). However, this unrolling necessitates a model architecture that can accommodate long context – or input – lengths *l*. State-space models (SSMs) efficiently model long contexts by sequentially computing updates to a constant-size state vector [49, 50, 51] but can “forget” information from earlier in the context due to their fixed state size. In contrast, self-attention, notably used within the Transformer architecture [52], uses a *l × l* attention matrix to aggregate information over the context, such that each token can losslessly access information from every previous token at the cost of memory and computational efficiency.

**Figure 4:**
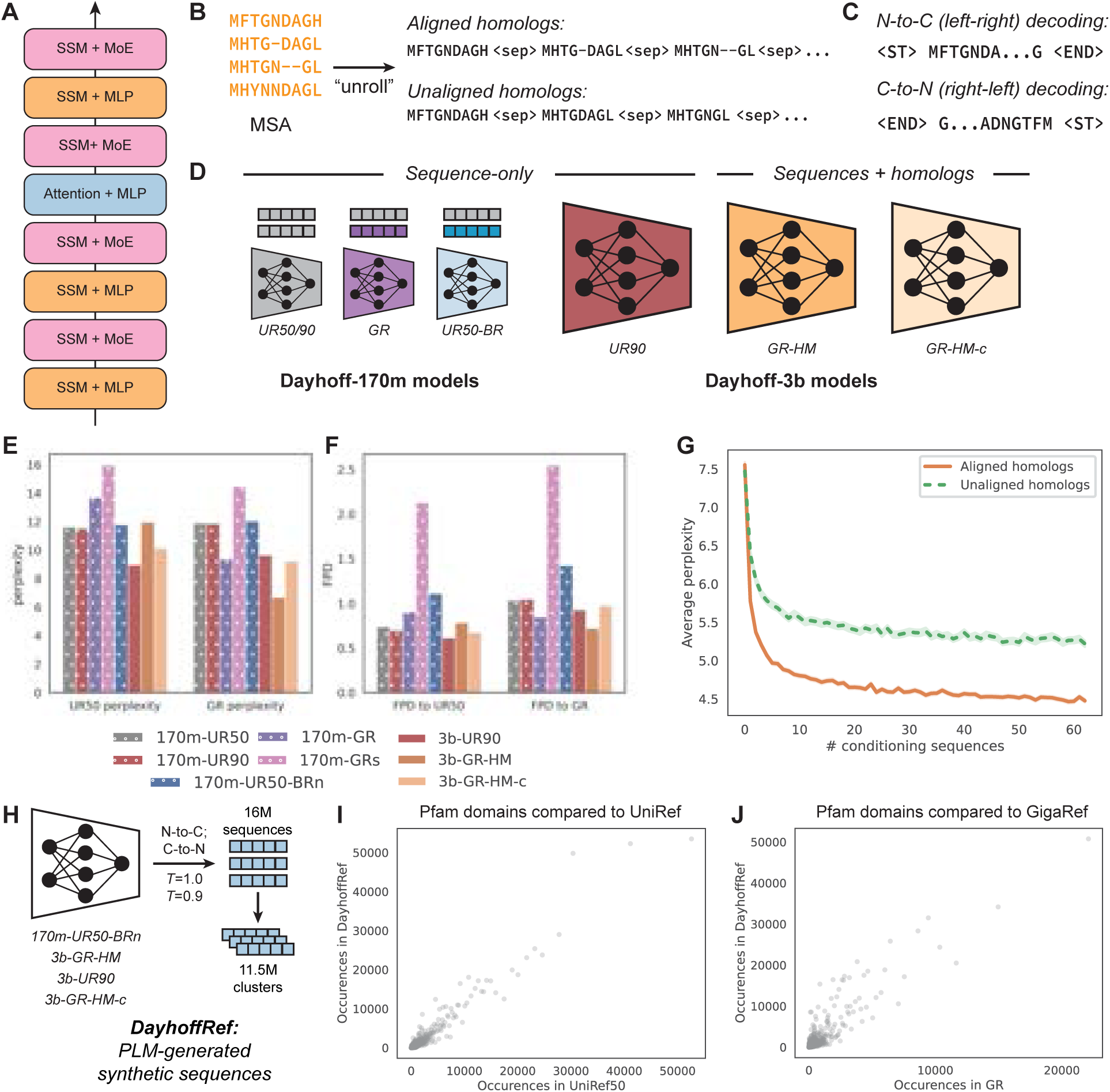
The Dayhoff models unify protein sequence datasets and modeling tasks. **(A)** Schematic of a single block within the Dayhoff model architecture. **(B)** Unrolling MSAs as sets of aligned homologs or of unaligned homologs enables unified modeling of single sequences and sets of homologs. **(C)** The Dayhoff models are trained on both left-right and right-left decoding and can generate proteins in either the N-to-C or C-to-N directions. **(D)** The Dayhoff sequence-only and joint sequence-homolog models across the 170m and 3b parameter scales. **(E)** Perplexities evaluated over held-out cluster representatives from UniRef50 (UR50) and GigaRef-clusters (GR) for each of the Dayhoff models. **(F)** Distributional distances, as measured by FPD, for generations from the Dayhoff models relative to held-out cluster representatives from UR50 and GR. **(G)** Validation perplexity for sets of homologuous sequences as a function of the number of conditioning sequences when homologs are aligned (orange) or unaligned (green). **(H)** DayhoffRef is created by sampling sequences from Dayhoff models in both the N-to-C and C-to-N directions at two temperatures. The 16M generated sequences are clustered at 50% sequence identity to yield a set of 11.5M clusters. **(I-J)** Number of occurrences of Pfam domains in synthetic sequences from DayhoffRef (y-axis) relative to natural sequences from UniRef50 (I, x-axis; Pearson *r*=0.96) and GigaRef (J, x-axis; Pearson *r*=0.89).

Hybrid architectures that combine SSM and transformer attention modules enjoy the benefits of both efficient long-context computation and explicit context retrieval via attention [53, 54]. We therefore utilized a hybrid SSM and transformer architecture as the backbone for all Dayhoff family models (**Fig. 4A**). The architecture most closely resembles that of Jamba, a recent natural language model, which interleaves a single self-attention layer for every seven Mamba SSM layers [54]. Dayhoff, like Jamba, also replaces the multi-layer perceptron in every other layer with a mixture-of-experts module to increase model capacity while minimizing the impact on model cost.

We trained Dayhoff with an autoregressive objective (see Methods), as prior work has shown that this yields better distributional coverage and scaling with model size than discrete diffusion and masked protein language models [17, 55]. However, left-to-right autoregressive models cannot perform order-agnostic infilling, a desired capability for protein design tasks such as inpainting and motif scaffolding. Prior work in language modeling has proposed ways to overcome this limitation, for example by training autoregressive models to “fill in the middle” [56], a method which has been adopted by a few PLMs [25, 24, 21]. Taking inspiration from this approach, we reasoned that we could still achieve any-order infilling by having a model trained both left-right (N-to-C) and right-left (C-to-N), but without explicit fill in the middle, such that arbitrary sub-sequence information could be used as context with decoding then occurring in the appropriate direction. We thus trained the Dayhoff models in both the N-to-C and C-to-N directions by sampling the direction for each sequence during training (**Fig. 4C**).

Using these modeling principles, we trained the suite of Dayhoff PLMs on different datasets from the Dayhoff Atlas (**Fig. 4D**, Table S7) and computed each model’s perplexity and FPD to held-out UniRef50 and GigaRef sequences in order to measure how well it learned each distribution of natural proteins (**Fig. 4E-F**, Tables S8-S18). Across all models, the perplexity decreased with longer contexts and more training in both the N-to-C and C-to-N directions, confirming that models learned to generate successfully in both directions (Fig. S7). To evaluate the structural fidelity of generations, we computed the ESMFold pLDDT and scPerplexity for generated sequences from each model (Tables S19-S29). Sequences from all models had similar pLDDTs and scPerplexities to each other, and all average pLDDTs for generated sequences were lower than for natural sequences while all average scPerplexities were higher. There were no statistically significant differences between the pLDDTs and scPerplexities when generating in the N-C versus C-N direction, further corroborating the generative ability of the Dayhoff models.

#### Sequence-only models elucidate the effects of dataset composition on downstream performance

Each of the Dayhoff PLMs enables us to systematically test a specific hypothesis about the influence of training data composition on model performance (**Fig. 1B, 4D**). We first investigated the effects of increasing the number of natural sequences by training 170 million-parameter, sequence-only models. Following previous work [57, 17], the base model was trained on the UniRef50 cluster representatives (Dayhoff-170m-UR50). Next, we expanded the genomic data by sampling one UniRef90 sequence per UniRef50 cluster per epoch (Dayhoff-170m-UR90). As expected from previous work [57], increasing the within-cluster diversity improved the perplexity and FPD on held-out UniRef50 cluster representatives (**Fig. 4E-F**, Table S8, S9-S10). To evaluate the effect of metagenomic data, we trained a model by sampling one sequence per GigaRef cluster per epoch (Dayhoff-170m-GR). This improved model performance on held-out GigaRef sequences at the expense of UniRef50 performance (**Fig. 4E-F**, Table S8, S9-S11).

To systematically evaluate the effects of structure-based synthetic data augmentation, we next trained models on the UniRef50 cluster representatives augmented by each of BRu, BRn, and BRq (Dayhoff-170m-UR50-BRu, Dayhoff-170m-UR50-BRn, and Dayhoff-170m-UR50-BRq, respectively). Structure-based synthetic data augmentation resulted in only slightly worse cross-entropy on the UniRef50 validation set (respective perplexities of 11.78, 11.67, 11.66 for Dayhoff-170m-UR50-BRu, Dayhoff-170m-UR50-BRn, Dayhoff-170m-UR50-BRq vs. 11.62 for Dayhoff-170m-UR50), indicating that these models still learned the natural sequence distribution (**Fig. 4E-F**, Table S8). However, the FPD was higher for sequences generated from models trained using sequences from BackboneRef (**Fig. 4E-F**, Table S8, S13-S15), indicating that structure-based synthetic data augmentation results in an appreciable distribution shift in generated sequences relative to natural proteins.

#### The Dayhoff models unify single-sequence and evolutionary modeling

We ultimately sought to train a single model that learned over both single protein sequences and sets of evolutionarily-related homologous sequences, in order to unify generative modeling of protein sequences at scale (**Fig. 4B,D**). As a step towards this, we scaled the single-sequence model size to 3 billion parameters and its training to UniRef90 (Dayhoff-3b-UR90) and found that this improved both perplexity and FPD (**Fig. 4E-F**, Table S8, S16). We then leveraged the Dayhoff Atlas datasets to source single sequences from GigaRef together with evolutionarily-related homologous sequences from the MSAs of OpenProteinSet. Homologs can be modeled as aligned sequences (with gaps inserted and unaligned residues deleted) or as unaligned, variable-length sequences. Aligned sequences directly encode information about which positions are evolutionarily related. However, the alignment (i.e., gap positions and deleted residues) returned by any alignment algorithm depends on hyperparameters and may be just one of several alignments of similar quality. Previous works have explored the impact of training on unaligned homologs [23, 25]. We thus used unaligned homologs to enable models to learn which positions are related between homologs without the biases of a pre-computed alignment. Furthermore, the ability to model unaligned sequences enables likelihood computations over insertions and deletions (a.k.a. “indels”) without having to recompute the MSA. Training on unaligned homologs would additionally enable homolog conditioning at generation time while reducing dependence on the specific alignment used.

To this end, we sourced unaligned homologs from the OpenProteinSet MSAs, unrolled the MSAs to obtain series of aligned sequences, and provided sets of unaligned sequences and aligned sequences at equal ratios during training (**Fig. 4B**). We first trained our joint sequence-homolog model on an equal mix of GigaRef sequences and OpenProteinSet homolog sets (Dayhoff-3b-GR-HM). Dayhoff-3b-GR-HM was able to model both single sequences and homologous sequences effectively, as demonstrated by its perplexity and FPD on held-out GigaRef sequences (**Fig. 4E-F**, Table S17) and its perplexity on held-out OpenProteinSet homologs, respectively (Table S8).

Based on the premise that UniRef sequences are higher quality than metagenomic sequences, we continued training the combined sequence-homolog model on UniRef90 and OpenProteinSet, but not GigaRef, with a cooldown learning rate cycle to help the model stabilize its predictions (Dayhoff-3b-GR-HM-c). This brought the perplexity and FPD on held-out UniRef50 sequences very close to that of the single sequence-only Dayhoff-3b-UR90 model while further improving the perplexity on both aligned and unaligned homologs at the cost of slightly worse performance on GigaRef (**Fig. 4E-F**, Table S8, S18). This tradeoff reflects the distinct advantages of each data source: GigaRef training produces generations that are distributionally closer to the broader metagenomic sequence space, while cooling on UniRef recovers proximity to the curated genomic sequence distribution For sets of aligned and unaligned homologs, the average perplexity per sequence decreased as more sequences were added to the context, confirming that Dayhoff can utilize the long context and that additional context improves performance (**Fig. 4G**). Together, these results demonstrate that the Dayhoff sequence-homolog PLMs unify modeling of single protein sequences, aligned homologs, and unaligned homologs into a single generative model.

#### Scaling generation produces the DayhoffRef dataset of synthetic protein sequences

Given the potential for generative models to expand the space of proteins and their functions, we next used the Dayhoff models to generate DayhoffRef, a PLM-generated database of synthetic protein sequences, containing over 16 million sequences derived from the Dayhoff-170m-UR50-BRn, Dayhoff-3b-UR90, Dayhoff-3b-GR-HM, and Dayhoff-3b-GR-HM-c models (**Fig. 4H**, Table S30). The overall sequence diversity of proteins in DayhoffRef was greater than that of UniRef50, as evidenced by the fact that clustering DayhoffRef at 50% sequence identity yielded an average cluster size of 1.45 sequences with 56% of clusters being singletons, in contrast to UniRef50’s average cluster size of 2.90 sequences and singleton proportion of 16.5% (Table S31). Still, DayhoffRef’s distribution of function annotations more closely recapitulated that of natural proteins than of structure-based synthetic sequences (Table S4), and the diversity of sequences in DayhoffRef effectively captured the natural distribution of Pfam domains (**Fig. 4I-J**, Pearson *r* = 0.96 relative to UniRef50, *r* = 0.89 relative to GigaRef) despite fewer domains per sequence in DayhoffRef (0.77) than in UniRef50 (1.58). Overall, DayhoffRef achieves a natural-like functional distribution, with substantially greater sequence diversity. We make DayhoffRef openly available as part of the Dayhoff Atlas to enable continued study and utilization of AI-generated protein sequences.

### The Dayhoff models are general-purpose protein language models

Having trained our models across different datasets and scales, we demonstrated their general-purpose utility by applying them across three scenarios key to protein engineering and design: prediction and scoring of wet-lab measurements of function, scaffolding functional motifs or domains, and guided generation based on homologous or functionally-related proteins (**Fig. 5A**). These evaluations span both predictive sequence-function tasks as well as generative functional design applications.

**Figure 5:**
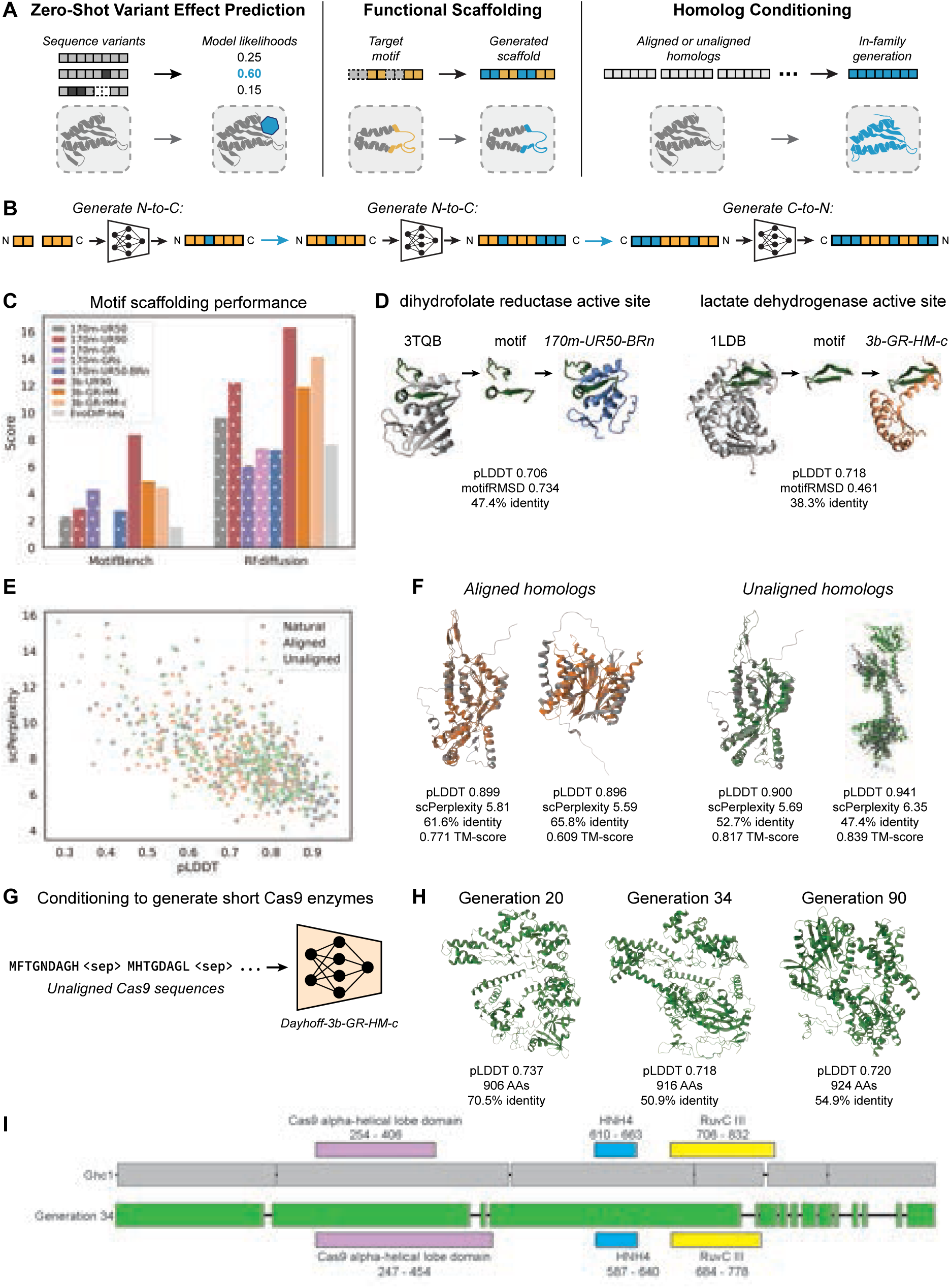
The Dayhoff models are general-purpose protein language models. **(A)** The Dayhoff models enable zero-shot mutation effect prediction, motif scaffolding, and flexible, family-conditioned generation. **(B)** Schematic of approach for any-order inpainting and scaffolding with autoregressive generation. **(C)** MotifBench scores on the MotifBench and RFdiffusion scaffolding benchmarks. **(D)** Scaffolding of the active sites of dihydrofolate reductase by Dayhoff-170m-UR50-BRn (left) and of lactate dehydrogenase by Dayhoff-3b-GR-HM-c (right). The ESMFold-predicted structure and computed metrics for the successful designs are shown, with the scaffolded motif in green, the ground truth structures in dark grey, and the predicted structures of the generated sequences shown in blue and orange for Dayhoff-170m-UR50-BRn and Dayhoff-3b-GR-HM-c, respectively. **(E)** Sequence pLDDT (x-axis) versus scPerplexity (y-axis) for natural query sequences from held-out OpenProteinSet MSAs, (grey, *n*=226), generated by Dayhoff-3b-GR-HM-c via aligned homolog conditioning (orange, *n*=226), and generated by Dayhoff-3b-GR-HM-c via unaligned homolog conditioning (green, *n*=226). **(F)** Predicted structures and metrics for selected proteins from Dayhoff-3b-GR-HM-c, generated via aligned homolog conditioning (orange) or unaligned homolog conditioning (green). **(G)** To generate short variants of Cas9, a set of unaligned Cas9 sequences was provided as conditioning input to Dayhoff-3b-GR-HM-c, and new sequences were generated. **(H)** Predicted structures and metrics for select generations produced via the procedure in (G). Sequence identities are relative to the closest natural Cas9. **(I)** Sequence alignment for the Generation 34 short Cas9 relative to the closest natural Cas9 by sequence identity, with functional domains labeled.

#### The Dayhoff models are strong zero-shot predictors of mutation effects

As a result of training on evolutionary-scale sequence datasets, protein language models can be used for *in silico* prediction of experimental measurements of protein function, based on the observation that PLM-computed likelihoods often correlate with sequence fitness scores [14]. Because it models sequences autoregressively, Dayhoff can naturally compute likelihoods for protein variants with indels in addition to those with only substitutions, unlike masked language models like ESM and structure-based models. This “zero-shot prediction” ability – scoring variants for fitness without any additional model training – across both indels and substitutions offers the flexibility to rapidly score candidate variants *in silico* to guide downstream protein engineering campaigns.

We evaluated the Dayhoff models on each variant in ProteinGym [58], a standardized set of benchmarks for predicting the effects of protein mutations on a range of experimental measures of fitness, including activity, binding, and stability. Given the same training data, average zero-shot scores on both the substitution and indel portions of ProteinGym improved as the model size increased (Table 1). Training on GigaRef generally decreased performance as compared to training on UniRef, suggesting that metagenomic sequences are more distant from the sequences and variants represented in ProteinGym. The Dayhoff-3b models demonstrated robust performance on both substitutions and indels. Dayhoff-3b-GR-HM-c outperformed every single-sequence model on ProteinGym-substitutions except for Vespa [59] and VespaG [60], which are trained on MSA scores, and Dayhoff-3b-UR90 was the second best model overall on ProteinGym-indels behind PoET [23] (as of evaluations in June 2025). Dayhoff-3b-GR-HM-c achieved strong zero-shot performance when scoring single sequences, suggesting that training on homologs provided beneficial representations. Surprisingly, providing aligned or unaligned homologs at inference time decreased performance relative to single-sequence scoring (Table 1). In the single-sequence setting, we average likelihoods in both the N-to-C and C-to-N directions, which may recover bidirectional scoring performance. In contrast, for homolog-conditioned scoring, we subsample the parent MSA and average the likelihoods, which does not afford equivalent bidirectional coverage. Additionally, because the model is trained to generate family-consistent sequences given homolog context, the subsampled MSA directly biases the downstream likelihood toward the consensus of the conditioning set. Notably, for the generation setting where homolog conditioning is most naturally applicable, it substantially improves output quality (Fig. 5E, S10-S13, Table S36)

**Table 1:**
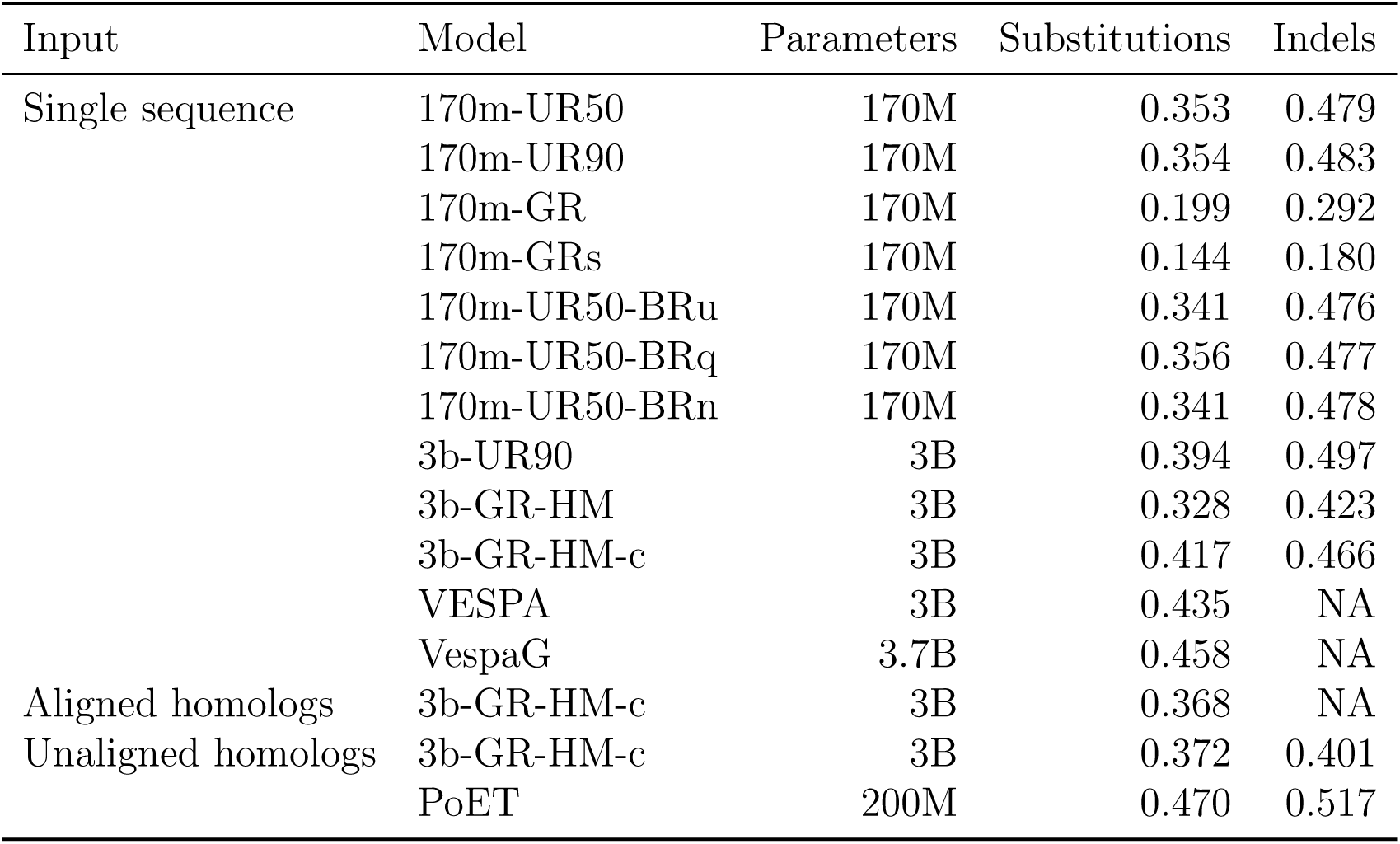
Zero-shot performance (Spearman’s correlation coefficient) on ProteinGym substitutions and indels.

These results reveal a tradeoff between dataset diversity and zero-shot fitness prediction: training on GigaRef decreased ProteinGym performance relative to UniRef, likely because metagenomic sequences are more distant from the well-characterized proteins and variants represented in ProteinGym. Cooling on UniRef90 recovered and surpassed single-sequence performance, suggesting that combining diverse pre-training with a focused cooling phase can capture the benefits of both broad and curated datasets. The strong overall performance of Dayhoff-3b-GR-HM-c supports its utility for zero-shot mutation effect prediction in the single-sequence setting. Stratifying ProteinGym by functional category confirmed that these trends are consistent across selection types. Scaling from 170M to 3B parameters significantly improved zero-shot variant-effect prediction, particularly for stability, activity, and organismal fitness (paired Wilcoxon, p < 0.01). Metagenomic augmentation degraded performance across nearly all categories at both scales (p < 0.001), but cooldown training on curated sequences largely reversed this penalty: Dayhoff-3b-GR-HM-c recovered to statistical parity with Dayhoff-3b-UR90 across all categories while retaining the diversity benefits of metagenomic pretraining. BackboneRef augmentation, despite its benefits for generation quality and expressibility, did not significantly improve zero-shot scoring in any category (Supplementary Figure S8, S9).

#### The Dayhoff models can scaffold structural motifs

We next deployed the Dayhoff models on an explicit generative design task – generating a protein scaffold around a desired structural motif (**Fig. 5A**), motivated by the fact that many protein functions, including receptor-ligand interactions, enzymatic activity, and epitope recognition, are mediated by small functional motifs. Importantly, motifs may not be contiguous in sequence space. Structure-based models [45] condition on the motif by scaffolding directly in 3D space, while sequence diffusion models [17, 61, 62] use order-agnostic generation. PLMs must be able to account for and leverage these sequence discontinuities in order to perform motif scaffolding. However, autoregressive language models like Dayhoff can only generate in a fixed direction, despite their ability to better learn the underlying sequence distribution relative to diffusion models [17]. We thus developed an approach to perform arbitrary sequence infilling autoregressively (**Fig. 5B**), capitalizing on the fact that the Dayhoff models are trained to do both N-to-C and C-to-N generation. We scaffolded motifs by sampling the number of residues between motif segments and then simply generating in the N-to-C direction conditioned on the most N-terminal motif segment, placing any further motif residues as they were encountered. The sequence was then completed by generating in the C-to-N direction conditioned on the motif and the partially generated scaffold, up to a pre-specified length (see Methods).

We used this approach to generate scaffolds across diverse, challenging motif-scaffolding problems from MotifBench [63] and RFdiffusion [45] (30 and 25 problems each, respectively; **Fig. 5C**, Table S32-S35). The problems include enzyme active sites, protein-protein binding interfaces, receptor-ligand interactions, and recognition sites for viral epitopes. During generation, only the motif’s amino-acid sequence was supplied as input information, and each Dayhoff model was used to sample 100 candidate sequences per problem. We compared Dayhoff against EvoDiff-seq, a sequence-based diffusion model explicitly trained on order-agnostic generation [17], and evaluated performance by predicting structures for generated sequences and counting a generation as “successful” if the structure pLDDT was greater than 0.7 and if the predicted motif coordinates had less than 1Å RMSD relative to those of the true structure. To assess a model’s ability to provide unique solutions across a range of scaffolding problems, we leveraged the MotifBench score [63], which ranges from 0 (no successes) to 100 (every generation is a unique success) and is designed to account the fact that it is preferable for a model to provide few solutions to more problems, rather than more solutions to fewer problems (Table S32-S35).

The Dayhoff models outscored EvoDiff-seq on both sets of motif-scaffolding problems, (**Fig. 5C**, Table S32-S35), confirming that order-agnostic generation is not necessary for motif scaffolding. EvoDiff-MSA, which conditions on a multiple sequence alignment of the target, scored higher than all Dayhoff models (Table S34, S35), consistent with the additional evolutionary context available to it. Increasing model size generally improved performance, with the best overall model being Dayhoff-3b-UR90. For the 170m-parameter models, training on metagenomic sequences resulted in better scaffolding scores, while structure-based synthetic data augmentation with BackboneRef did not. We visualized predicted structures for selected successful scaffolds for two distinct enzyme active sites, lactate dehydrogenase and dihydrofolate reductases, noting their high predicted pLDDTs and low sequence identity to any natural protein despite sub-angstrom motif RMSDs (**Fig. 5D**).

#### The Dayhoff joint sequence-homolog models can use evolutionary conditioning to generate plausible variants

By learning over unrolled sets of homologous sequences, the Dayhoff joint sequence-homolog models enable guided generation by simply taking related sequences as input context at inference time (**Fig. 5A**). Generation conditioned on homologs without family-specific training has previously been demonstrated using non-aligned sequence sets [23, 25], and multiple sequence alignments [17]. In contrast to these approaches, the Dayhoff models support homolog-conditioned generation within a single model that also performs single-sequence scoring and unconditional generation and that can condition on either aligned or unaligned homologs. We thus tested the ability of Dayhoff-3b-GR-HM-c to generate plausible candidates of a protein family in this unified setting. Using 226 held-out MSAs, we regenerated the query sequence conditioned on up to 56 randomly-chosen aligned or unaligned homologs (see Methods). As baselines, we generated from EvoDiff-MSA [17], Prot-xLSTM [24], and CCMGen [64] using the same conditioning sequences (note that our held-out MSAs may be in the training sets for EvoDiff-MSA or Prot-xLSTM). We then predicted structures for each generation and evaluated structural plausibility using pLDDT and scPerplexity (**Fig. 5E**, S10, S11), structural similarity to the original query using TM-score (Fig. S12), and sequence novelty using identity to the most similar conditioning sequence (Fig. S13). **Fig. 5F** shows examples of generated queries aligned to the original queries, and performance metrics are summarized in Table S36.

As expected, generating with evolutionary conditioning produced sequences with much higher pLDDT and lower scPerplexity than unconditional generation with the same model (Table S29, S36). Conditioning Dayhoff with unaligned homologs resulted in sequences that were more structurally-plausible than those from conditioning with aligned homologs (**Fig. 5E**, S10, S11). Relative to using aligned homologs, Dayhoff with unaligned homologs generated sequences that were a closer structural match to the original query and more novel relative to the conditioning sequences (Fig. S12, S13). Compared to Prot-xLSTM, which uses unaligned homologs, Dayhoff with unaligned homolog conditioning generated sequences with higher structural plausibility but slightly less sequence novelty. Likewise, when conditioning with aligned homologs, Dayhoff produced more structurally-plausible sequences with less sequence novelty compared to EvoDiff-MSA and CCMgen. These results demonstrate that Dayhoff can generate novel and structurally-faithful sequences with evolutionary conditioning, and that conditioning on unaligned homologs may yield more plausible and diverse generations than conditioning on alignments.

To demonstrate the utility of leveraging homologs in-context, we applied evolutionary conditioning to generate short, plausible Cas9 proteins (**Fig. 5G**). Shortening gene editors like Cas9 can improve their delivery efficiency, tissue penetration, and safety profile [65]. Beginning with 798 natural Type 2 Cas9 proteins ranging in length from 961 to 1809 [66], we clustered at 90% identity to reduce redundancy and removed sequences longer than 1050 residues, resulting in 45 clusters containing 93 natural, short Cas9s. We chose random subsets of 32 conditioning sequences from different clusters, provided these sequences as input context to Dayhoff-3b-GR-HM-c, and repeated until we generated 100 new sequences between 801 and 960 residues in length (**Fig. 5G**). We predicted structures for the 100 generated sequences using ESMFold and identified the most similar natural Cas9 to each generated sequence. 27 generations had ESMFold pLDDT > 0.7; of those, 21 contained large (>20 residue) C-terminal truncations compared to the closest natural Cas9. Of the 6 remaining generations, we randomly selected 3 for further analysis.

These 3 sequences range from 906 to 924 residues, have 51-71% sequence identity to the closest natural Cas9, and are between 114 and 122 residues shorter than the closest natural Cas9 (**Fig. 5H, S14**). To determine their domain architecture and to analyze the locations of deletions, we assigned Pfam domains to each generated sequence and its closest natural Cas9 using InterPro [67] and realigned the sequences for visualization (**Fig. 5I**, S14). Each sequence has a similar domain architecture to its closest natural Cas9 and possesses domains consistent with Cas9 function, such as the HNH endonuclease HNH4 and RuvC III. There is no consistent pattern of deletions, with deletions spread throughout the three generations as opposed to always truncating the N- or C-terminal domains. Notably, Dayhoff-3b-GR-HM-c is able to do this without finetuning, in contrast to previous attempts to use PLMs to generate novel Cas9s [15]. This example highlights Dayhoff’s ability to condition on sequences in-context to generate plausible candidates of a particular family and with desired attributes as measured *in silico*. Experimental validation of endonuclease activity, PAM recognition, and specificity for these shortened Cas9 candidates remains an important next step.

### Impact of scale, metagenomic data, and structure-based augmentation on expression of generated proteins

Beyond *in silico* evaluations, a capable protein language model should generate proteins that can be expressed *in vitro* in cellular systems, as this reflects the biological plausibility and stability of the synthetic proteins. We therefore sought to validate generations experimentally and, in doing so, to establish a ground-truth set of expression outcomes against which we could later assess how well common *in silico* metrics predict expressibility. Based on the assumption that learning from genomic-derived sequences results in the highest quality generations, we first manually-curated 25 Dayhoff-3b-UR90 generations, nominated as likely to express (see Methods), for *in vitro* validation. Generations were expressed in two *E. coli* strains each (BLR(DE3) and BL21-AI, see Methods); a protein was considered to be expressed successfully if expression in at least one strain was detected at the correct molecular weight via SDS-PAGE. 16/25 sequences expressed in at least one strain and had reasonable secondary structure characteristics (Fig. S15), indicating that Dayhoff can generate proteins that express and are biophysically sound.

Given the successful expression of manually-nominated sequences from Dayhoff-3b-UR90, we set up an unbiased head-to-head comparison to examine the effects of model scale and training data composition on the expressibility of generated sequences (**Fig. 6A**). We generated 200 sequences from each of Dayhoff-170m-UR90, Dayhoff-170m-GR, Dayhoff-170m-UR50-BRn,

**Figure 6:**
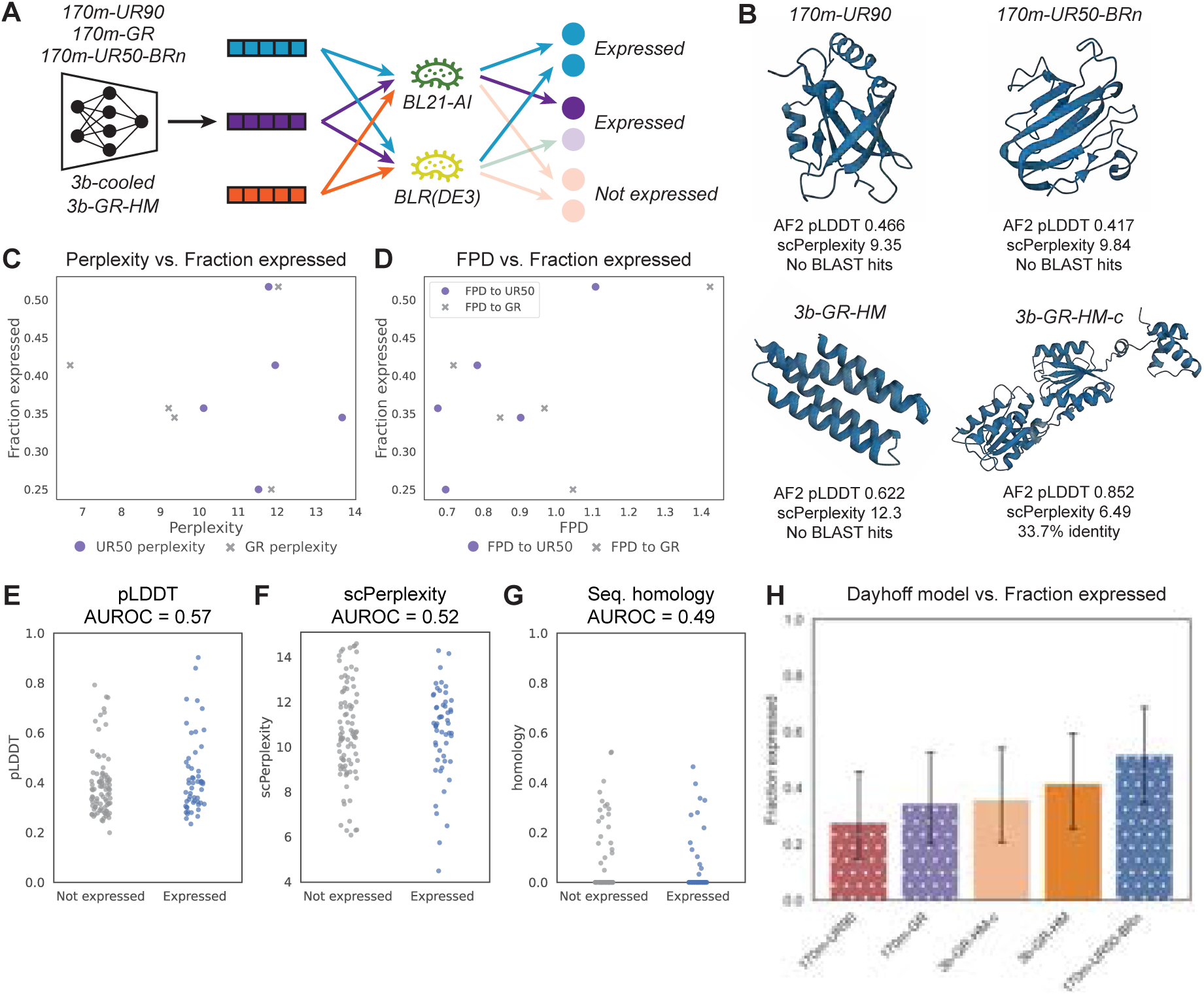
Effects of model scale and dataset composition on *in vitro* expressibility. **(A)** Unconditional generations from the Dayhoff models across the 170m and 3b scales were expressed in each of the BL21-AI and BLR(DE3) *E. coli* strains. A protein was measured as an expression success if it expressed successfully in either or both of the two strains. **(B)** Predicted structures and metrics for select successfully-expressed proteins. **(C-D)** Fraction of proteins successfully expressed versus test-set perplexities (C) and FPD (D) against UniRef50 (UR50) and GigaRef (GR) across Dayhoff model variants. **(E-G)** Distributions of pLDDT (E), scPerplexity (F), and sequence homology to closest natural protein hit (G; BLAST over NCBI NR protein database) for unconditional generations tested for *in vitro* expressibility in *E. coli*, with distributions split between proteins that did (blue) or did not (grey) express. The AUROC value for binary classification based on each metric is provided. **(H)** Fraction of proteins successfully expressed for unconditional generations from Dayhoff model variants (*n*=29 per model except for *n*=28 for Dayhoff-3b-GR-HM-c). with Wilson 95% confidence intervals denoted

Dayhoff-3b-GR-HM, and Dayhoff-3b-GR-HM-c; selected 29 sequences 60-1500 residues long from each set, except for 28 sequences from Dayhoff-3b-GR-HM-c; and tested these *in vitro*. To enable an unbiased comparison, no filtering based on predicted structural metrics or sequence homology was performed, although we did filter out sequences predicted to have signal peptides, mitochondrial transit peptides, or transmembrane helices (see Methods). A total of 55/144 (38.2%) of sequences expressed successfully, with selected expressed proteins visualized in **Fig. 6B**. While several metrics have been proposed to evaluate the quality of machine-generated proteins *in silico*, it has yet to be shown whether any of these metrics are actually predictive of *in vitro* expression and function. We leveraged our scaled, unbiased expression screen to assess this and found that, surprisingly, neither model-level metrics, including perplexity and FPD (**Fig. 6C-D**), nor sequence-level metrics, including ESMFold pLDDT, scPerplexity, and homology to the closest BLAST [68] hit (**Fig. 6E-G**), were predictive of successful expression in *E. coli*. In particular, many of the successful generations had no BLAST hits (**Fig. 6B**). These results not only demonstrate that Dayhoff can generate novel proteins that express successfully, but also reveal the limitations of current *in silico* metrics in predicting *in vitro* expressibility.

Despite the fact that neither model-level nor sequence-level metrics correlated with successful expression, we aimed to explore differences in model scale and training data composition on expression rates. We assessed the expressibility of proteins generated by each model variant (N = 28–29 per model). Expression rates ranged from 28% (Dayhoff-170m-UR90, 8/29) to 52% (Dayhoff-170m-UR50-BRn, 15/29) and were reported with Wilson 95% confidence intervals (**Fig. 6H**). We note that the sample size of this study left it underpowered to rank models by expressibility, as indicated by the estimated confidence intervals and no significant difference across the five model variants (Pearson’s chi-squared test of independence, *χ*^2^ = 3.998, *df* = 4, *p* = 0.41).

## Discussion

High-quality, diverse, and well-curated data is critical for training generative models of proteins. However, there is currently no resource that centralizes the complementarity of genomic, metagenomic, evolutionary, and structure-based information to enable the development of robust, general-purpose protein language models. We created the Dayhoff Atlas of both datasets and protein language models to serve this need, establishing large-scale, open-source datasets of both natural sequences (GigaRef) and synthetic sequences (BackboneRef, DayhoffRef) and a suite of PLMs that systematically unify protein sequence datasets and modeling tasks. Through a controlled evaluation of the Dayhoff models across downstream tasks, we compare the effects of model scale and training dataset composition on the performance and utility of PLMs. Our results demonstrate that novel, high-quality metagenomic and synthetic data can induce protein language models to generate distributions that are further from the distribution of genomic sequences.

Notably, our results suggest that these macroscopic design choices – such as training dataset composition or model size could yield differences in *in vitro* expression in *E. coli*. However, additional studies at greater scale and statistical power are needed to validate these initial observations. Further, we find that common *in silico* metrics of generation quality (i.e., pLDDT) and model learning (i.e., perplexity) are not predictive of expressibility. The failure of in silico metrics to predict expression may reflect several factors: (1) metric limitations, as pLDDT and scPerplexity assess structural confidence and designability, which are necessary but not sufficient for in vitro expression; (2) biological complexity, as expression success depends on translation efficiency, folding kinetics, proteolytic stability, and aggregation propensity, none of which are directly captured by structure prediction confidence; and (3) assay context, as expression in E. coli imposes host-specific constraints that are orthogonal to predicted structural quality. These results underscore the importance of experimental validation and motivate the further development of expression-aware metrics. These results motivate critical discussion of the limitations of these surrogate *in silico* metrics for evaluating sequences generated by protein language models, particularly in the context of downstream experimental validation. Successful expression in a cellular system is a readout of a synthetic protein’s biophysical quality and is often the first step to functional characterization. If, as our results suggest, current *in silico* metrics do not correlate with expression in *E. coli*, their limitations should be taken into account during interpretation, and new metrics should be developed to provide a more holistic readout of model performance. Additionally, our findings point to the need for large-scale experimental screens to measure the expressibility of both natural and AI-generated proteins across expression systems. Measurements from such screens could be used during model training [69, 70, 71] or generation [72, 73] to guide PLMs towards generating proteins that are more likely to express in a system of interest. Importantly, our head-to-head wet-lab comparison of the Dayhoff PLMs revealed computationally and experimentally actionable insights, underscoring the multi-faceted value of experimental validation of generative protein models. We note that successful *in vitro* expression is a necessary but not sufficient condition for protein function. The expression assay measures the biophysical viability of generated sequences and their ability to fold and remain soluble in a cellular environment, but not catalytic activity, binding affinity, or other functional properties. Expression is just a first step in generating proteins – establishing how the function of expressed proteins correlates with model training data composition remains an important next step for future work.

Through the Dayhoff joint sequence-homolog models, our work establishes, for the first time, unified modeling of single sequences, aligned homologs, and unaligned homologs within a single architecture trained with a simple autoregressive objective. While it has been appreciated that sets of homologous protein sequences can provide valuable information for protein design and generation, previous models have treated homologs separately from single sequences and take either aligned homologs [22, 17] or unaligned homologs [25, 24, 23] exclusively. In contrast to these previous approaches, we trained a single model that efficiently generates single sequences and aligned or unaligned homologs. We achieve this via principled architecture design (i.e., a hybrid SSM-transformer) and the unrolling of MSAs into series of sequences that can be provided as a single input, due to the long context length afforded by Dayhoff’s hybrid architecture. Training on unaligned homologs in this way may promote the learning of patterns of evolutionary conservation and change across related sequences – in other words, an implicit, soft evolutionary alignment. At generation time, our unified approach enables guided, conditional generation by simply providing unaligned conditioning sequences as the model’s input context. Leveraging this evolutionary context conditioning, we used a Dayhoff model to design putative Cas9 sequences that exhibit functional domain architectures while being 114-122 residues shorter than the closest natural Cas9, a desirable property for efficient cellular delivery. While these results are promising based on in silico structural prediction and domain annotation, experimental characterization will be required to establish whether the generated sequences exhibit endonuclease function in in vitro and cellular assays We showcase Dayhoff’s flexibility across protein design tasks by demonstrating the ability to do arbitrary sub-sequence inpainting and motif scaffolding with just autoregressive generation. Finally, through head-to-head comparisons across the Dayhoff suite of models, we show that scaling to larger model and dataset sizes improves both the quality of generated proteins and performance in zero-shot prediction of mutation effects on protein function. Across our evaluations, no single dataset composition universally dominated all downstream tasks. Though better-powered experimental analyses are required, models trained on GigaRef exhibited higher expression rates, but Dayhoff-170M-GR resulted in lower ProteinGym scores relative to UniRef-only trained models, likely because metagenomic sequences are more distant from the well-characterized proteins in ProteinGym. Interestingly, Dayhoff-3B-GR-HM-c, a gigaref-containing model, is able to recover ProteinGym performance through a high-quality cooldown phase, maintaining the benefits of diverse training. Structure-based augmentation yielded the highest in vitro expression rate and maintained competitive zero-shot fitness prediction. These tradeoffs suggest that practitioners should select models based on their intended application: Dayhoff-3b-GR-HM-c for zero-shot fitness prediction and family-conditioned generation, and BackboneRef-augmented models for applications prioritizing biophysical viability of de novo designs We release the Dayhoff Atlas – datasets, model weights, and code – under a permissive license to empower the broader community.

Despite its data and modeling contributions, there are notable limitations to Dayhoff that highlight avenues for continued work. First, while Dayhoff is an Atlas resource, neither it nor our evaluations are completely comprehensive. We intend the Dayhoff Atlas to be a first centralized open resource of protein sequence datasets and models. Continued evaluation and experimental validation will be necessary to establish the abilities of the Dayhoff models across downstream biological applications and use cases, such as family-conditioned generation, inpainting of functional motifs, and generation of highly thermostable proteins. Second, there are open opportunities for continued expansion of each of the GigaRef and BackboneRef datasets. For GigaRef, we initially followed assumptions from prior work, which suggested that genomic sequences are more reliable than metagenomic sequences and that singleton metagenomic sequences are unreliable. Likewise, while some singleton sequences may be spurious, “true” singleton sequences reflect the staggering diversity sampled by evolution. Although we omitted singleton sequences from model training, identifying and learning on reliable singletons could lead to increased diversity and quality of generations. For BackboneRef, our results suggest that synthetic data augmentation may effectively transfer distilled structural information into protein sequence space. Notably, our rarefaction analysis indicated that BackboneRef has not yet exhausted all possible novel synthetic backbones; scaling the augmentation dataset and the downstream PLM could additionally lead to improved diversity, stability, and biophysical properties of generated proteins. Last, while we trained models on metagenomic data, structure-based synthetic data, and single sequences together with homologs, we did not combine all data sources into training a single, scaled model to determine whether observed individual gains can be stacked, a clear opportunity for future work.

In sum, we present an open-source and experimentally-validated Atlas that distills protein sequence, evolutionary, and structural information into datasets and trained models for protein sequence design. We envision that the public availability of the Dayhoff Atlas will illuminate new avenues to improve protein design, carrying on the legacy of its namesake.

## Acknowledgments

The authors thank Hannes Schulz and Sean Whitzell for assistance with Microsoft’s compute resources; Philip Rosenfield for his immense effort in managing the project; Hannah Richardson for her efforts in dataset and code release and compliance; and Amelia Zug, Darren Xu, Bruce Wittmann, and Alex Lu for helpful discussions.

## Author contributions

Conceptualization: K.K.Y., S.A., A.J.L., N.T., A.P.A.; Methodology: K.K.Y., S.A., A.J.L., K.K.L., S.C., G.B., C.D.E., C.W., N.T., A.P.A.; Software Programming: K.K.Y., S.A., A.J.L., K.K.L., S.C., G.B., N.T.; Experimental Design: K.K.Y., S.A., A.J.L., N.F., N.T., A.P.A.; Investigation: K.K.Y., S.A., A.J.L., K.K.L., S.C., G.B., C.D.E., C.W., S.L., N.T., A.P.A.; Validation: K.K.Y., S.A., A.J.L., K.K.L., S.C., G.B., S.L., N.T., A.P.A.; Resources Provision: K.K.Y., S.A., N.F., N.T., A.P.A.; Data Curation: K.K.Y., S.A., A.J.L., K.K.L., S.L., N.T., A.P.A.; Visualization: K.K.Y., S.A., A.J.L., S.C., A.P.A.; Writing - Original Draft: K.K.Y., S.A., A.J.L., S.C., N.T., A.P.A.; Writing - Review & Editing: K.K.Y., S.A., A.J.L., K.K.L., S.C., G.B., C.D.E., C.W., S.L., N.F., N.T., A.P.A.; ; Supervision: K.K.Y., S.A., N.T., A.P.A.;

## Resource availability

### Lead contact

Further information and requests for resources should be directed to the corresponding authors, Neil Tenenholtz and Ava P. Amini.

### Materials availability

No new materials were generated in this study.

### Data and code availability

- Code is available at https://github.com/microsoft/dayhoff.
- GigaRef, BackboneRef, and DayhoffRef are available at https://huggingface.co/collections/microsoft/dayhoff-atlas-6866d679465a2685b06ee969.
- Dayhoff models are available on the Azure AI Foundry at https://aka.ms/dayhoff/foundry and on HuggingFace at https://huggingface.co/collections/microsoft/dayhoff-atlas-6866d679465a2685b06ee969.
- Sequences and metrics to reproduce the analyses are available at https://zenodo.org/records/15265289.

## Methods

### Deduplication and clustering

All sequence clustering steps were performed using the linclust function in MMseqs2 [41] in cluster mode 2 and coverage mode 2 at 80% coverage. MMseqs2 selects the longest sequence in each cluster as the representative. Clustering for GigaRef was performed using v15.6f452, and clustering for DayhoffRef was performed using v17.b804f.

### Metagenomic datasets

Metagenomic datasets for GigaRef were downloaded from:

- Mgnify: https://ftp.ebi.ac.uk/pub/databases/metagenomics/peptide_database/2024_04/
- SMAG: https://www.genoscope.cns.fr/tara/localdata/data/SMAGs-v1/
- MetaEuk: https://wwwuser.gwdguser.de/~compbiol/metaeuk/
- MGV: https://portal.nersc.gov/MGV/
- GPD: https://ftp.ebi.ac.uk/pub/databases/metagenomics/genome_sets/gut_phage_database/
- SRC: https://github.com/soedinglab/plass
- MERC: https://github.com/soedinglab/plass
- TOPAZ: https://osf.io/gm564/

A multi-step clustering process was performed to combine, deduplicate, and recluster these datasets with UniRef100. MERC and SRC were combined and preclustered at 70% identity, and MGnify was preclustered at 70% identity. The representatives resulting from each of these preclustering steps were then combined with sequences from SMAG, MetaEuk, MGV, GPD, TOPAZ, and UniRef100, yielding a total of 3.94 billion sequences. These were clustered at 90% sequence identity followed by clustering of the resulting representatives at 50% identity to produce the GigaRef dataset.

### Structural plausibility

We characterized the structural plausibility of sequences using the ESM-Fold [42] pLDDT with default parameters and the self-consistency perplexity (scPerplexity). We computed scPerplexity by predicting a structure for each sequence with ESMFold, redesigning each predicted structure with ProteinMPNN [43] at *T* =1.0, and computing the perplexity against the original sequence.

### Fréchet protein distance (FPD)

To estimate the divergence between distributions of protein sequences, we sampled 1024 sequences from each distribution, embedded them with the ProtBert-BFD [35] model, and computed the earth mover’s distance

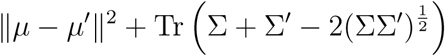

where *µ* and *µ*^′^ are average vectors in the ProtBert embedding space and Σ and Σ^′^ are covariance matrices for the two sets of embeddings. Tr refers to the trace of this matrix. We report FPDs to the UniRef50 and GigaRef test sets. For the UniRef50 ↔ UniRef50 and GigaRef-clusters ↔ GigaRef-clusters comparisons, we computed the FPD between the train and test sets. For the UniRef50 ↔ GigaRef-clusters comparison, we computed the FPD between the test sets.

### ProtNote MMD (PNMMD)

We used ProtNote [44] to annotate samples of 10M sequences for each dataset. ProtNote outputs a probability for each of the 32,102 functions in the Gene Ontology [74], indicating the likelihood that a protein sequence is annotated with that function. For numerical stability we used ProtNote’s logits scaled by 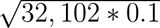. Then, we computed the kernel Maximum Mean Discrepancy (MMD) [75] with Radial Basis Function kernels, selecting the kernel’s bandwidth hyperparameter using the median heuristic [76]. This heuristic sets the bandwidth to the median pairwise distance between all points in the combined sample (i.e. the union of both samples). Computing the MMD or median heuristic involves calculating all pairwise distances between points in a sample, which is memory-intensive for samples of 10M points. We thus used Monte Carlo sampling to subsample batches without replacement of 1K points for 10K trials. The estimates converge around 2K trials for the MMD, and 200 trials for the median heuristic (Fig. S2-S3). Note that for the baseline comparisons GR vs. GR and UR50 vs. UR50, we compare the first half of the dataset with the second half, resulting in 5K trials instead of 10K. There is no need to generate an additional 10M samples for these datasets, as the results stabilize at around 2K trials.

### Pfam annotation with HMMER

Pfam annotations were computed using Pfam-A profiles from Pfam v37.2 using version 3.3 of HMMER [77] hmmscan with default parameters to compute Pfam [78] annotations for 8M randomly-selected sequences from UniRef50, 4M randomly-selected GigaRef cluster representatives (no singletons), 4M randomly-selected GigaRef singletons, and all sequences in BRu, BRq, BRn, and DayhoffRef. For **Fig. 2G** and **Fig. 4I-J**, we downsampled the domains to produce more comprehensible plots. The predicted domains were sorted by the x-axis count, and every 10th domain was plotted.

### Synthetic backbone generation for BackboneRef

We used RFdiffusion (checkpoint “Base_ckpt”) [45] for all backbone generations. Structures were sampled according to the length distribution of UniRef50 [2] to recapitulate the lengths of natural proteins but with a minimum length of 40 and maximum length of 512 for computational efficiency.

### Secondary structure analysis

For all secondary structure analyses, we predicted 3-class (loops, helices, and strands) type assignment using DSSP [79]. We performed this analysis directly on the backbones, rather than on the sequence-designed and folded structures. For visualization of percentage strandedness and helicity for each backbone, we averaged the count of the given annotation class over the structure.

### Self-consistency analysis for BackboneRef backbones

For each backbone, we used Protein-MPNN [43] at temperature 0.1 to design 10 sequences per backbone. We then used OmegaFold [80] to predict a structure for each sequence, and then computed the scRMSD between the C*_α_* positions of the original backbone and the predicted structures.

We also separately conducted fixed-backbone sequence design with temperatures 0.2, 0.3, 0.4, 0.6, 0.8, and 1.0, designing ten sequences per backbone per temperature prior to structure prediction using OmegaFold. scRMSD was then computed between each backbone’s predicted structures at the the various temperatures and the original backbone.

### BackboneRef Foldseek novelty search

Rather than using Foldseek’s [47] easy-search directly on synthetic backbones in C*_α_* mode, we used the lowest scRMSD all-atom structure (generated by backbone-conditioned sequence design with ProteinMPNN and then predicting the structure with OmegaFold, as above) for each generated backbone. We then used Foldseek to query these structures against the AlphaFold/UniProt database, ‘AFDB/UniProt’ [48], which comprises AlphaFold2 structure predictions for every protein in UniProt. We report the maximum TM-score returned per query; this represents the closest structural match to the query structure.

### BackboneRef Foldseek clustering

For clustering analyses of BackboneRef, we again selected the lowest scRMSD all-atom structure and its corresponding sequence for each generated backbone. We then clustered these structures along with all structures from the PDB (209,850 structures) using Foldseek’s [47] easy-cluster with default settings. For PDB files with multiple chains, we selected the first one.

### BackboneRef data preparation

We first generated an initial set of 300 sequences per backbone at temperature 0.2, producing a parent dataset with 72,243,300 sequences. To create the BRu dataset, we randomly selected 42 sequences per backbone from this parent dataset, yielding 10,114,860 sequences. We then removed exact duplicates, and as this yielded an excess of 10M sequences, we then subsampled the remaining sequences randomly to produce a dataset of 10M sequences. To create the BRq dataset, we removed any backbones with average scRMSD score greater than 2Å, leaving 127,633 backbones. We randomly selected 80 sequences per backbone from the parent dataset, and again removed exact duplicates before random subsampling to produce a dataset of 10M sequences. To create the BRn dataset, we removed any backbones with maximum TM-score larger than 0.5 to any structure in the ‘AFDB/UniProt’ database [48], leaving 138,044 backbones. We randomly sampled 74 sequences per backbone from the parent dataset and again removed exact duplicates before random subsampling to produce another dataset of 10M sequences. We constructed combined natural-synthetic datasets by taking the union of the UniRef50 training split (63,662,039 sequences) and each of the synthetic datasets described above. This produced training datasets of approximately 73M sequences each.

### BackboneRef rarefaction analysis

To quantify whether BackboneRef samples were saturating in structural diversity, we conducted a rarefaction analysis (Fig. S5). We randomly sub-sampled the combined dataset of BackboneRef synthetic backbones and PDB structures at ten different frequencies (0.05, 0.1, 0.2, 0.3, 0.4, 0.5, 0.6, 0.7, 0.8, and 0.9), producing ten sub-datasets. We conducted this procedure once with resampling both PDB and BackboneRef structures and once resampling only BackboneRef structures. Due to computational resource limitations, we did not re-run Foldseek on each sub-dataset. Instead, we used the cluster assignments from the combined PDB and BackboneRef Foldseek clustering experiment. For each resampling frequency, we computed the number of distinct structural clusters, derived from the original Foldseek clustering, present in each subsample.

### Model training and architecture

All Dayhoff models use either a 170m- or 3b-parameter model architecture, based on Jamba [54] and trained with an autoregressive objective. Every eighth layer is a transformer module, with the remainder being Mamba modules [49]. Every other Mamba module uses mixture of experts with 16 experts instead of a multilayer perceptron layer. All Mamba layers have state dimension 16, convolution dimension 4, and expansion dimension 2. For all models, we use the Adam optimizer with betas (0.9, 0.999).

The 170m-parameter models have 24 layers with model dimension 256, intermediate layer widths of 1024 inside each transformer block, 16 attention heads per transformer, and 8 key-value heads per transformer, resulting in 170,298,096 total parameters.

170m-parameter models were trained on either 8 NVIDIA A100 or 8 NVIDIA H100 GPUs with adaptive batch sizes such that each batch had at most 360,000 tokens per GPU using distributed-data-parallel. Each sample was randomly selected to be N-to-C or C-to-N. We used an inverse square root scheduler with linear warmup of 10,000 steps, max learning rate of 4e-4, 76,000 total steps, and MoE auxiliary loss weight 1.0.

The 3b-parameter models have 32 layers with model dimension 1280, intermediate layer widths of 2048 inside each transformer block, 32 attention heads per transformer, and 8 key-value heads per transformer, resulting in 2,981,431,616 total parameters.

Dayhoff-3b-UR90 was trained on 176 NVIDIA H100s with PyTorch fully-sharded data parallel leveraging a hybrid (within-node) ZeRO-2 strategy [81, 82, 83] with adaptive batch sizes such that each batch had at most 90,000 tokens per GPU. We used an inverse square root scheduler with linear warmup of 11,000 steps, max learning rate of 2.75e-4, 43,300 total steps, and MoE auxiliary loss weight 0.001.

Dayhoff-3b-GR-HM was trained on 144 NVIDIA H100s with PyTorch fully-sharded data parallel using a ZeRO-3 strategy [83] with adaptive batch sizes such that each batch had at most 120,000 tokens per GPU. Half of the GPUs received sequences from GigaRef, while the other half received homologs from OpenProteinSet. GigaRef sequences were sampled uniformly from clusters, and each sample was randomly selected to be N-to-C or C-to-N. For homologs, each sample was randomly selected to be N-to-C or C-to-N, to be aligned or unaligned, and to be in query mode or no-query mode. Whether the homologs are aligned or unaligned is denoted by surrounding the sequence of sequences with start_align and stop_align or start_unalign and stop_unalign tokens. In query mode, the query sequence from the MSA is denoted by surrounding it with the single-sequence start and stop tokens and placed either before or after the other homologs with equal probability. The hybrid Mamba-Transformer architecture uses rotary positional embeddings that support a maximum context length limit of 262,144 tokens. Due to computational constraints, single sequences were truncated to 2,048 residues during training. Aligned MSAs were subsampled to a maximum of 64 homologs of up to 512 residues each ( 32,768 tokens total). For unaligned homologs, aligned homologs of up to 512 residues each were sampled, then insertions were re-inserted, resulting in the possibility of unaligned homolog sequences of longer than 512 and overall context lengths exceeding 32k tokens. We used an inverse square root scheduler with linear warmup of 11,000 steps, max learning rate of 2.75e-4, 52,000 total steps, MoE auxiliary loss weight 0.001, and grad norms clipped to 1.0.

Beginning with the weights of Dayhoff-3b-GR-HM, we then performed 2 cooldown cycles with UniRef90 and homologs from OpenProteinSet to train Dayhoff-3b-GR-HM-c. In the first cycle, we warmed up the learning rate linearly from 0 to 2.75e-4 over 2000 steps, then performed a cosine decay to 0 over the next 10,000 steps. Unfortunately, this model diverged sometime after 8,000 decay steps, so we began the second cycle from the weights after 8,000 decay steps. In the second cycle, we warmed up the learning rate linearly from 0 to 2.75e-4 over 6000 steps, performed a cosine decay to 2.75e-5 over the next 18,000 steps, and then trained for 1000 more steps at a learning rate of 2.75e-5.

### DayhoffRef

DayhoffRef sequences were sampled in both the N-to-C and C-to-N directions from Dayhoff-3b-GR-HM, Dayhoff-3b-GR-HM-C, and Dayhoff-170m-UR50-BRn at temperatures 0.9 and 1.0 on 8x AMD MI300X GPUs for each combination of model, direction, and temperature. Clustering to 90% identity was performed with MMseqs2 linclust using default parameters, and then the cluster representatives were clustered to 50% identity using default parameters.

### Computation of test set perplexity

A model’s perplexity on held-out sequences from the test set was computed as:

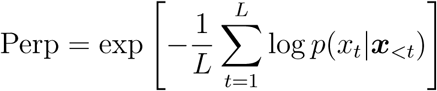

We computed perplexity under the same conditions as training: single sequences were randomly selected to be presented N-to-C or C-to-N; homologuous sequences were randomly selected to be aligned or unaligned and to be in query mode or no-query mode; and in query mode, the query was randomly placed at the beginning or end.

### Unconditional generations

We generated 1024 sequences in the N-to-C and C-to-N direction for all Dayhoff models at each of the temperatures 0.7, 0.8, 0.9, 1.0, 1.1, and 1.2, with a maximum sequence length of 2048. We characterized the quality of generated sequences using the ESMFold [42] pLDDT and the self-consistency perplexity (scPerplexity). We computed scPerplexity by folding each sequence with ESMFold, redesigning each predicted structure with ProteinMPNN [43] (inverse folding *T* =1.0), and computing the perplexity against the original sequence. These results are included in Tables S20-S29 and summarized in Table S19.

### Zero-shot scoring of mutation effects

We used the datasets from ProteinGym [58] for this evaluation. For each variant, we computed the validation likelihood with teacher forcing. We then used the ProteinGym software [58] to compute the Spearman correlation coefficient for each dataset and to compute a weighted average over the datasets. When scoring without homologs, we averaged the likelihoods in the N-to-C and C-to-N directions. When scoring with homologs, we randomly resampled homologs from the parent MSA to 64 sequences 5 times and averaged the likelihoods.

### Motif scaffolding

The Dayhoff models were evaluated on two motif scaffolding benchmark sets: RFDiffusion [45] and MotifBench [63]. 8 scaffolding problems appear in both sets: 04_5WN9, 06_6E6R, 07_6E6R, 12_4JHW, 13_4JHW, 14_5IUS, 22_1BCF, and 28_5YUI, using the MotifBench naming. For each scaffolding problem, we generated 100 designs at *T* = 1.0 and followed the approach used in EvoDiff [17] to determine success rates. For scaffolding problems that contain multiple motif segments, the distance between segments was kept fixed to the distance in between segments in the given PDB sequence. Each scaffolding problem specifies a length constraint describing how many residues the final scaffold designs should contain. In cases where keeping a fixed distance between motif segments made the scaffold longer than the length constraint, we randomly reduced the length of one inter-motif segment at a time until the scaffold met the length restriction. Sequences were evaluated using blast p against the BLAST non-redundant protein database downloaded on June 17th 2025 [68]. Designs were considered successful if the RMSD of the motif between the original and the designed scaffolds (the motifRMSD) was ≤ 1.0Å, the ESMFold pLDDT of the entire sequence was ≥ 0.7, and the sequence was not an exact match to any existing protein in the BLAST non-redundant database. The total number of successes and number of problems solved for each model is given in Tables S32-S35.

To compare performance across models, we leveraged the MotifBench score [63], defined as:

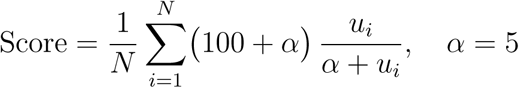

where *N* is the number of motif scaffolding problems in each benchmark and *u_i_* is the number of unique solutions for problem *i*. To determine the number of unique solutions for each problem, we clustered the structures of the designed scaffolds. In most cases, we used the Foldseek’s [47] easy-cluster method, as used in MotifBench and which clusters structures using 3Di+AA tokens that capture both structure and amino acid information, and counted each Foldseek cluster as an unique solution. In the RFdiffusion benchmark there are cases where the designed scaffold is less than 50 residues, which is too short to be handled by Foldseek. In these cases, we measured pairwise TM-scores across all folded designs and applied single-linkage hierarchical clustering using a cutoff of TM-score *>* 0.6 to determine unique clusters, which we used to count unique solutions. Briefly, any two designs with a TM-score *>* 0.6 were joined, and clusters were defined as the union over all disjoint sets (implemented via a union–find data structure). The resulting clusters contained non-overlapping sets, where each cluster’s first member in sorted filename order was taken as its representative solution.

### Evolution-guided generation

For these evaluations, we generated with temperature 1.0 and min-p sampling [84] (minP=0.05) for 226 randomly-chosen MSAs from the test set where the query sequence begins with methionine, is between 64 and 451 amino acids in length, and has at least 4 homologs. In order to generate on a single A6000 GPU, we conditioned Dayhoff on up to 56 randomly-chosen homologs in no-query mode. For baselines, we used those same sequences to train and sample with CCMgen using default parameters, to regenerate the query using EvoDiff-MSA, and to condition generation with Prot-xLSTM at *T* = 1.0, top-k of 20, and top-p of 0.9. We characterized the sequence novelty of each generation by aligning the generation with each sequence in the original MSA using the BioPython [85] pairwise aligner with default parameters and then finding the most similar natural sequence. We also predicted structures and computed pLDDT for each generation and original (natural) query using ESMFold. These structures were used to compute the TM-score to the natural structure. scPerplexity was computed using ProteinMPNN at *T* = 1.0.

### Generation of short Cas9s

We used MMseqs2 linclust to cluster natural Cas9s from Makarova et al. [66] at 90% identity, yielding 413 clusters. We then removed sequences longer than 1050 residues, resulting in 45 clusters containing a total of 93 sequences. Conditioning sequences were chosen by sampling one sequence from each of 32 randomly-sampled natural clusters, ensuring that at least one conditioning sequence had fewer than 1000 residues. These were then used in-context to generate a sequence from Dayhoff-GR-HM-c at *T* = 1.0 and minP=0 in the N-to-C direction in no-query mode. We rejected samples that were not between 801 and 960 residues. We generated one accepted shortened sequence for each set of conditioning sequences. If a set of conditioning sequences did not generate an accepted sample after 5 tries, we chose a new\ set of conditioning sequences and generated a new sequence. This process was repeated until we had 100 accepted samples. We predicted a structure for each sequence and computed its pLDDT using ESMFold. We found the closest natural Cas9 to each generated sequence using the BioPython pairwise aligner with the BLOSUM62 substitution matrix, a gap open penalty of 10, and a gap extend penalty of 0.5, then used this alignment to determine whether there was a large C-terminal truncation. For visualization, we realigned each selected generated Cas9 with the closest natural Cas9 with Emboss Needle [86] using default parameters and predicted domains for each natural and generated Cas9 using the InterPro web interface [67].

### Nomination and in vitro expression of Dayhoff-3b-UR90 generations

We generated sequences from Dayhoff-3b-UR90 in each of the N-to-C and C-to-N directions at temperatures 0.7, 0.8, 0.9, and 1.0. We eliminated sequences with OmegaFold pLDDT *<* 0.8 and then manually nominated sequences with low scPerplexity, globular predicted structures, no homology hits to membrane proteins, strong homology hits to bacterial proteins, and no cysteine residues. The final 25 sequences included 13 generated at *T* = 0.7 and 4 sequences at each of *T* = 0.8, 0.9, and 1.0. Sequences were codon-optimized for expression in *E. coli*, appended with a C-terminal twin-strepII tag for purification, and inserted into a pET-29b(+) vector with an inducible T7/lacI promoter for expression. A GFP construct inserted into the same vector was used as a positive control for all downstream expression, purification, and characterization. The resulting plasmids were transformed into the *E. coli* strains BLR(DE3) and BL21-AI (a derivative of BL21 designed for strong and non-leaky expression). Following verification by sequencing, overnight cultures were used to inoculate growth in 4 replicate 1 mL cultures in LB media at 37^◦^ C for 5 hours or 3 hours for BLR(DE3) and BL21-AI, respectively. Expression was induced using 1mM IPTG or 1mM IPTG + 0.2% L-arabinose for BLR(DE3) and BL21-AI, respectively. Post-induction, BLR(DE3) cells were grown for 15 hours at 16^◦^C, and BL21-AI cells were grown for 3 hours at 37^◦^C. Cells were pelleted via centrifugation and then chemically lysed using BugBuster (Millipore Sigma) supplemented with cOmplete™, EDTA-free Protease Inhibitor Cocktail (Roche).

Proteins in the soluble fraction (supernatant) were purified on a Hamilton Microlab STAR Liquid Handler (Hamilton) using IMCStips 100*µ*L Streptactin 4Flow XT tips (IMCS). Samples were buffer exchanged using Zeba desalting plates (Thermo Fisher Scientific) in 10 mM sodium phosphate buffer. Protein purity and abundance in the soluble fraction was assessed with SDS-PAGE and with the Pierce 660nm Protein Assay Reagent (Thermo Fisher Scientific) with colorimetric readout. A protein was considered successfully expressed if any soluble protein was detected at the correct molecular weight by SDS-PAGE. *In vitro* expression and purification experiments were conducted by Ginkgo Bioworks. Sequences and metrics are provided in Zenodo.

### Structural characterization by UV circular dichroism

Circular dichroism (CD) was used to characterize secondary structures in manually-nominated generated proteins from Dayhoff-3b-UR90. All CD data collection was performed with the Jasco high-throughput circular dichroism (HTCD) measurement on the Jasco J-1500 CD spectrophotometer (Jasco), with secondary structure characteristics determined via measurement of the far-UV CD spectra. Measurements were acquired in the far-UV spectra ranging from 200 to 250 nm; all reported measurements were acquired within the linear range of the instrument and aggregated over 5 replicate accumulations. A buffer blank and GFP were used as negative and positive controls, respectively. CD experiments were conducted by Ginkgo Bioworks.

### Head-to-head comparisons of in vitro expression

Starting from 200 generations each from each of Dayhoff-170m-UR90, Dayhoff-170m-GR, Dayhoff-170m-UR50-BRn, Dayhoff-3b-GR-HM, and Dayhoff-3b-GR-HM-c, we selected a total of 144 sequences for experimental characterization. Sequences were filtered out from the original set if they were *<* 60 amino acids or *>* 1500 amino acids. We also filtered out sequences predicted to have a signal peptide by TargetP [87], mitochondrial transit peptide by TargetP, or a transmembrane helix by DeepTMHMM [88]. Lastly, sequences were also filtered out if they exhibited a grand average of hydropathicity (GRAVY) score *>* 0, which indicates a highly hydrophobic protein, an instability index *>* 70, or an isoelectric point *>* ± 2 units of the purification buffer pH of 7.5; these metrics were calculated using ProtParam [89]. After filtering, there were 66 Dayhoff-170m-UR90, 59 Dayhoff-170m-GR, 47 Dayhoff-170m-BBRn, 50 Dayhoff-3b-GR-HM, and 45 Dayhoff-3b-GR-HM-c sequences remaining. We randomly selected 29 from each of the first four sets and 28 from Dayhoff-3b-GR-HM-c to yield a total of 144 sequences for *in vitro* testing.

Sequences were codon-optimized for expression in *E. coli*, appended with a C-terminal twin-strepII tag for purification, and inserted into a pET-29b(+) vector with an inducible T7/lacI promoter for expression. A GFP construct inserted into the same vector was used as a positive control for all downstream expression, purification, and characterization. The resulting plasmids were transformed into BLR(DE3) and BL21-AI. Following verification by sequencing, overnight cultures were grown up and used to inoculate growth in 2 replicate 2 mL cultures in LB media at 37^◦^ C for 5 hours or 3 hours for BLR(DE3) and BL21-AI, respectively. Expression was induced using 1mM IPTG or 1mM IPTG + 0.2% L-arabinose for BLR(DE3) and BL21-AI, respectively. Post-induction, BLR(DE3) and BL21-AI cells were grown for 16-18 hours at 16^◦^C. Cells were pelleted via centrifugation and then chemically lysed using BugBuster (Millipore Sigma) supplemented with cOmplete™, EDTA-free Protease Inhibitor Cocktail (Roche).

From the soluble fraction (supernatant), proteins were purified on a Hamilton Microlab STAR Liquid Handler (Hamilton) using IMCStips 100*µ*L Streptactin 4Flow XT tips (IMCS). Samples were buffer exchanged using Zeba desalting plates (Thermo Fisher Scientific) in 10 mM sodium phosphate buffer. Protein abundance in the soluble fraction was assessed using SDS-PAGE and the Lunatic UV/Vis spectrophometer (Unchained Labs). A protein was considered successfully expressed if any soluble protein was detected at the correct molecular weight by SDS-PAGE. *In vitro* expression and purification experiments were conducted by Ginkgo Bioworks. Sequences and metrics are provided in Zenodo.

## Supplemental Figures and Tables

**Table S1:**
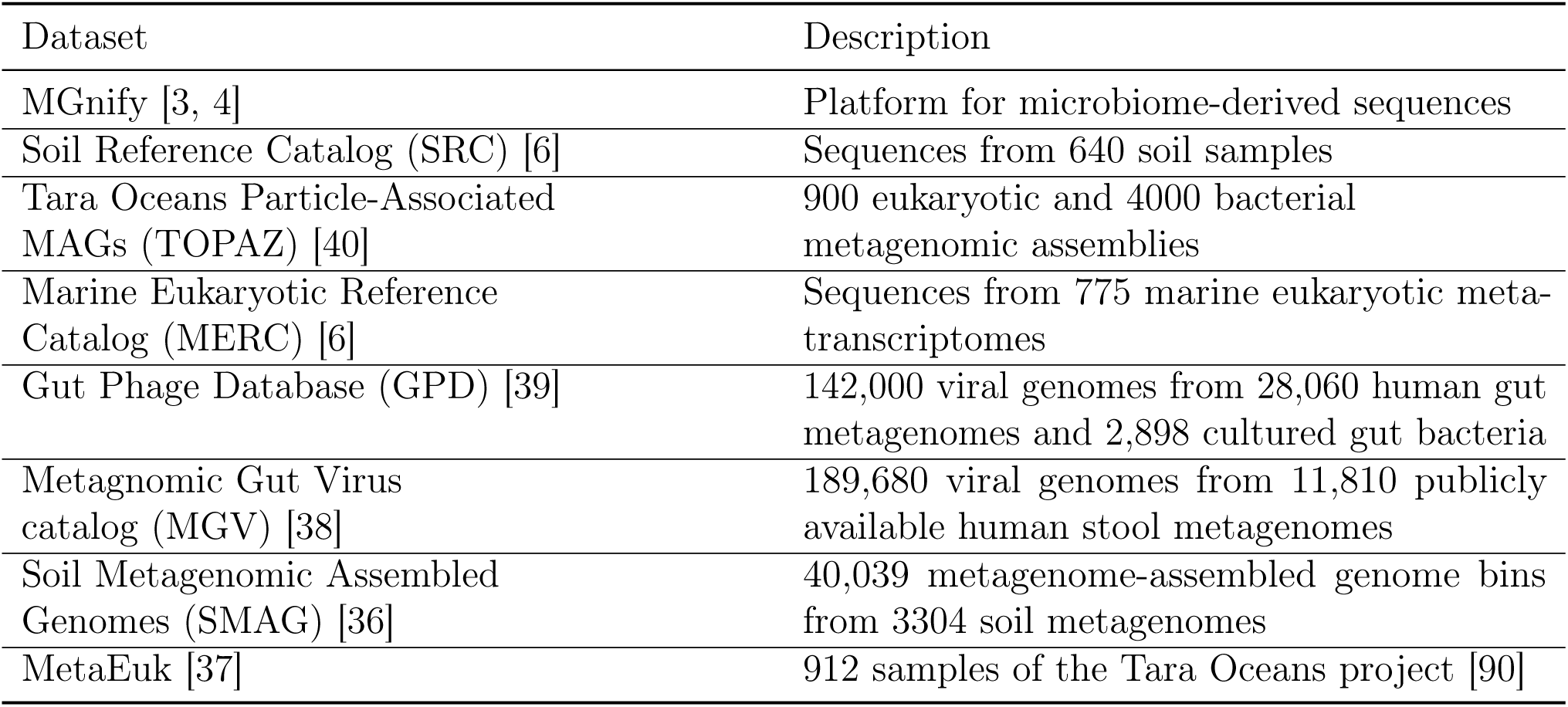
Metagenomics datasets used to construct GigaRef.

| Dataset | Description |
| --- | --- |
| MGnify [3, 4] | Platform for microbiome-derived sequences |
| Soil Reference Catalog (SRC) [6] | Sequences from 640 soil samples |
| Tara Oceans Particle-Associated MAGs (TOPAZ) [40] | 900 eukaryotic and 4000 bacterial metagenomic assemblies |
| Marine Eukaryotic Reference Catalog (MERC) [6] | Sequences from 775 marine eukaryotic meta-transcriptomes |
| Gut Phage Database (GPD) [39] | 142,000 viral genomes from 28,060 human gut metagenomes and 2,898 cultured gut bacteria |
| Metagenomic Gut Virus catalog (MGV) [38] | 189,680 viral genomes from 11,810 publicly available human stool metagenomes |
| Soil Metagenomic Assembled Genomes (SMAG) [36] | 40,039 metagenome-assembled genome bins from 3304 soil metagenomes |
| MetaEuk [37] | 912 samples of the Tara Oceans project [90] |

**Table S2:** Summary metrics for the GigaRef dataset.

| Dataset | $n$ before deduplication | $n$ in clusters | $n$ in singletons | Fraction singleton |
| --- | --- | --- | --- | --- |
| MGnify | 2,973,257,435 | 1,331,641,136 | 33,661,828 | 0.02 |
| SRC | 2,022,891,389 | 326,171,842 | 1,267,562,989 | 0.80 |
| UniRef100 | 408,368,587 | 173,487,339 | 0 | 0.0 |
| MERC | 292,137,902 | 46,221,844 | 144,579,297 | 0.76 |
| MGV | 11,837,198 | 863,949 | 0 | 0.0 |
| SMAG | 10,207,435 | 2,267,280 | 5,755,133 | 0.72 |
| TOPAZ | 8,090,866 | 4,539,508 | 0 | 0.0 |
| GPD | 7,581,807 | 561,073 | 219,211 | 0.28 |
| MetaEuk | 6,158,526 | 1,206,140 | 1,991,621 | 0.62 |
| **Total** | 5,740,531,145 | 1,886,960,111 | 1,453,770,079 | 0.44 |

**Table S3:** Distribution shifts between Dayhoff datasets as measured by Frechet ProtBert Distance (FPD)

| Datasets | FPD |
| --- | --- |
| UniRef50 ↔ UniRef50 | 0.040 |
| GigaRef ↔ GigaRef | 0.040 |
| GigaRef ↔ GigaRef-s | 0.14 |
| UniRef50 ↔ GigaRef | 0.32 |
| UniRef50 ↔ GigaRef-s | 0.46 |
| UniRef50 ↔ BRu | 4.78 |
| UniRef50 ↔ BRn | 4.34 |
| UniRef50 ↔ BRq | 5.01 |
| GigaRef ↔ BRu | 5.10 |
| GigaRef ↔ BRn | 4.64 |
| GigaRef ↔ BRq | 5.31 |

**Figure S1:**
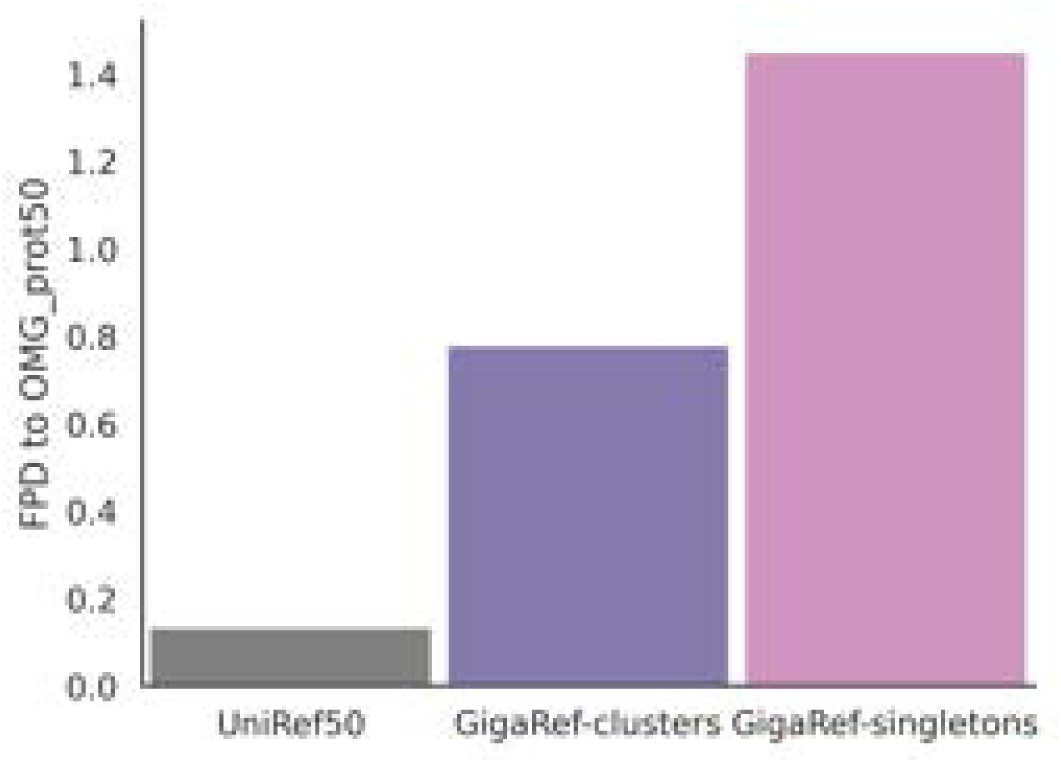
Distributional distances, as measured by FPD, between 10,000 randomly sampled OMG_prot50 sequences relative to each of the UniRef50 and GigaRef-clusters test sets, as well as sequences sampled from GigaRef-singletons.

**Table S4:**
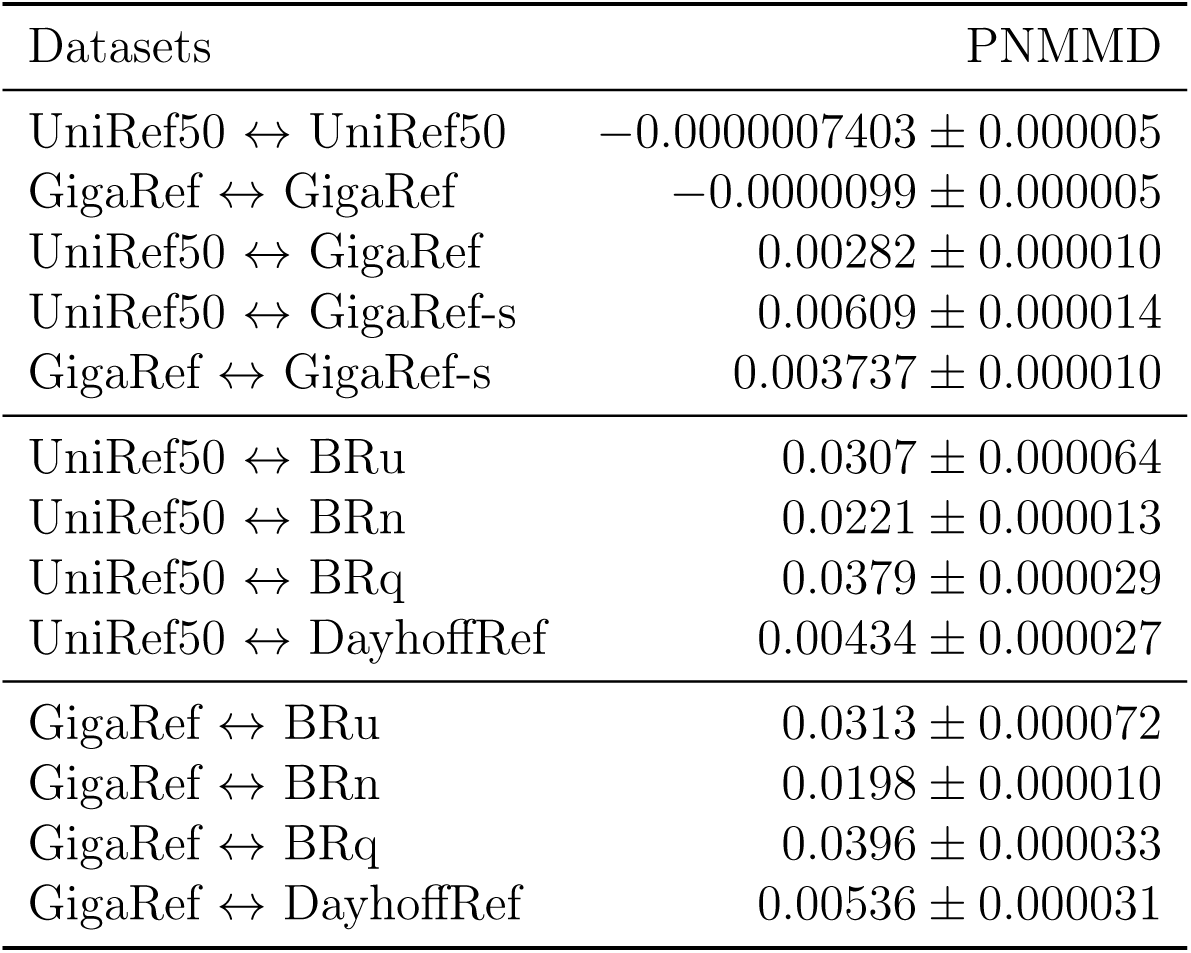
Distribution shifts between Dayhoff datasets as measured by ProtNote maximum mean discrepancy (PNMMD). Lower values indicate more similar datasets. Uncertainties are standard errors over 10k paired sets of 1000 proteins each. Note that the unbiased MMD estimator can yield negative values, especially when the true MMD is close to zero, reflecting the estimator’s ability to fluctuate around the true value.

**Figure S2:**
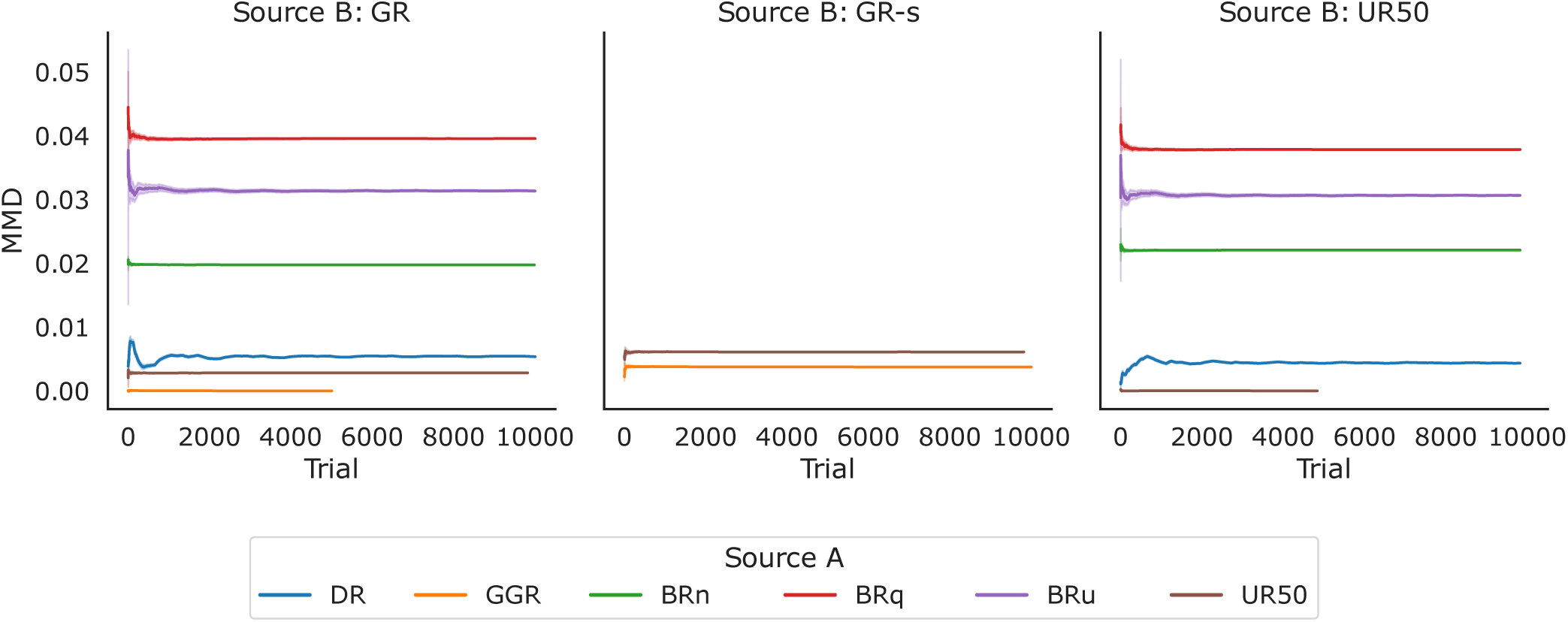
ProtNote Maximum Mean Discrepancy (PMMD) between each sample source and a fixed reference dataset shown in each panel. Each line shows the running mean of the PMMD metric across trials, while the shaded region indicates the running standard error. The PMMD is computed between the sample source (as indicated in the legend) and the reference dataset named in the panel title. Lower values indicate greater similarity. PMMD values are computed over 10,000 trials for all cases except GR vs. GR and UR50 vs. UR50, which have 5,000 trials.

**Figure S3:**
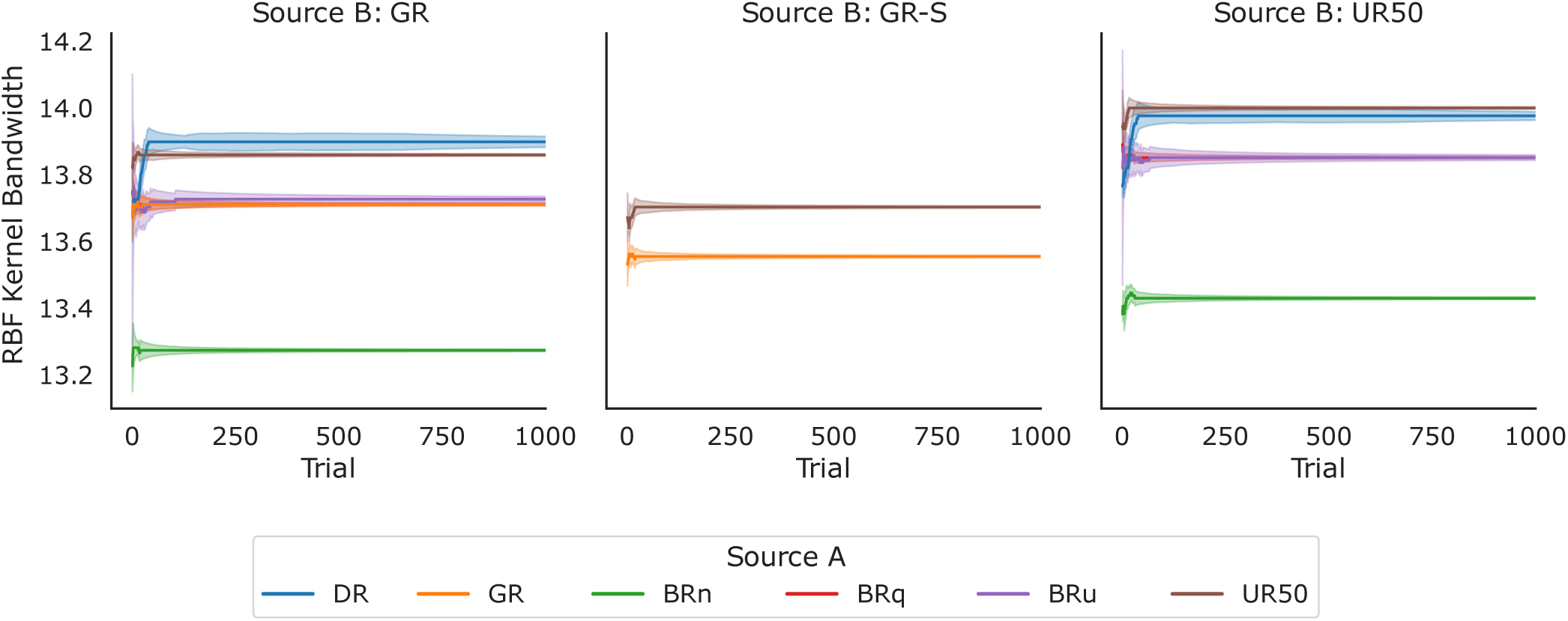
Estimated RBF kernel bandwidth hyperparameter for ProtNote Maximum Mean Discrepancy (PMMD). The bandwidth is estimated using the median heuristic, which sets it to the median pairwise distance between all points in the combined sample – i.e., the union of a sample source (as indicated in the legend) and the reference dataset (named in each panel title). Each line shows the running mean of this median distance across trials for the corresponding combined sample, and the shaded region indicates the running standard error.

**Table S5:** Metrics on Pfam annotations for Dayhoff datasets, with Pearson’s *r* correlation coefficients.

| Dataset | Domains / sequence | $r$ to UniRef50 |
| --- | --- | --- |
| UniRef50 | 1.58 | 1.00 |
| GigaRef | 1.11 | 0.86 |
| GigaRef-singletons | 1.00 | 0.87 |
| BackboneRef-u | 1.24 | 0.11 |
| BackboneRef-q | 1.68 | 0.10 |
| BackboneRef-n | 0.84 | 0.16 |
| DayhoffRef | 0.77 | 0.96 |

**Figure S4:**
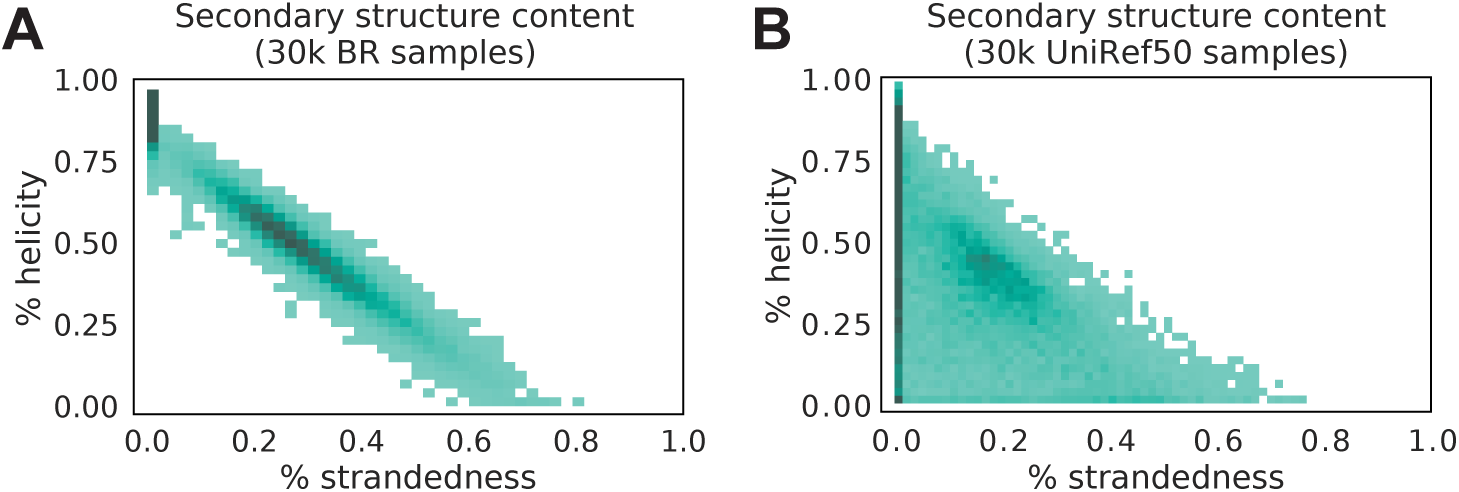
Structural diffusion models preferentially generate helical structures. (**A**) Histograms of percentage of residues called as stranded (x-axis) or helical (y-axis) for (**A**) *n* = 30, 000 randomly-selected synthetic backbones from BackboneRef and (**B**) *n* = 30, 000 randomly-selected structures for sequences in UniRef50, with structures downloaded from AlphaFold-DB.

**Table S6:**
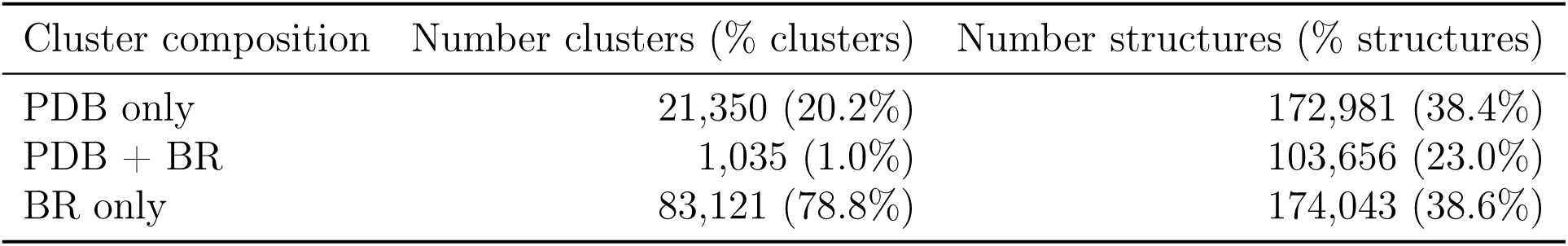
Numbers of clusters and structures for inferred clusters in the BackboneRef novelty analysis.

| Cluster composition | Number clusters (% clusters) | Number structures (% structures) |
| --- | --- | --- |
| PDB only | 21,350 (20.2%) | 172,981 (38.4%) |
| PDB + BR | 1,035 (1.0%) | 103,656 (23.0%) |
| BR only | 83,121 (78.8%) | 174,043 (38.6%) |

**Figure S5:**
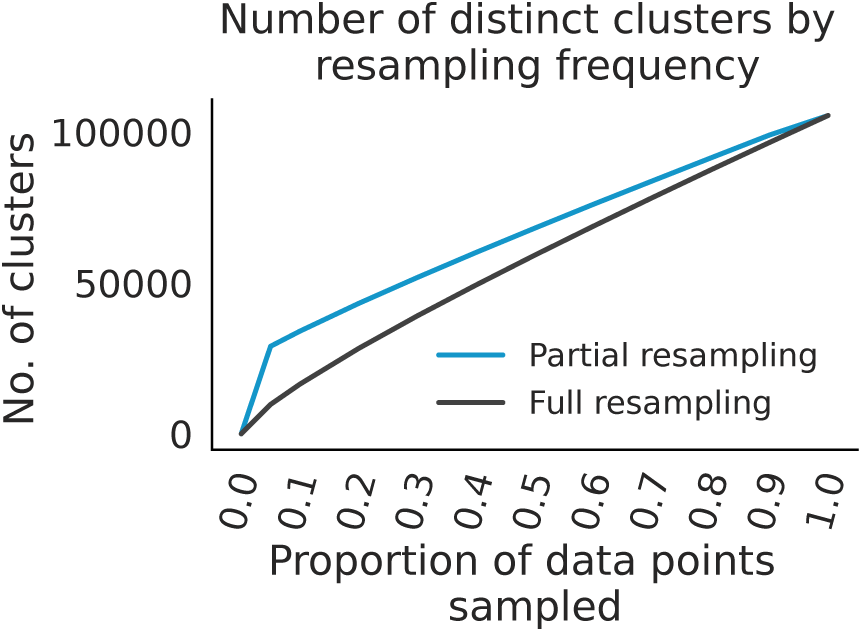
Rarefaction analysis of inferred structural clusters from Foldseek. Foldseek was used to cluster a dataset comprised of structures downloaded from the PDB concatenated to the lowest scRMSD predicted structures from BackboneRef. Structures were resampled at varying frequencies (either across the whole dataset, grey, or only resampling BackboneRef samples, blue), and the number of distinct Foldseek clusters was determined. Note that we do not recompute clusters at different frequencies and instead resample structures and then annotate from the full set of inferred Foldseek clusters.

**Figure S6:**
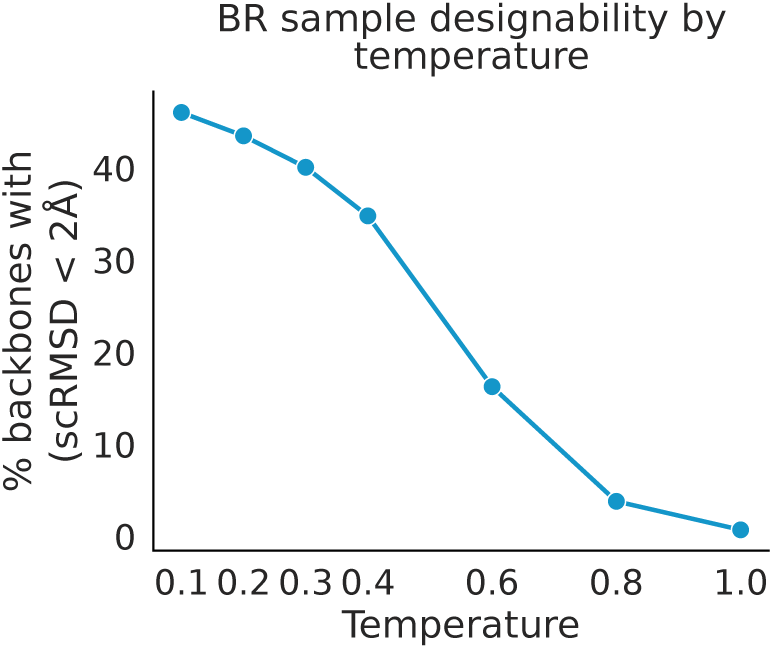
Percentage of designable samples from BackboneRef by temperature, with the percentage of designable BackboneRef samples computed as those with average scRMSD *<* 2Å over 10 predicted structures at a given temperature.

**Figure S7:**
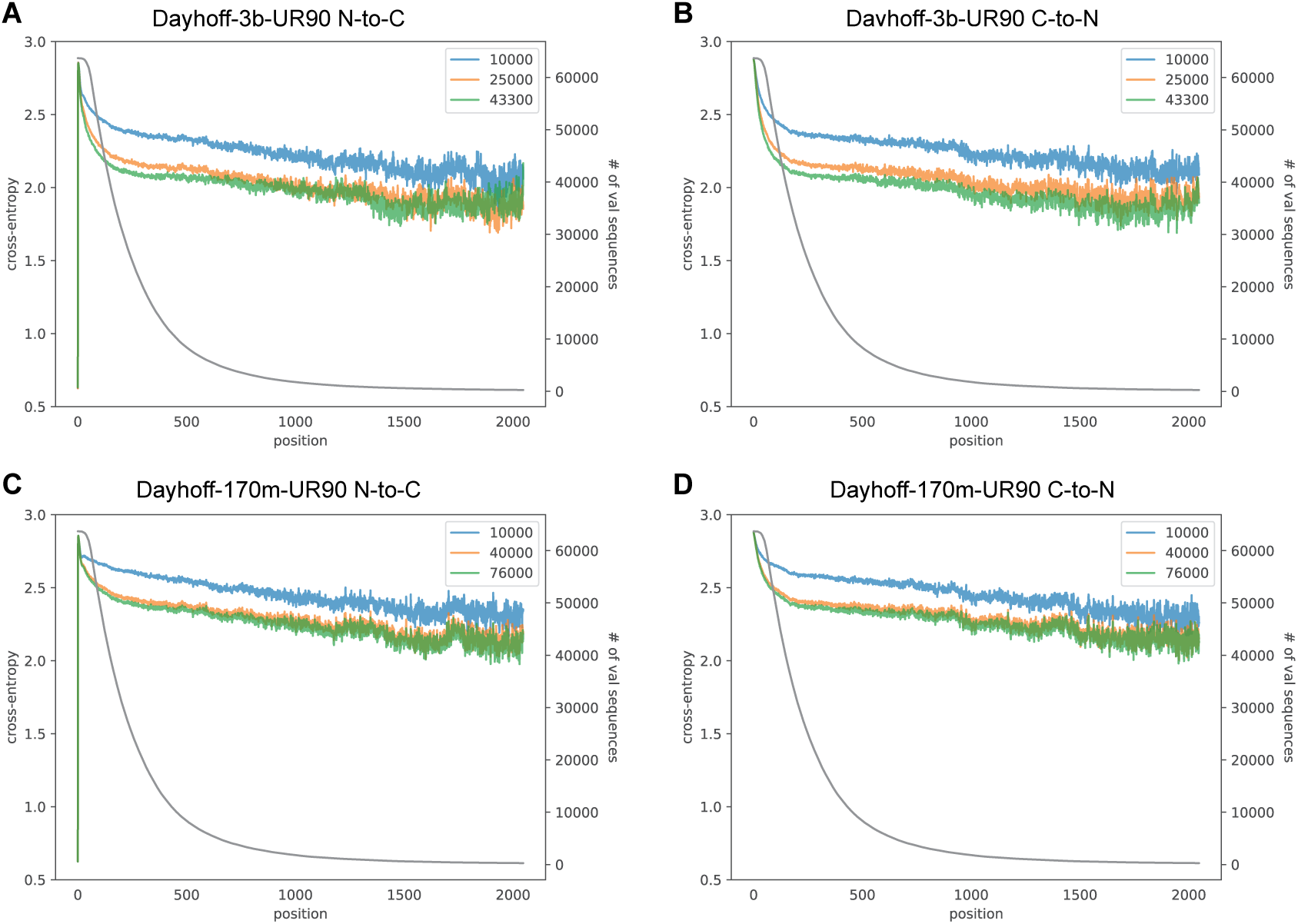
Token-wise predictive performance as a function of sequence position. Cross entropy loss (left y-axis) as a function of position (x-axis) within validation sequences for Dayhoff-3b-UR90 **(A-B)** and Dayhoff-170m-UR90 **(C-D)**, in each of the N-to-C **(A, C)** and C-to-N **(B, D)** directions. Grey lines indicate the number of validation sequences with at least a given sequence length (right x-axis). Blue, orange, and green lines indicate different model checkpoints, with the number of steps in the legends.

**Table S7:** Summary of training information for Dayhoff models. ^∗^For Dayhoff-3b-GR-HM-c, the cooldown period on UniRef90 and OpenProteinSet was performed off of Dayhoff-3b-GR-HM, which was trained on GigaRef and OpenProteinSet. Batch sizes are per GPU.

| Model | Training data | Steps | Batch size | Hardware |
| --- | --- | --- | --- | --- |
| 170m-UR50 | UR50 | 76,000 | 36,000 | 32 × 80GB H100 |
| 170m-UR90 | UR90 | 76,000 | 36,000 | 32 × 80GB H100 |
| 170m-GR | GR | 76,000 | 36,000 | 32 × 80GB H100 |
| 170m-UR50-BRu | UR50 + BRu | 76,000 | 36,000 | 32 × 80GB H100 |
| 170m-UR50-BRq | UR50 + BRq | 76,000 | 36,000 | 32 × 80GB H100 |
| 170m-UR50-BRn | UR50 + BRn | 76,000 | 36,000 | 32 × 80GB H100 |
| 170m-GRs | GRs | 112,000 | 36,000 | 4 × 40GB A100 |
| 3b-UR90 | UR90 | 43,300 | 90,000 | 176 × 80GB H100 |
| 3b-GR-HM | GR + OpenProteinSet | 52,000 | 120,000 | 144 × 80GB H100 |
| 3b-GR-HM-c | GR + OpenProteinSet + UR90* | 89,000 | 120,000 | 144 × 80GB H100 |

**Table S8:** Model perplexity on held-out test sequences for Dayhoff models. Entries are left blank in cases where the model did not support aligned or unaligned homologs.

| Model | UniRef50 | GigaRef | Aligned homologs | Unaligned homologs |
| --- | --- | --- | --- | --- |
| 170m-UR50 | 11.62 | 11.88 |  |  |
| 170m-UR90 | 11.52 | 11.85 |  |  |
| 170m-GR | 13.67 | 9.36 |  |  |
| 170m-GRs | 15.93 | 14.45 |  |  |
| 170m-UR50-BRn | 11.78 | 12.03 |  |  |
| 170m-UR50-BRq | 11.67 | 11.91 |  |  |
| 170m-UR50-BRu | 11.66 | 11.87 |  |  |
| 3b-UR90 | 8.95 | 9.64 |  |  |
| 3b-GR-HM | 11.95 | 6.68 | 4.34 | 4.60 |
| 3b-GR-HM-c | 10.11 | 9.21 | 3.57 | 3.56 |

**Table S9:** Frechet ProtBert Distance (FPD) for generations from Dayhoff-170m-UR50 relative to test sequences.

| Model | Direction | Temperature | UniRef50 | GigaRef |
| --- | --- | --- | --- | --- |
| Dayhoff-170m-UR50 | N-to-C | 0.7 | 7.06 | 7.52 |
|  |  | 0.8 | 2.33 | 2.78 |
|  |  | 0.9 | 0.73 | 1.12 |
|  |  | 1.0 | 0.74 | 1.03 |
|  |  | 1.1 | 0.85 | 1.11 |
|  |  | 1.2 | 1.00 | 1.22 |
|  | C-to-N | 0.7 | 6.74 | 7.20 |
|  |  | 0.8 | 0.74 | 2.81 |
|  |  | 0.9 | 0.76 | 1.08 |
|  |  | 1.0 | 0.76 | 1.06 |
|  |  | 1.1 | 0.88 | 1.16 |
|  |  | 1.2 | 1.00 | 1.26 |

**Table S10:**
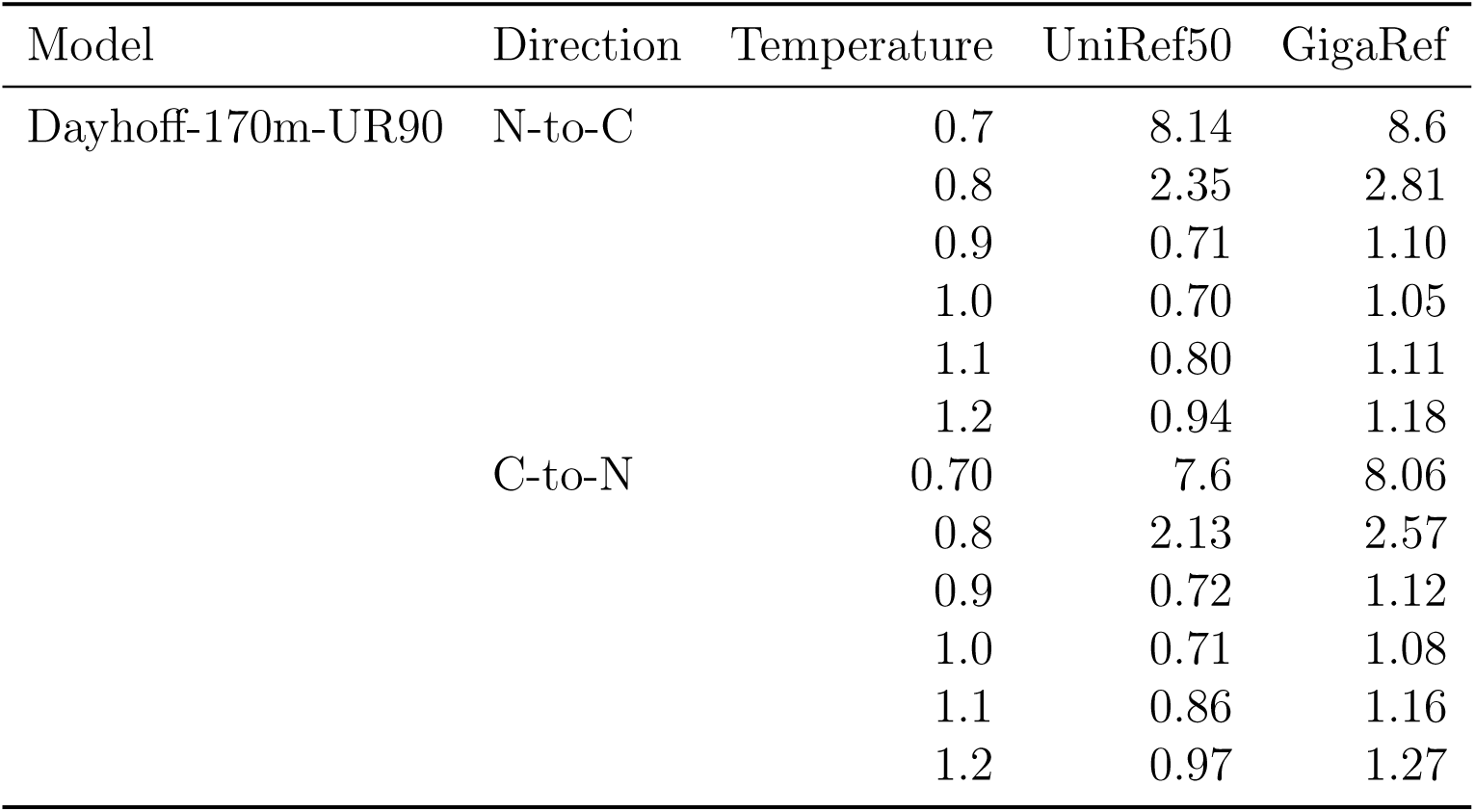
Frechet ProtBert Distance (FPD) for generations from Dayhoff-170m-UR90 relative to test sequences.

| Model | Direction | Temperature | UniRef50 | GigaRef |
| --- | --- | --- | --- | --- |
| Dayhoff-170m-UR90 | N-to-C | 0.7 | 8.14 | 8.6 |
|  |  | 0.8 | 2.35 | 2.81 |
|  |  | 0.9 | 0.71 | 1.10 |
|  |  | 1.0 | 0.70 | 1.05 |
|  |  | 1.1 | 0.80 | 1.11 |
|  |  | 1.2 | 0.94 | 1.18 |
|  | C-to-N | 0.70 | 7.6 | 8.06 |
|  |  | 0.8 | 2.13 | 2.57 |
|  |  | 0.9 | 0.72 | 1.12 |
|  |  | 1.0 | 0.71 | 1.08 |
|  |  | 1.1 | 0.86 | 1.16 |
|  |  | 1.2 | 0.97 | 1.27 |

**Table S11:** Frechet ProtBert Distance (FPD) for generations from Dayhoff-170m-GR relative to test sequences.

| Model | Direction | Temperature | UniRef50 | GigaRef |
| --- | --- | --- | --- | --- |
| Dayhoff-170m-GR | N-to-C | 0.7 | 3.4 | 3.47 |
|  |  | 0.8 | 1.21 | 1.17 |
|  |  | 0.9 | 0.84 | 0.78 |
|  |  | 1.0 | 0.9 | 0.85 |
|  |  | 1.1 | 1.05 | 1.02 |
|  |  | 1.2 | 1.21 | 1.18 |
|  | C-to-N | 0.7 | 3.44 | 3.45 |
|  |  | 0.8 | 1.33 | 1.27 |
|  |  | 0.9 | 0.89 | 0.83 |
|  |  | 1.0 | 0.94 | 0.91 |
|  |  | 1.1 | 1.17 | 1.13 |
|  |  | 1.2 | 1.31 | 1.29 |

**Table S12:**
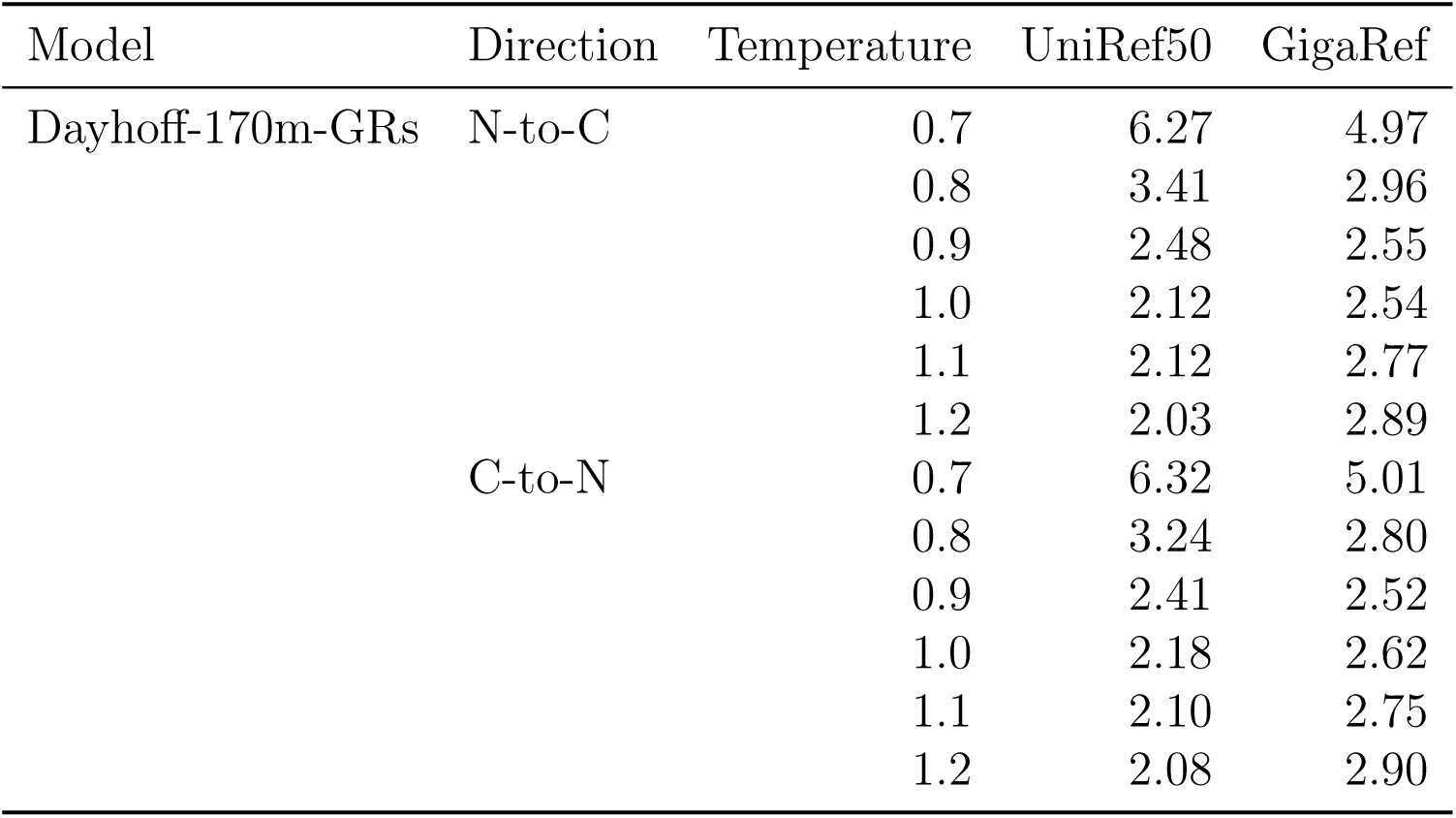
Frechet ProtBert Distance (FPD) for generations from Dayhoff-170m-GRs relative to test sequences.

| Model | Direction | Temperature | UniRef50 | GigaRef |
| --- | --- | --- | --- | --- |
| Dayhoff-170m-GRs | N-to-C | 0.7 | 6.27 | 4.97 |
|  |  | 0.8 | 3.41 | 2.96 |
|  |  | 0.9 | 2.48 | 2.55 |
|  |  | 1.0 | 2.12 | 2.54 |
|  |  | 1.1 | 2.12 | 2.77 |
|  |  | 1.2 | 2.03 | 2.89 |
|  | C-to-N | 0.7 | 6.32 | 5.01 |
|  |  | 0.8 | 3.24 | 2.80 |
|  |  | 0.9 | 2.41 | 2.52 |
|  |  | 1.0 | 2.18 | 2.62 |
|  |  | 1.1 | 2.10 | 2.75 |
|  |  | 1.2 | 2.08 | 2.90 |

**Table S13:** Frechet ProtBert Distance (FPD) for generations from Dayhoff-170m-UR50-BRu relative to test sequences.

| Model | Direction | Temperature | UniRef50 | GigaRef |
| --- | --- | --- | --- | --- |
| Dayhoff-170m-UR50-BRu | N-to-C | 0.7 | 7.08 | 7.55 |
|  |  | 0.8 | 2.72 | 3.16 |
|  |  | 0.9 | 1.21 | 1.59 |
|  |  | 1.0 | 1.04 | 1.37 |
|  |  | 1.1 | 1.13 | 1.44 |
|  |  | 1.2 | 1.11 | 1.37 |
|  | C-to-N | 0.7 | 7.77 | 8.22 |
|  |  | 0.8 | 3.32 | 3.71 |
|  |  | 0.9 | 1.67 | 2.05 |
|  |  | 1.0 | 1.28 | 1.61 |
|  |  | 1.1 | 1.11 | 1.43 |
|  |  | 1.2 | 1.08 | 1.39 |

**Table S14:** Frechet ProtBert Distance (FPD) for generations from Dayhoff-170m-UR50-BRq relative to test sequences.

| Model | Direction | Temperature | UniRef50 | GigaRef |
| --- | --- | --- | --- | --- |
| Dayhoff-170m-UR50-BRq | N-to-C | 0.7 | 7.77 | 8.26 |
|  |  | 0.8 | 3.19 | 3.67 |
|  |  | 0.9 | 1.16 | 1.56 |
|  |  | 1.0 | 0.97 | 1.33 |
|  |  | 1.1 | 1.02 | 1.33 |
|  |  | 1.2 | 1.07 | 1.35 |
|  | C-to-N | 0.7 | 7.28 | 7.74 |
|  |  | 0.8 | 3.30 | 3.76 |
|  |  | 0.9 | 1.36 | 1.75 |
|  |  | 1.0 | 1.04 | 1.40 |
|  |  | 1.1 | 0.98 | 1.30 |
|  |  | 1.2 | 1.03 | 1.33 |

**Table S15:** Frechet ProtBert Distance (FPD) for generations from Dayhoff-170m-UR50-BRn relative to test sequences.

| Model | Direction | Temperature | UniRef50 | GigaRef |
| --- | --- | --- | --- | --- |
| Dayhoff-170m-UR50-BRn | N-to-C | 0.7 | 7.32 | 7.81 |
|  |  | 0.8 | 2.87 | 3.32 |
|  |  | 0.9 | 1.34 | 1.75 |
|  |  | 1.0 | 1.11 | 1.42 |
|  |  | 1.1 | 1.14 | 1.41 |
|  |  | 1.2 | 1.23 | 1.45 |
|  | C-to-N | 0.7 | 6.59 | 7.01 |
|  |  | 0.8 | 2.85 | 3.27 |
|  |  | 0.9 | 1.28 | 1.64 |
|  |  | 1.0 | 1.04 | 1.37 |
|  |  | 1.1 | 1.05 | 1.33 |
|  |  | 1.2 | 1.09 | 1.33 |

**Table S16:**
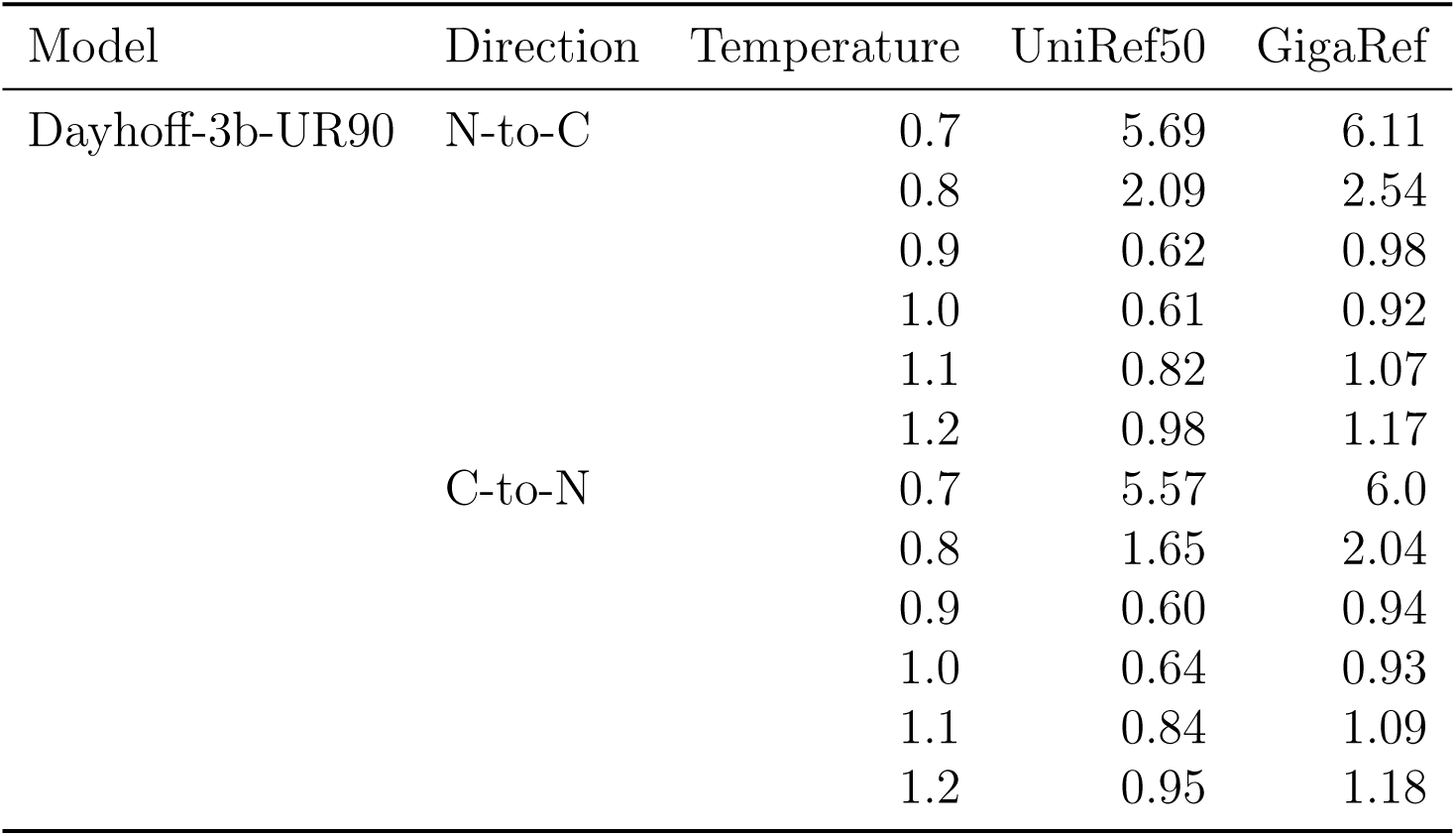
Frechet ProtBert Distance (FPD) for generations from Dayhoff-3b-UR90 relative to test sequences.

| Model | Direction | Temperature | UniRef50 | GigaRef |
| --- | --- | --- | --- | --- |
| Dayhoff-3b-UR90 | N-to-C | 0.7 | 5.69 | 6.11 |
|  |  | 0.8 | 2.09 | 2.54 |
|  |  | 0.9 | 0.62 | 0.98 |
|  |  | 1.0 | 0.61 | 0.92 |
|  |  | 1.1 | 0.82 | 1.07 |
|  |  | 1.2 | 0.98 | 1.17 |
|  | C-to-N | 0.7 | 5.57 | 6.0 |
|  |  | 0.8 | 1.65 | 2.04 |
|  |  | 0.9 | 0.60 | 0.94 |
|  |  | 1.0 | 0.64 | 0.93 |
|  |  | 1.1 | 0.84 | 1.09 |
|  |  | 1.2 | 0.95 | 1.18 |

**Table S17:**
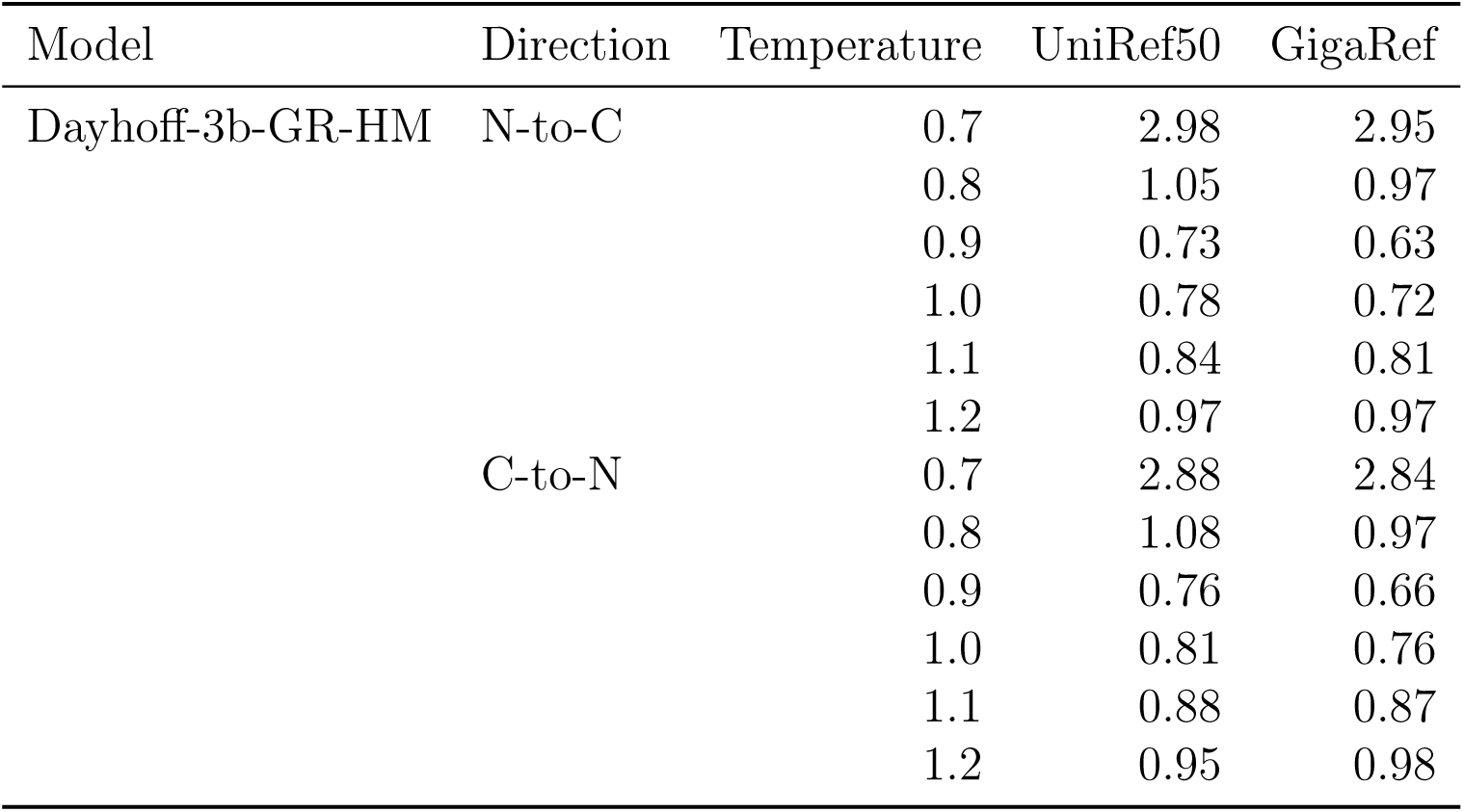
Frechet ProtBert Distance (FPD) for generations from Dayhoff-3b-GR-HM relative to test sequences.

**Table S18:** Frechet ProtBert Distance (FPD) for generations from Dayhoff-3b-GR-HM-c relative to test sequences.

| Model | Direction | Temperature | UniRef50 | GigaRef |
| --- | --- | --- | --- | --- |
| Dayhoff-3b-GR-HM-c | N-to-C | 0.7 | 6.2 | 6.64 |
|  |  | 0.8 | 2.1 | 2.51 |
|  |  | 0.9 | 0.63 | 0.99 |
|  |  | 1.0 | 0.67 | 0.97 |
|  |  | 1.1 | 0.81 | 1.06 |
|  |  | 1.2 | 0.97 | 1.17 |
|  | C-to-N | 0.7 | 7.08 | 7.52 |
|  |  | 0.8 | 2.44 | 2.82 |
|  |  | 0.9 | 0.67 | 1.03 |
|  |  | 1.0 | 0.65 | 0.99 |
|  |  | 1.1 | 0.84 | 1.11 |
|  |  | 1.2 | 0.97 | 1.23 |

**Table S19:** Quality of generated sequences as measured by ESMFold pLDDT and scPerplexity. Dataset statistics are for 1024 randomly-sampled sequences. Model statistics are for 1024 generations at temperature 1.0 in the N-to-C direction.

| Model or dataset | pLDDT | scPerplexity |
| --- | --- | --- |
| UniRef50 | $0.653 \pm 0.196$ | $9.45 \pm 2.89$ |
| GigaRef-clusters | $0.619 \pm 0.199$ | $9.69 \pm 2.83$ |
| GigaRef-singletons | $0.561 \pm 0.201$ | $10.07 \pm 2.88$ |
| 170m-UR50 | $0.421 \pm 0.132$ | $11.97 \pm 2.14$ |
| 170m-UR90 | $0.407 \pm 0.125$ | $12.12 \pm 2.14$ |
| 170m-GR | $0.422 \pm 0.129$ | $11.83 \pm 2.12$ |
| 170m-UR50-BR<sub>u</sub> | $0.441 \pm 0.157$ | $11.71 \pm 2.18$ |
| 170m-UR50-BR<sub>q</sub> | $0.434 \pm 0.152$ | $11.72 \pm 2.24$ |
| 170m-UR50-BR<sub>n</sub> | $0.432 \pm 0.131$ | $11.77 \pm 2.24$ |
| 3b-UR90 | $0.454 \pm 0.150$ | $11.79 \pm 2.38$ |
| 3b-GR-HM | $0.406 \pm 0.126$ | $11.50 \pm 2.16$ |
| 3b-GR-HM-c | $0.423 \pm 0.132$ | $11.91 \pm 2.18$ |

**Table S20:** Dayhoff-170m-UR50 generation fidelity, as measured by ESMFold pLDDT and scPerplexity of unconditionally generated sequences.

| Model | Direction | Temperature | pLDDT | scPerplexity |
| --- | --- | --- | --- | --- |
| Dayhoff-170m-UR50 | N-to-C | 0.7 | $0.48 \pm 0.19$ | $9.42 \pm 3.44$ |
| | | 0.8 | $0.48 \pm 0.17$ | $10.71 \pm 2.84$ |
| | | 0.9 | $0.45 \pm 0.15$ | $11.55 \pm 2.33$ |
| | | 1.0 | $0.42 \pm 0.13$ | $12.02 \pm 2.09$ |
| | | 1.1 | $0.41 \pm 0.12$ | $12.19 \pm 2.03$ |
| | | 1.2 | $0.40 \pm 0.11$ | $12.26 \pm 1.97$ |
| | C-to-N | 0.7 | $0.49 \pm 0.19$ | $9.24 \pm 3.47$ |
| | | 0.8 | $0.49 \pm 0.16$ | $10.55 \pm 2.91$ |
| | | 0.9 | $0.46 \pm 0.15$ | $11.56 \pm 2.31$ |
| | | 1.0 | $0.42 \pm 0.14$ | $11.92 \pm 2.19$ |
| | | 1.1 | $0.39 \pm 0.12$ | $12.33 \pm 2.04$ |
| | | 1.2 | $0.38 \pm 0.12$ | $12.53 \pm 1.99$ |

**Table S21:** Dayhoff-170m-UR90 generation fidelity, as measured by ESMFold pLDDT and scPerplexity of unconditionally generated sequences.

| Model | Direction | Temperature | pLDDT | scPerplexity |
| --- | --- | --- | --- | --- |
| Dayhoff-170m-UR90 | N-to-C | 0.7 | $0.48 \pm 0.19$ | $9.09 \pm 3.38$ |
| | | 0.8 | $0.48 \pm 0.18$ | $10.50 \pm 2.87$ |
| | | 0.9 | $0.44 \pm 0.15$ | $11.72 \pm 2.39$ |
| | | 1.0 | $0.41 \pm 0.12$ | $12.08 \pm 2.11$ |
| | | 1.1 | $0.39 \pm 0.11$ | $12.36 \pm 1.99$ |
| | | 1.2 | $0.39 \pm 0.11$ | $12.38 \pm 1.92$ |
| | C-to-N | 0.7 | $0.47 \pm 0.19$ | $9.06 \pm 3.63$ |
| | | 0.8 | $0.49 \pm 0.16$ | $10.74 \pm 2.95$ |
| | | 0.9 | $0.45 \pm 0.15$ | $11.61 \pm 2.47$ |
| | | 1.0 | $0.40 \pm 0.13$ | $12.15 \pm 2.17$ |
| | | 1.1 | $0.37 \pm 0.11$ | $12.50 \pm 1.96$ |
| | | 1.2 | $0.35 \pm 0.10$ | $12.67 \pm 1.91$ |

**Table S22:** Dayhoff-170m-GR generation fidelity, as measured by ESMFold pLDDT and scPerplexity of unconditionally generated sequences.

| Model | Direction | Temperature | pLDDT | scPerplexity |
| --- | --- | --- | --- | --- |
| Dayhoff-170m-GR | N-to-C | 0.7 | $0.51 \pm 0.18$ | $10.45 \pm 3.14$ |
| | | 0.8 | $0.48 \pm 0.17$ | $11.10 \pm 2.66$ |
| | | 0.9 | $0.45 \pm 0.15$ | $11.48 \pm 2.31$ |
| | | 1.0 | $0.43 \pm 0.13$ | $11.8 \pm 2.18$ |
| | | 1.1 | $0.42 \pm 0.13$ | $11.99 \pm 2.04$ |
| | | 1.2 | $0.41 \pm 0.13$ | $12.17 \pm 1.96$ |
| | C-to-N | 0.7 | $0.51 \pm 0.17$ | $10.53 \pm 2.97$ |
| | | 0.8 | $0.47 \pm 0.15$ | $11.32 \pm 2.51$ |
| | | 0.9 | $0.44 \pm 0.14$ | $11.71 \pm 2.24$ |
| | | 1.0 | $0.42 \pm 0.13$ | $11.86 \pm 2.07$ |
| | | 1.1 | $0.41 \pm 0.13$ | $12.22 \pm 2.94$ |
| | | 1.2 | $0.41 \pm 0.13$ | $12.32 \pm 2.92$ |

**Table S23:**
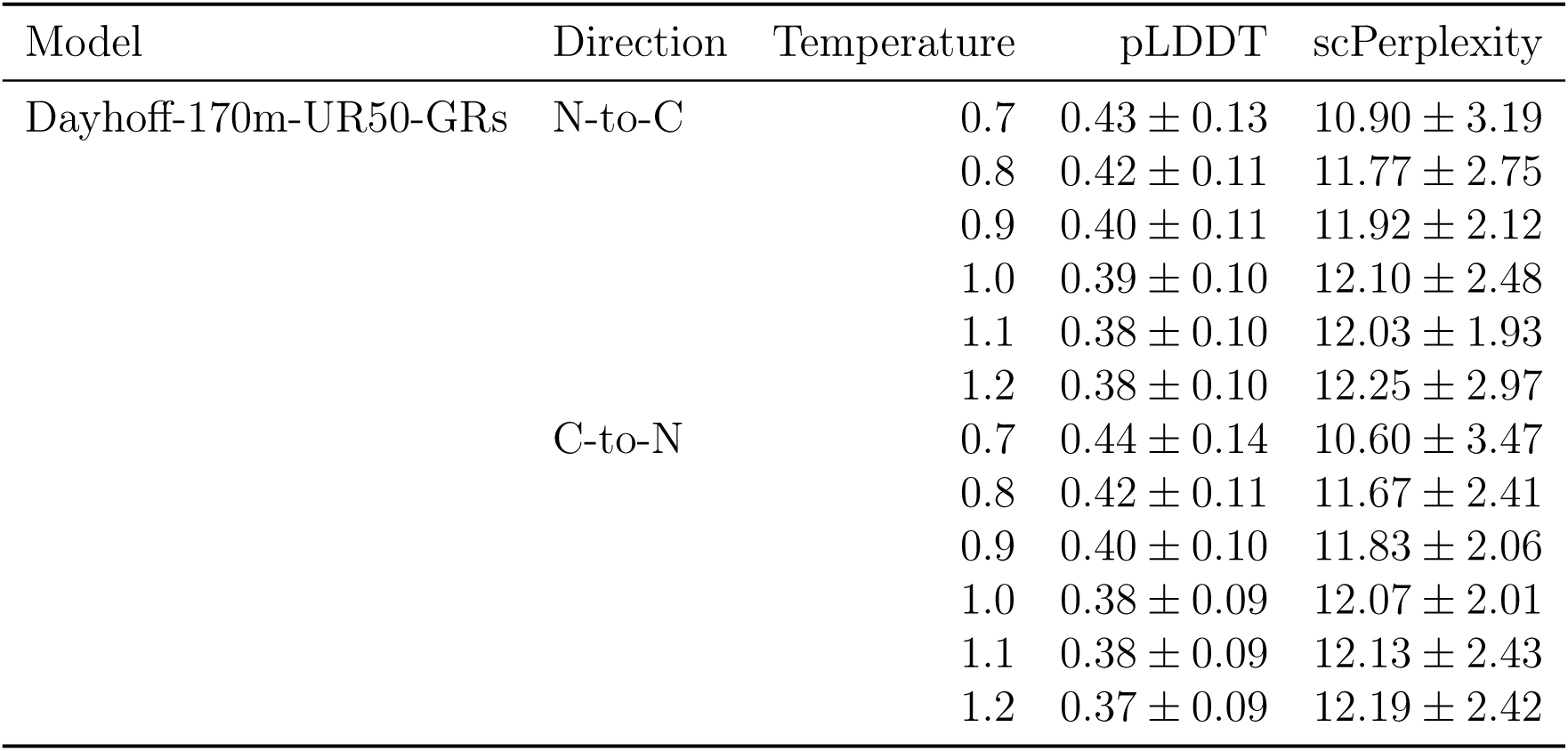
Dayhoff-170m-UR50-GRs generation fidelity, as measured by ESMFold pLDDT and scPerplexity of unconditionally generated sequences.

| Model | Direction | Temperature | pLDDT | scPerplexity |
| --- | --- | --- | --- | --- |
| Dayhoff-170m-UR50-GRs | N-to-C | 0.7 | $0.43 \pm 0.13$ | $10.90 \pm 3.19$ |
| | | 0.8 | $0.42 \pm 0.11$ | $11.77 \pm 2.75$ |
| | | 0.9 | $0.40 \pm 0.11$ | $11.92 \pm 2.12$ |
| | | 1.0 | $0.39 \pm 0.10$ | $12.10 \pm 2.48$ |
| | | 1.1 | $0.38 \pm 0.10$ | $12.03 \pm 1.93$ |
| | | 1.2 | $0.38 \pm 0.10$ | $12.25 \pm 2.97$ |
| | C-to-N | 0.7 | $0.44 \pm 0.14$ | $10.60 \pm 3.47$ |
| | | 0.8 | $0.42 \pm 0.11$ | $11.67 \pm 2.41$ |
| | | 0.9 | $0.40 \pm 0.10$ | $11.83 \pm 2.06$ |
| | | 1.0 | $0.38 \pm 0.09$ | $12.07 \pm 2.01$ |
| | | 1.1 | $0.38 \pm 0.09$ | $12.13 \pm 2.43$ |
| | | 1.2 | $0.37 \pm 0.09$ | $12.19 \pm 2.42$ |

**Table S24:** Dayhoff-170m-UR50-BRu generation fidelity, as measured by ESMFold pLDDT and scPerplexity of unconditionally generated sequences.

| Model | Direction | Temperature | pLDDT | scPerplexity |
| --- | --- | --- | --- | --- |
| Dayhoff-170m-UR50-BRu | N-to-C | 0.7 | $0.52 \pm 0.21$ | $8.93 \pm 3.18$ |
| | | 0.8 | $0.50 \pm 0.18$ | $10.49 \pm 2.83$ |
| | | 0.9 | $0.48 \pm 0.17$ | $11.29 \pm 2.37$ |
| | | 1.0 | $0.43 \pm 0.15$ | $11.75 \pm 2.16$ |
| | | 1.1 | $0.42 \pm 0.14$ | $11.97 \pm 2.09$ |
| | | 1.2 | $0.40 \pm 0.13$ | $12.10 \pm 1.92$ |
| | C-to-N | 0.7 | $0.56 \pm 0.21$ | $9.29 \pm 3.22$ |
| | | 0.8 | $0.54 \pm 0.19$ | $10.49 \pm 2.69$ |
| | | 0.9 | $0.49 \pm 0.18$ | $11.31 \pm 2.33$ |
| | | 1.0 | $0.45 \pm 0.16$ | $11.66 \pm 2.20$ |
| | | 1.1 | $0.40 \pm 0.14$ | $12.10 \pm 2.08$ |
| | | 1.2 | $0.37 \pm 0.13$ | $12.35 \pm 2.00$ |

**Table S25:** Dayhoff-170m-UR50-BRq generation fidelity, as measured by ESMFold pLDDT and scPerplexity of unconditionally generated sequences.

| Model | Direction | Temperature | pLDDT | scPerplexity |
| --- | --- | --- | --- | --- |
| Dayhoff-170m-UR50-BRq | N-to-C | 0.7 | $0.51 \pm 0.21$ | $9.27 \pm 3.20$ |
| | | 0.8 | $0.49 \pm 0.18$ | $10.57 \pm 2.79$ |
| | | 0.9 | $0.47 \pm 0.17$ | $11.40 \pm 2.48$ |
| | | 1.0 | $0.44 \pm 0.15$ | $11.65 \pm 2.28$ |
| | | 1.1 | $0.42 \pm 0.13$ | $11.92 \pm 2.24$ |
| | | 1.2 | $0.40 \pm 0.12$ | $12.09 \pm 2.03$ |
| | C-to-N | 0.7 | $0.56 \pm 0.21$ | $9.17 \pm 3.14$ |
| | | 0.8 | $0.53 \pm 0.19$ | $10.20 \pm 2.73$ |
| | | 0.9 | $0.48 \pm 0.17$ | $11.22 \pm 2.42$ |
| | | 1.0 | $0.43 \pm 0.15$ | $11.80 \pm 2.19$ |
| | | 1.1 | $0.40 \pm 0.13$ | $12.12 \pm 2.08$ |
| | | 1.2 | $0.37 \pm 0.12$ | $12.38 \pm 2.03$ |

**Table S26:** Dayhoff-170m-UR50-BRn generation fidelity, as measured by ESMFold pLDDT and scPerplexity of unconditionally generated sequences.

| Model | Direction | Temperature | pLDDT | scPerplexity |
| --- | --- | --- | --- | --- |
| Dayhoff-170m-UR50-BRn | N-to-C | 0.7 | $0.48 \pm 0.18$ | $9.45 \pm 3.21$ |
| | | 0.8 | $0.48 \pm 0.16$ | $10.51 \pm 2.83$ |
| | | 0.9 | $0.46 \pm 0.14$ | $11.34 \pm 2.50$ |
| | | 1.0 | $0.43 \pm 0.13$ | $11.76 \pm 2.26$ |
| | | 1.1 | $0.42 \pm 0.12$ | $11.81 \pm 2.15$ |
| | | 1.2 | $0.42 \pm 0.12$ | $12.09 \pm 2.00$ |
| | C-to-N | 0.7 | $0.52 \pm 0.17$ | $9.12 \pm 3.20$ |
| | | 0.8 | $0.50 \pm 0.15$ | $10.33 \pm 2.76$ |
| | | 0.9 | $0.47 \pm 0.13$ | $11.28 \pm 2.43$ |
| | | 1.0 | $0.43 \pm 0.13$ | $11.78 \pm 2.22$ |
| | | 1.1 | $0.41 \pm 0.13$ | $12.09 \pm 2.05$ |
| | | 1.2 | $0.39 \pm 0.13$ | $12.36 \pm 1.95$ |

**Table S27:** Dayhoff-3b-UR90 generation fidelity, as measured by ESMFold pLDDT and scPerplexity of unconditionally generated sequences.

| Model | Direction | Temperature | pLDDT | scPerplexity |
| --- | --- | --- | --- | --- |
| Dayhoff-3b-UR90 | N-to-C | 0.7 | $0.55 \pm 0.22$ | $8.91 \pm 3.30$ |
| | | 0.8 | $0.52 \pm 0.19$ | $10.45 \pm 2.96$ |
| | | 0.9 | $0.48 \pm 0.17$ | $11.22 \pm 2.60$ |
| | | 1.0 | $0.45 \pm 0.15$ | $11.77 \pm 2.44$ |
| | | 1.1 | $0.44 \pm 0.14$ | $11.93 \pm 2.22$ |
| | | 1.2 | $0.43 \pm 0.13$ | $11.99 \pm 2.24$ |
| | C-to-N | 0.7 | $0.53 \pm 0.21$ | $9.02 \pm 3.45$ |
| | | 0.8 | $0.53 \pm 0.18$ | $10.51 \pm 2.99$ |
| | | 0.9 | $0.50 \pm 0.17$ | $11.25 \pm 2.63$ |
| | | 1.0 | $0.45 \pm 0.15$ | $11.80 \pm 2.32$ |
| | | 1.1 | $0.42 \pm 0.13$ | $12.02 \pm 2.13$ |
| | | 1.2 | $0.41 \pm 0.13$ | $12.30 \pm 2.11$ |

**Table S28:**
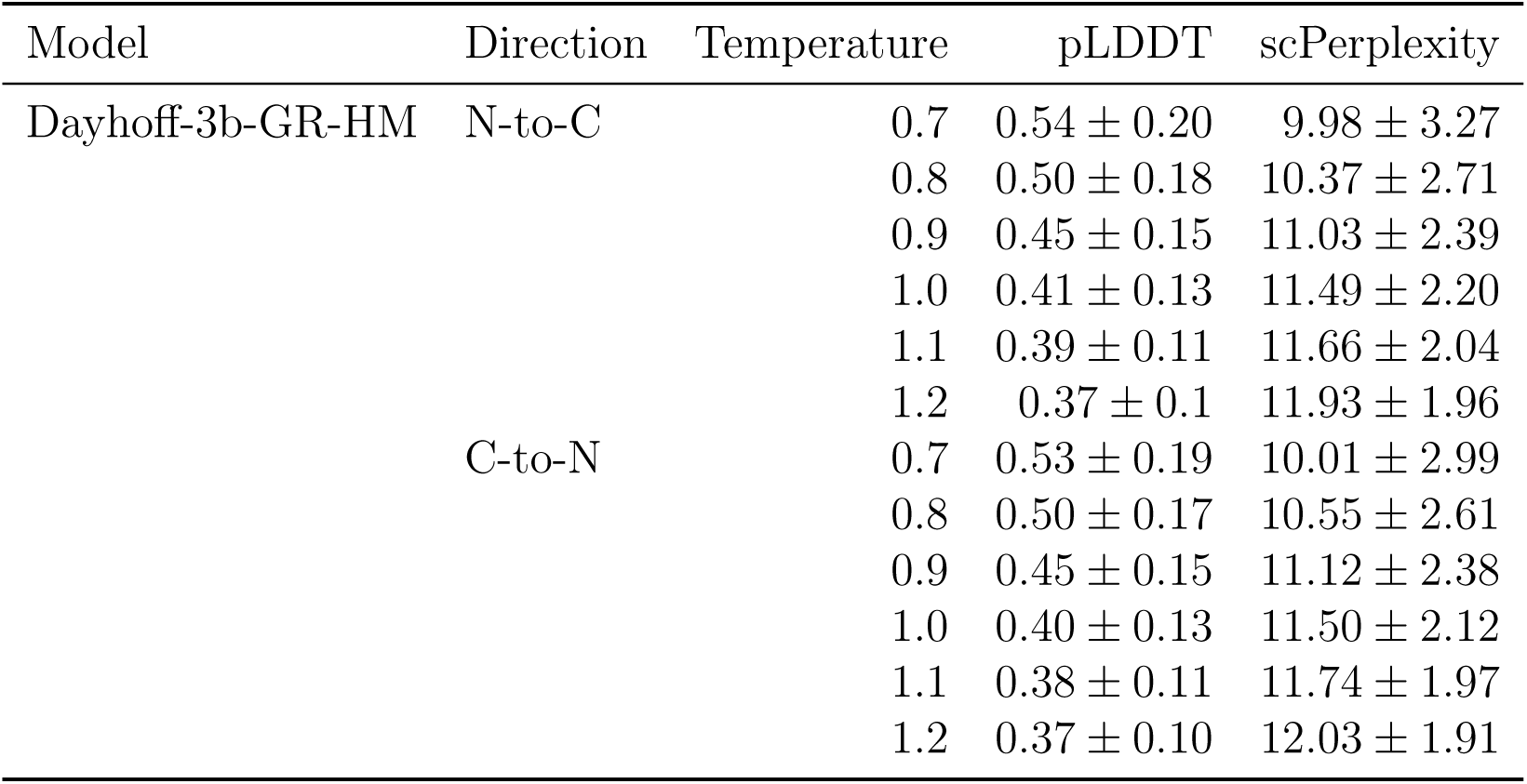
Dayhoff-3b-GR-HM generation fidelity, as measured by ESMFold pLDDT and scPerplexity of unconditionally generated sequences.

| Model | Direction | Temperature | pLDDT | scPerplexity |
| --- | --- | --- | --- | --- |
| Dayhoff-3b-GR-HM | N-to-C | 0.7 | $0.54 \pm 0.20$ | $9.98 \pm 3.27$ |
| | | 0.8 | $0.50 \pm 0.18$ | $10.37 \pm 2.71$ |
| | | 0.9 | $0.45 \pm 0.15$ | $11.03 \pm 2.39$ |
| | | 1.0 | $0.41 \pm 0.13$ | $11.49 \pm 2.20$ |
| | | 1.1 | $0.39 \pm 0.11$ | $11.66 \pm 2.04$ |
| | | 1.2 | $0.37 \pm 0.1$ | $11.93 \pm 1.96$ |
| | C-to-N | 0.7 | $0.53 \pm 0.19$ | $10.01 \pm 2.99$ |
| | | 0.8 | $0.50 \pm 0.17$ | $10.55 \pm 2.61$ |
| | | 0.9 | $0.45 \pm 0.15$ | $11.12 \pm 2.38$ |
| | | 1.0 | $0.40 \pm 0.13$ | $11.50 \pm 2.12$ |
| | | 1.1 | $0.38 \pm 0.11$ | $11.74 \pm 1.97$ |
| | | 1.2 | $0.37 \pm 0.10$ | $12.03 \pm 1.91$ |

**Table S29:**
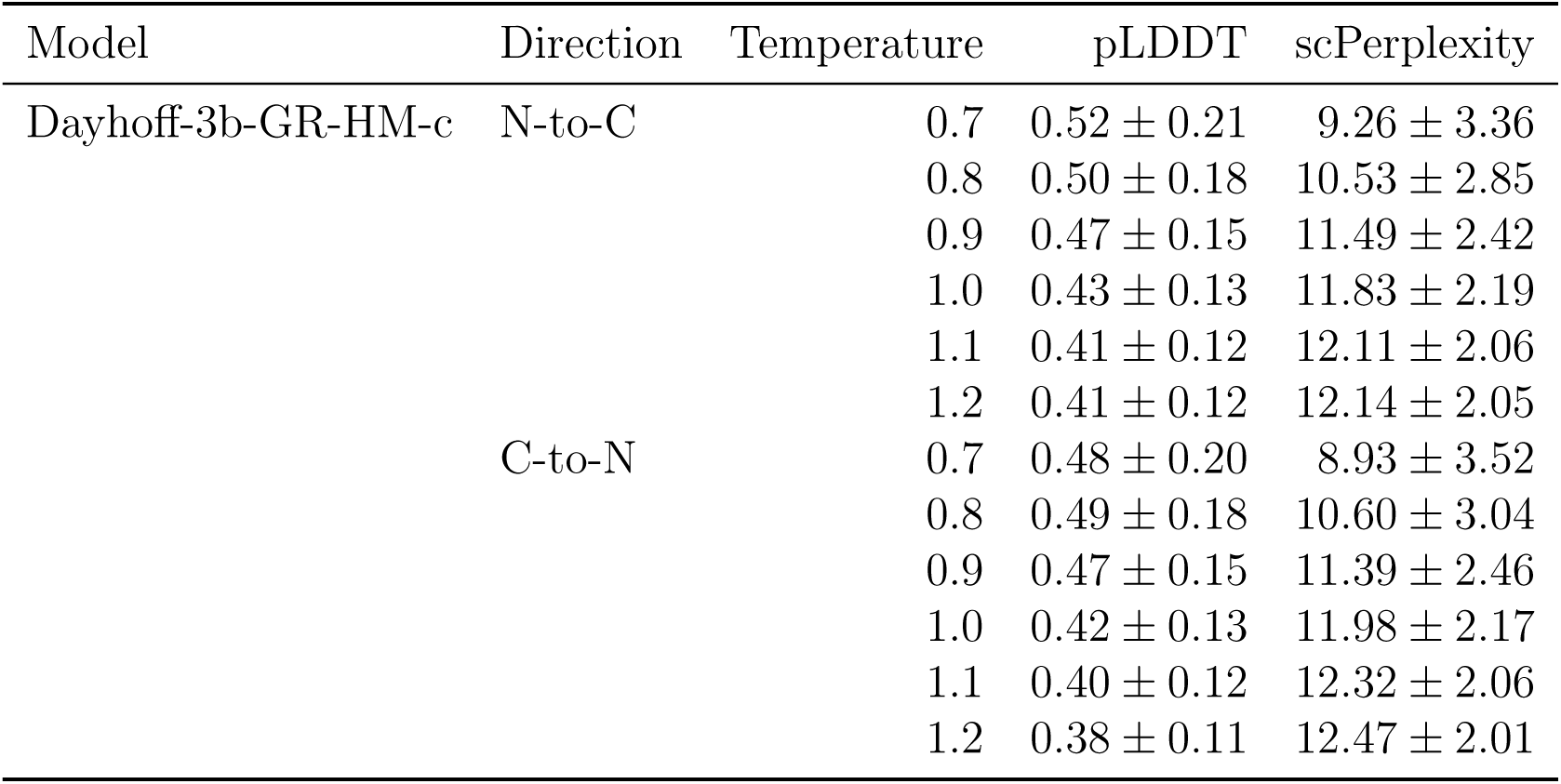
Dayhoff-3b-GR-HM-c generation fidelity, as measured by ESMFold pLDDT and scPerplexity of unconditionally generated sequences.

| Model | Direction | Temperature | pLDDT | scPerplexity |
| --- | --- | --- | --- | --- |
| Dayhoff-3b-GR-HM-c | N-to-C | 0.7 | $0.52 \pm 0.21$ | $9.26 \pm 3.36$ |
| | | 0.8 | $0.50 \pm 0.18$ | $10.53 \pm 2.85$ |
| | | 0.9 | $0.47 \pm 0.15$ | $11.49 \pm 2.42$ |
| | | 1.0 | $0.43 \pm 0.13$ | $11.83 \pm 2.19$ |
| | | 1.1 | $0.41 \pm 0.12$ | $12.11 \pm 2.06$ |
| | | 1.2 | $0.41 \pm 0.12$ | $12.14 \pm 2.05$ |
| | C-to-N | 0.7 | $0.48 \pm 0.20$ | $8.93 \pm 3.52$ |
| | | 0.8 | $0.49 \pm 0.18$ | $10.60 \pm 3.04$ |
| | | 0.9 | $0.47 \pm 0.15$ | $11.39 \pm 2.46$ |
| | | 1.0 | $0.42 \pm 0.13$ | $11.98 \pm 2.17$ |
| | | 1.1 | $0.40 \pm 0.12$ | $12.32 \pm 2.06$ |
| | | 1.2 | $0.38 \pm 0.11$ | $12.47 \pm 2.01$ |

**Table S30:** Summary of sequence representation in DayhoffRef, by generating model and sampling parameters.

| Direction | N-to-C |  | C-to-N |  | Total |
| --- | --- | --- | --- | --- | --- |
|  | 1.0 | 0.9 | 1.0 | 0.9 |  |
| 170m-UR50-BRn | 1,001,024 | 976,024 | 1,204,524 | 1,001,024 | 4,182,596 |
| 3b-UR90 | 1,543,438 | 418,304 | 284,858 | 834,908 | 3,081,508 |
| 3b-GR-HM | 2,043,840 | 1,775,515 | 993,702 | 886,864 | 5,699,921 |
| 3b-GR-HM-c | 822,500 | 961,100 | 780,900 | 1,054,500 | 3,619,000 |
| **Total** | 5,410,802 | 4,130,943 | 3,263,984 | 3,777,296 | **16,583,025** |

**Figure S8:**
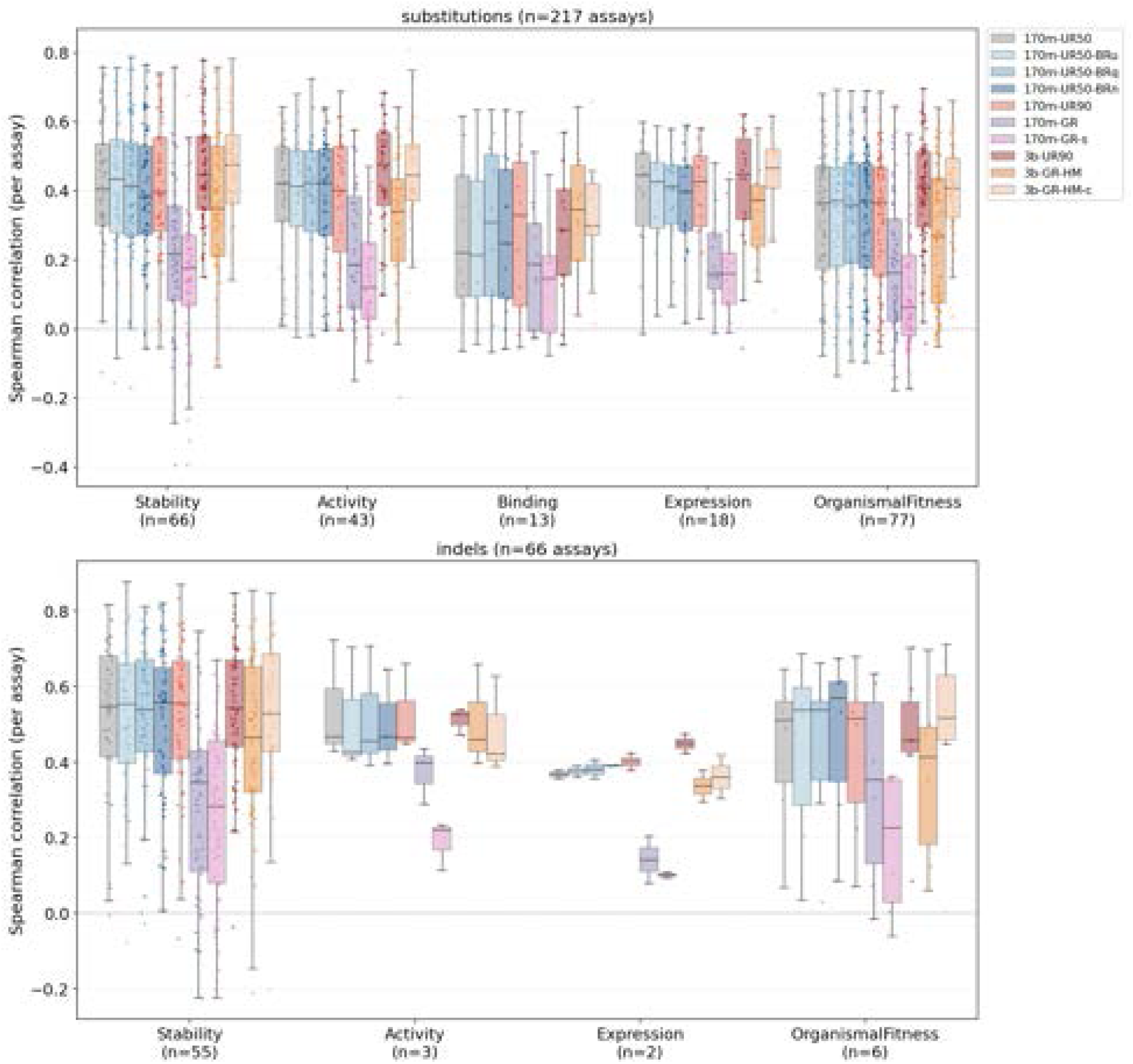
Zero-shot variant-effect performance across ProteinGym categories. Per-assay Spearman correlations, stratified by Protein Gym assay type for substitutions (top) and indels (bottom). Boxes show the median and interquartile range across assays; points are individual assays, n=number of assays.

**Figure S9:**
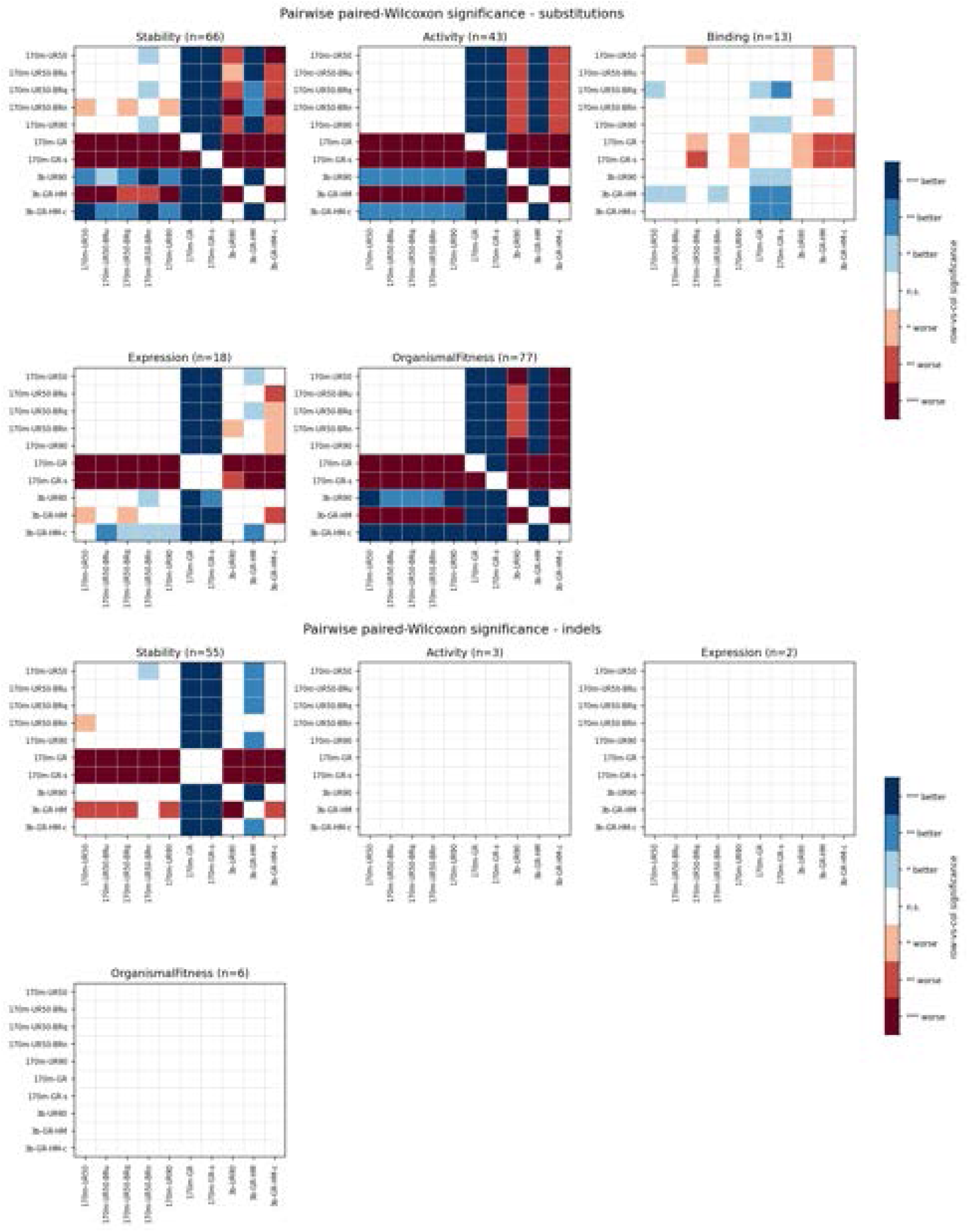
Pairwise statistical comparison of per-assay zero-shot performance on ProteinGym substitutions (top) and indels (bottom). Each panel is one ProteinGym selection type (n = number of assays). For each model pair we compared per-assay Spearman correlations over their shared assays using a two-sided paired Wilcoxon signed-rank test. Blue = row model significantly better than column model; red = significantly worse; white = not significant or diagonal. Intensity denotes the significance tier (* p*<*0.05, ** p*<*0.01, *** p*<*0.001, *n.s.*). Pairs with fewer than six shared assays were not tested.

**Table S31:** Summary metrics for clustering DayhoffRef at 50% sequence identity. Metrics are broken down by generating model source. Clustering was performed over the entirety of DayhoffRef. The mean cluster size is the mean size of the clusters that a given set of sequences are in; these clusters may contain sequences generated by other models. “Total” represents the mean cluster size and fraction of singletons when clustering over the entirety of DayhoffRef.

| Model | Mean cluster size | Fraction singleton |
| --- | --- | --- |
| 170m-UR50-BRn | 3.85 | 0.19 |
| 3b-UR90 | 1.23 | 0.87 |
| 3b-GR-HM | 1.20 | 0.82 |
| 3b-GR-HM-c | 3.19 | 0.29 |
| **Total** | 2.31 | 0.56 |

**Table S32:** Number of successes out of 100 model-attempted solutions and MotifBench scores for motif scaffolding problems from the RFdiffusion benchmark.

| Problem | 170m-UR50 | 170m-UR90 | 170m-GR | 170m-UR50-BRn | 3b-UR90 | 3b-GR-HM | 3b-GR-HM-c | EvoDiff-Seq |
| --- | --- | --- | --- | --- | --- | --- | --- | --- |
| 1PRW | 62 | 72 | 81 | 95 | 94 | 81 | 79 | 82 |
| 1BCF | 0 | 0 | 5 | 0 | 10 | 8 | 0 | 7 |
| 5TPN | 0 | 0 | 0 | 0 | 0 | 0 | 0 | 0 |
| 5IUS | 0 | 0 | 0 | 0 | 0 | 0 | 0 | 0 |
| 3IXT | 12 | 17 | 12 | 14 | 18 | 11 | 14 | 20 |
| 5YUI | 0 | 0 | 0 | 0 | 0 | 0 | 0 | 0 |
| 1QJG | 0 | 0 | 0 | 0 | 0 | 0 | 0 | 0 |
| 1YCR | 2 | 5 | 0 | 6 | 2 | 3 | 4 | 2 |
| 2KL8 | 0 | 1 | 0 | 1 | 1 | 1 | 1 | 1 |
| 7MRX_60 | 1 | 0 | 0 | 0 | 42 | 0 | 9 | 0 |
| 7MRX_85 | 0 | 0 | 0 | 0 | 19 | 1 | 1 | 0 |
| 7MRX_128 | 0 | 0 | 0 | 0 | 0 | 0 | 0 | 0 |
| 4JHW | 0 | 0 | 0 | 0 | 0 | 0 | 0 | 0 |
| 4ZYP | 0 | 0 | 0 | 0 | 0 | 0 | 0 | 0 |
| 5WN9 | 0 | 0 | 0 | 0 | 0 | 0 | 0 | 0 |
| 6VW1 | 1 | 1 | 1 | 0 | 0 | 0 | 0 | 0 |
| 5TRV_short | 0 | 0 | 0 | 0 | 0 | 0 | 0 | 0 |
| 5TRV_med | 0 | 0 | 0 | 0 | 0 | 0 | 0 | 0 |
| 5TRV_long | 0 | 0 | 0 | 0 | 0 | 0 | 0 | 0 |
| 6E6R_short | 2 | 2 | 1 | 3 | 14 | 7 | 8 | 6 |
| 6E6R_med | 0 | 1 | 2 | 0 | 4 | 0 | 2 | 0 |
| 6E6R_long | 0 | 1 | 0 | 0 | 3 | 0 | 1 | 0 |
| 6EXZ_short | 0 | 0 | 0 | 0 | 0 | 0 | 0 | 0 |
| 6EXZ_med | 0 | 0 | 0 | 0 | 0 | 0 | 0 | 0 |
| 6EXZ_long | 0 | 0 | 0 | 0 | 0 | 0 | 0 | 0 |
| **Problems** | **6** | **8** | **6** | **5** | **10** | **7** | **9** | **6** |
| **Successes** | **80** | **100** | **102** | **119** | **207** | **112** | **119** | **118** |
| **Score** | **9.65** | **12.25** | **6.10** | **7.26** | **16.32** | **11.90** | **14.14** | **7.67** |

**Table S33:**
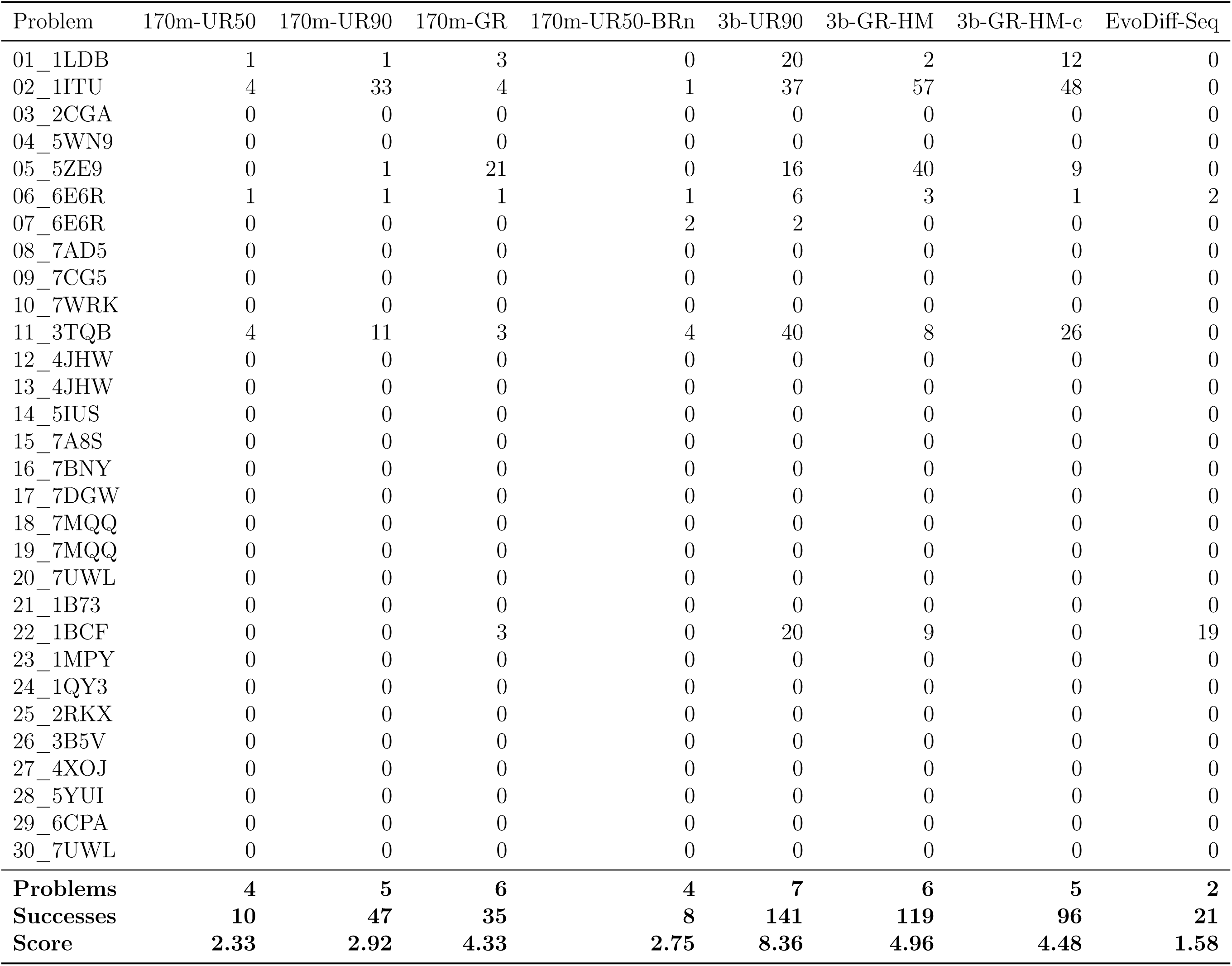
Number of successes out of 100 model-attempted solutions and MotifBench scores for motif scaffolding problems from the MotifBench benchmark.

**Table S34:** Number of successes out of 100 model-attempted solutions and MotifBench scores on the RFdiffusion benchmark for additional GRs, BR models and EvoDiff-MSA baseline.

| Problem | 170m-GRs | 170m-UR50-BRq | 170m-UR50-BRu | EvoDiff-MSA |
| --- | --- | --- | --- | --- |
| 1PRW | 60 | 91 | 90 | 82 |
| 1BCF | 0 | 0 | 0 | 100 |
| 5TPN | 0 | 0 | 0 | 0 |
| 5IUS | 0 | 0 | 0 | 5 |
| 3IXT | 11 | 18 | 12 | 11 |
| 5YUI | 0 | 0 | 0 | 4 |
| 1QJG | 0 | 0 | 0 | 25 |
| 1YCR | 4 | 7 | 6 | 0 |
| 2KL8 | 2 | 0 | 1 | 0 |
| 7MRX_60 | 0 | 0 | 2 | 67 |
| 7MRX_85 | 0 | 0 | 0 | 30 |
| 7MRX_128 | 0 | 0 | 0 | 3 |
| 4JHW | 0 | 0 | 0 | 0 |
| 4ZYP | 0 | 0 | 1 | 0 |
| 5WN9 | 0 | 0 | 0 | 0 |
| 6VW1 | 0 | 0 | 1 | 93 |
| 5TRV_short | 0 | 0 | 0 | 7 |
| 5TRV_med | 0 | 0 | 0 | 9 |
| 5TRV_long | 0 | 0 | 0 | 3 |
| 6E6R_short | 1 | 3 | 2 | 44 |
| 6E6R_med | 1 | 0 | 2 | 27 |
| 6E6R_long | 0 | 0 | 1 | 0 |
| 6EXZ_short | 0 | 0 | 0 | 61 |
| 6EXZ_med | 0 | 0 | 0 | 43 |
| 6EXZ_long | 0 | 0 | 0 | 2 |
| **Solved** | **6** | **4** | **10** | **18** |
| **Successes** | **79** | **119** | **118** | **616** |
| **Score** | **7.35** | **10.62** | **14.36** | **20.72** |

**Table S35:** Number of successes out of 100 model-attempted solutions and MotifBench scores on the MotifBench benchmark for additional GRs, BR models and EvoDiff-MSA baseline.

| Problem | 170m-GRs | 170m-UR50-BRq | 170m-UR50-BRu | EvoDiff-MSA |
| --- | --- | --- | --- | --- |
| 01_1LDB | 0 | 0 | 1 | 12 |
| 02_1ITU | 0 | 1 | 4 | 69 |
| 03_2CGA | 0 | 0 | 0 | 0 |
| 04_5WN9 | 0 | 0 | 0 | 0 |
| 05_5ZE9 | 0 | 0 | 0 | 22 |
| 06_6E6R | 0 | 2 | 1 | 23 |
| 07_6E6R | 0 | 0 | 0 | 0 |
| 08_7AD5 | 0 | 0 | 0 | 0 |
| 09_7CG5 | 0 | 0 | 0 | 0 |
| 10_7WRK | 0 | 0 | 0 | 0 |
| 11_3TQB | 0 | 3 | 7 | 59 |
| 12_4JHW | 0 | 0 | 0 | 0 |
| 13_4JHW | 0 | 0 | 0 | 0 |
| 14_5IUS | 0 | 0 | 0 | 7 |
| 15_7A8S | 0 | 0 | 0 | 2 |
| 16_7BNY | 0 | 0 | 0 | 0 |
| 17_7DGW | 0 | 0 | 0 | 11 |
| 18_7MQQ | 0 | 0 | 0 | 0 |
| 19_7MQQ | 0 | 0 | 0 | 0 |
| 20_7UWL | 0 | 0 | 0 | 0 |
| 21_1B73 | 0 | 0 | 0 | 0 |
| 22_1BCF | 0 | 0 | 0 | 100 |
| 23_1MPY | 0 | 0 | 0 | 46 |
| 24_1QY3 | 0 | 0 | 0 | 0 |
| 25_2RKX | 0 | 0 | 0 | 27 |
| 26_3B5V | 0 | 0 | 0 | 0 |
| 27_4XOJ | 0 | 0 | 0 | 35 |
| 28_5YUI | 0 | 0 | 0 | 5 |
| 29_6CPA | 0 | 0 | 0 | 98 |
| 30_7UWL | 0 | 0 | 0 | 0 |
| **Problems** | **0** | **3** | **4** | **14** |
| **Successes** | **0** | **6** | **13** | **516** |
| **Score** | **0.00** | **2.17** | **2.75** | **12.45** |

**Table S36:** Evolutionary conditioning results. TM-score is between ESMFold predictions for the generated sequence and the original query. Identity to closest conditioning sequence is from BioPython pairwise alignment with default parameters. Aligned and unaligned homolog conditioning values are from Dayhoff-3b-GR-HM-c.

|  | ESMFold pLDDT | scPerplexity | TM-score | Identity |
| --- | --- | --- | --- | --- |
| Aligned homolog conditioning | $0.69 \pm 0.12$ | $8.25 \pm 1.85$ | $0.59 \pm 0.15$ | $0.61 \pm 0.14$ |
| Unaligned homolog conditioning | $0.72 \pm 0.13$ | $8.20 \pm 2.06$ | $0.63 \pm 0.17$ | $0.57 \pm 0.15$ |
| CCMgen | $0.64 \pm 0.13$ | $8.81 \pm 1.83$ | $0.68 \pm 0.17$ | $0.50 \pm 0.08$ |
| Prot-xLSTM | $0.70 \pm 0.14$ | $8.34 \pm 2.15$ | $0.63 \pm 0.18$ | $0.54 \pm 0.15$ |
| EvoDiff-MSA | $0.61 \pm 0.16$ | $9.34 \pm 2.37$ | $0.64 \pm 0.20$ | $0.47 \pm 0.06$ |
| Original queries | $0.76 \pm 0.13$ | $7.88 \pm 1.99$ | - | $0.60 \pm 0.18$ |

**Table S37:** Abbreviation key for datasets defined in the Dayhoff Atlas.

| Dataset acronym | Definition |
| --- | --- |
| UR50 | UniRef50 |
| UR90 | UniRef90 |
| GR | GigaRef |
| GRs | GigaRef, singletons |
| BR | BackboneRef |
| BRu | BackboneRef, unfiltered |
| BRq | BackboneRef, quality-filtered ($\text{scRMSD} < 2 \text{ \AA}$) |
| BRn | BackboneRef, novelty-filtered ($\text{max TM} > 0.5$) |
| HM | homologs (both aligned and unaligned from OpenProteinSet) |
| -c | cooled |

**Table S38:** Model glossary defining the training datasets used to train each model.

| Model name | Training data |
| --- | --- |
| Dayhoff-170m-UR50 | UniRef50 |
| Dayhoff-170m-UR90 | UniRef90 |
| Dayhoff-170m-GR | GigaRef |
| Dayhoff-170m-GRs | GigaRef singletons |
| Dayhoff-170m-UR50-BRu | UniRef50 + BackboneRef (unfiltered) |
| Dayhoff-170m-UR50-BRq | UniRef50 + BackboneRef (quality-filtered) |
| Dayhoff-170m-UR50-BRn | UniRef50 + BackboneRef (novelty-filtered) |
| Dayhoff-3b-UR90 | UniRef90 |
| Dayhoff-3b-GR-HM | GigaRef + OpenProteinSet homologs (aligned + unaligned) |
| Dayhoff-3b-GR-HM-c | GigaRef + OpenProteinSet homologs (aligned + unaligned),<br>then cooldown continuation on UniRef90<br>+ OpenProteinSet homologs (aligned + unaligned) |

**Figure S10:**
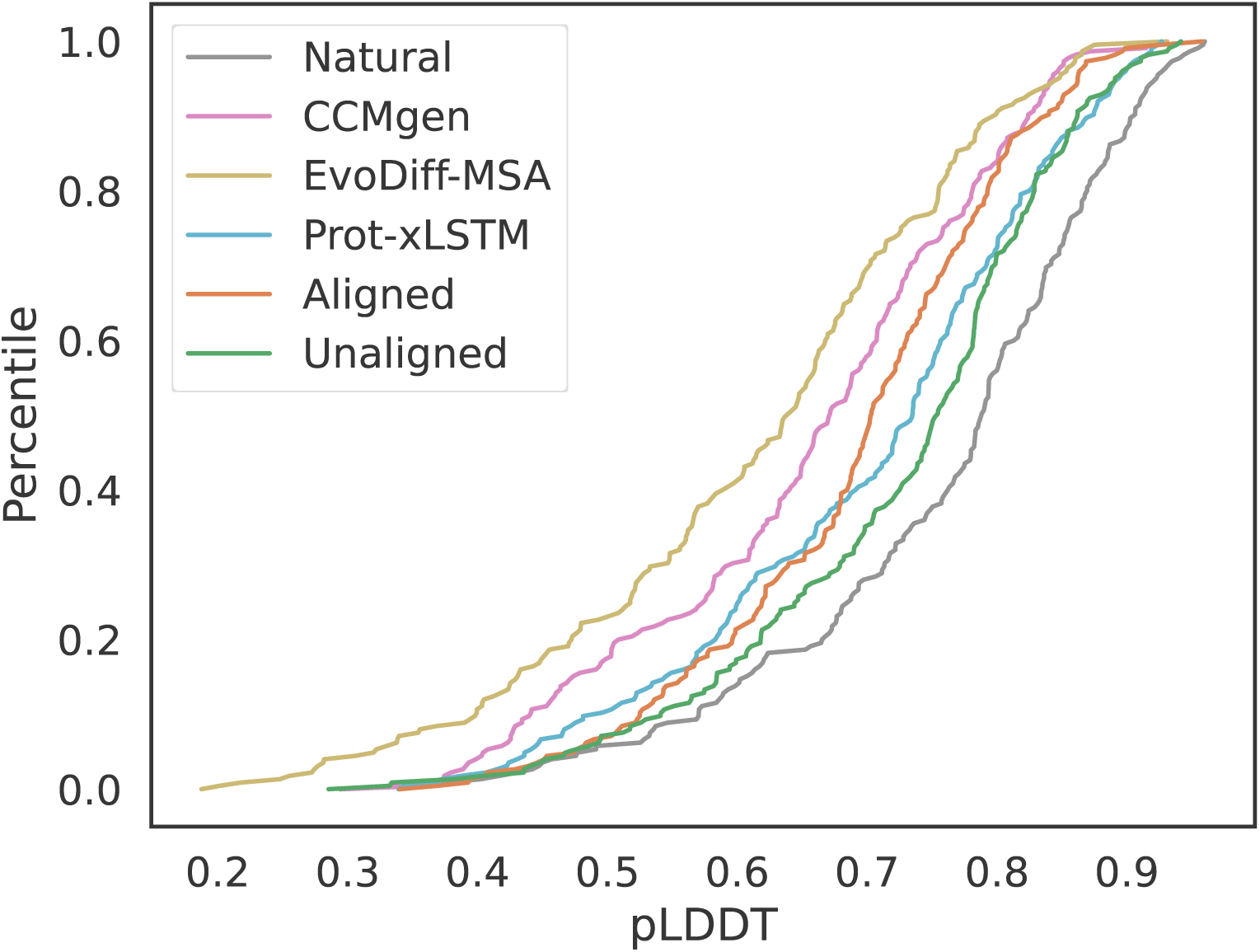
ESMFold pLDDT when regenerating queries conditioned on homologs. *n*=226 sequences per group.

**Figure S11:**
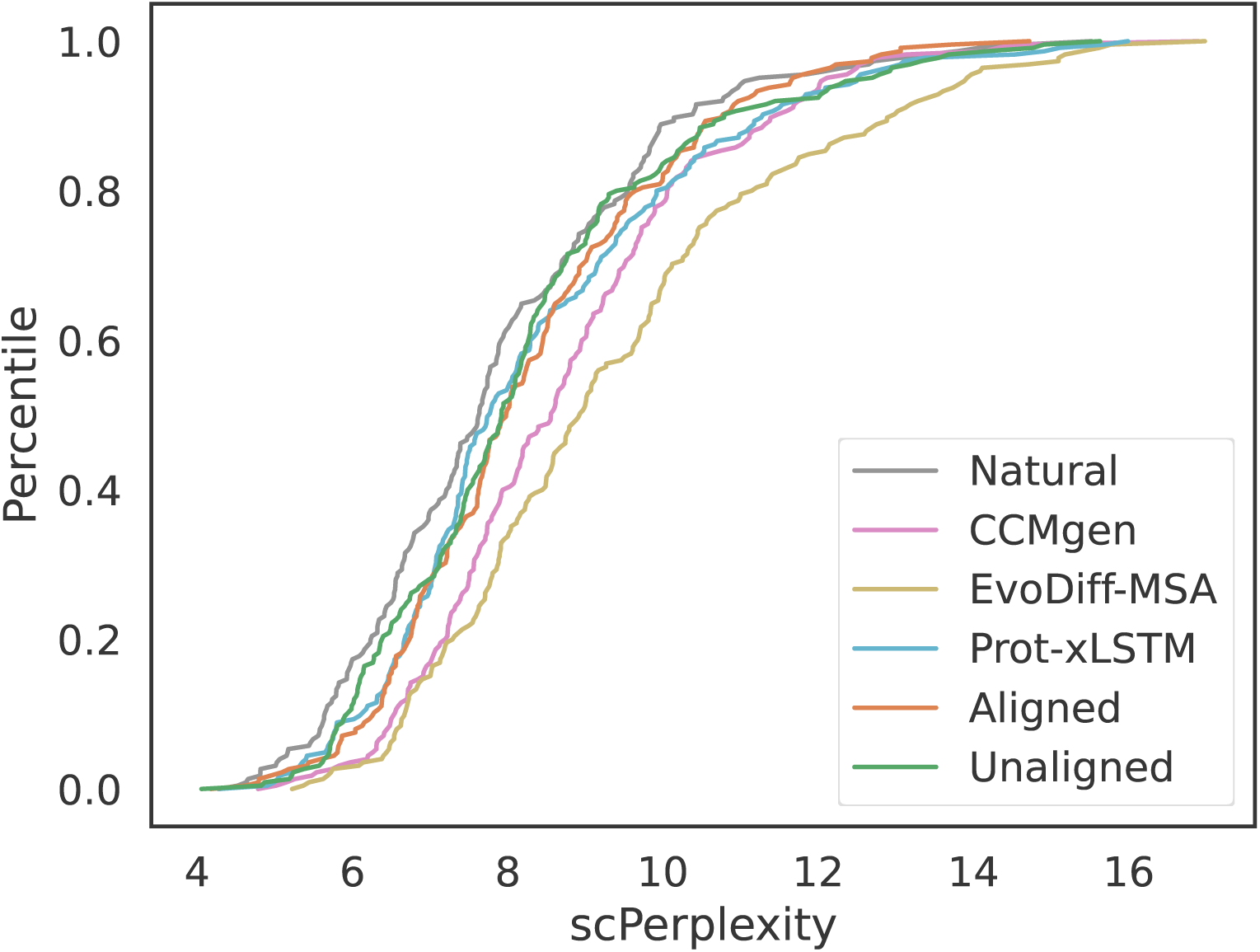
scPerplexity when regenerating queries conditioned on homologs. *n*=226 sequences per group.

**Figure S12:**
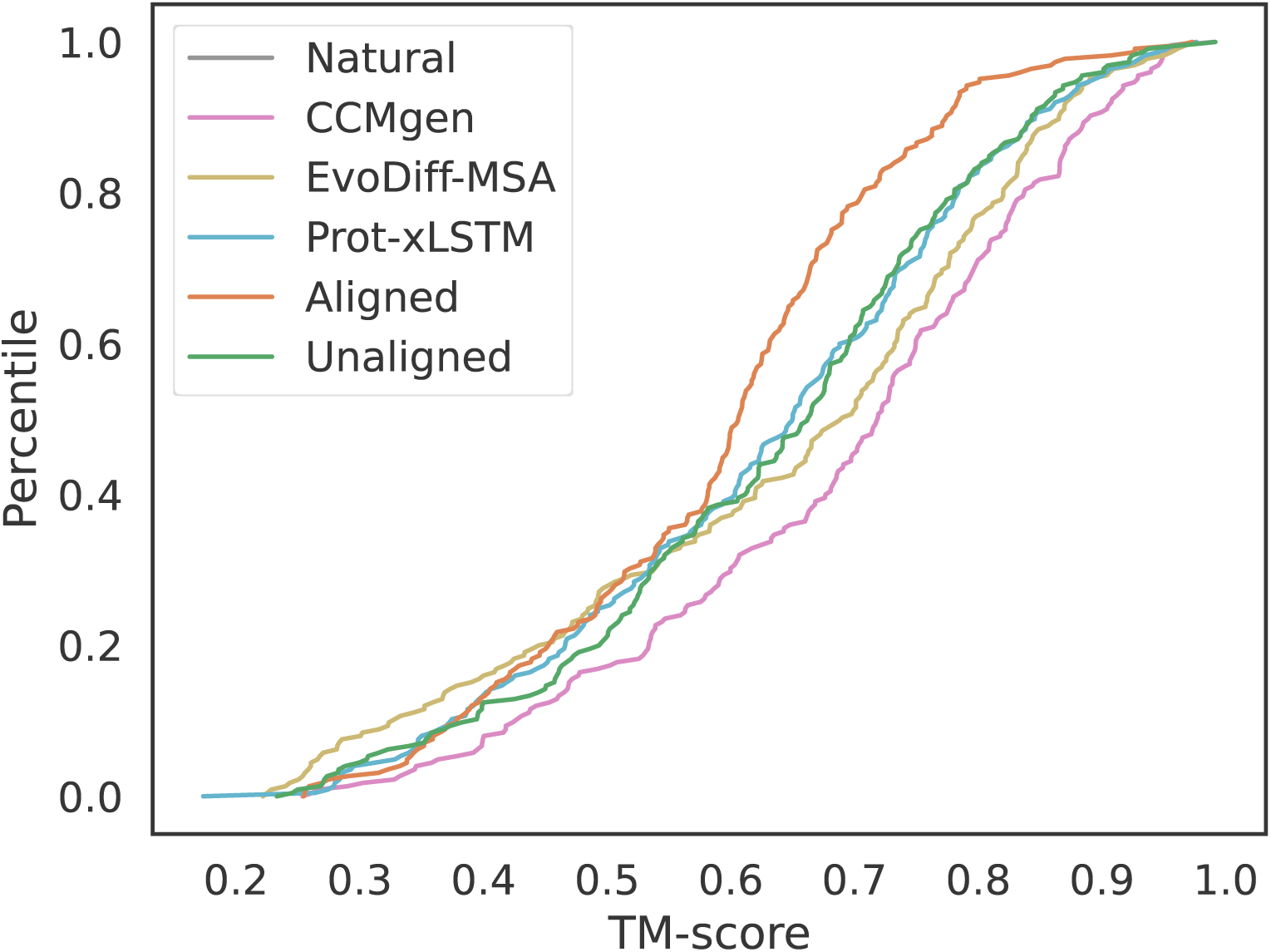
TM-score between ESMFold predictions of original and regenerated queries when regenerating queries conditioned on homologs. *n*=226 sequences per group.

**Figure S13:**
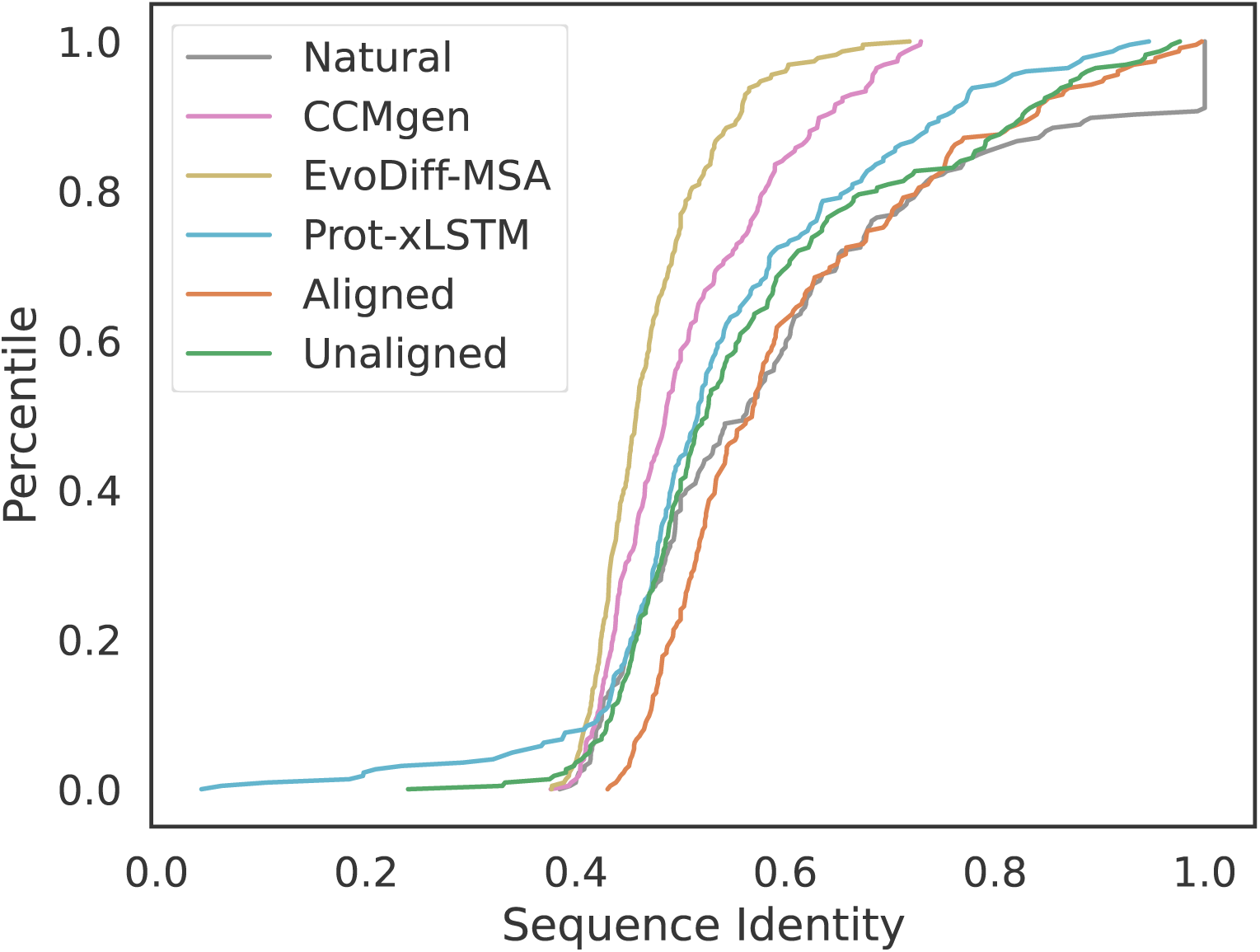
Identity to closest conditioning sequence when regenerating queries conditioned on homologs. *n*=226 sequences per group.

**Figure S14:**
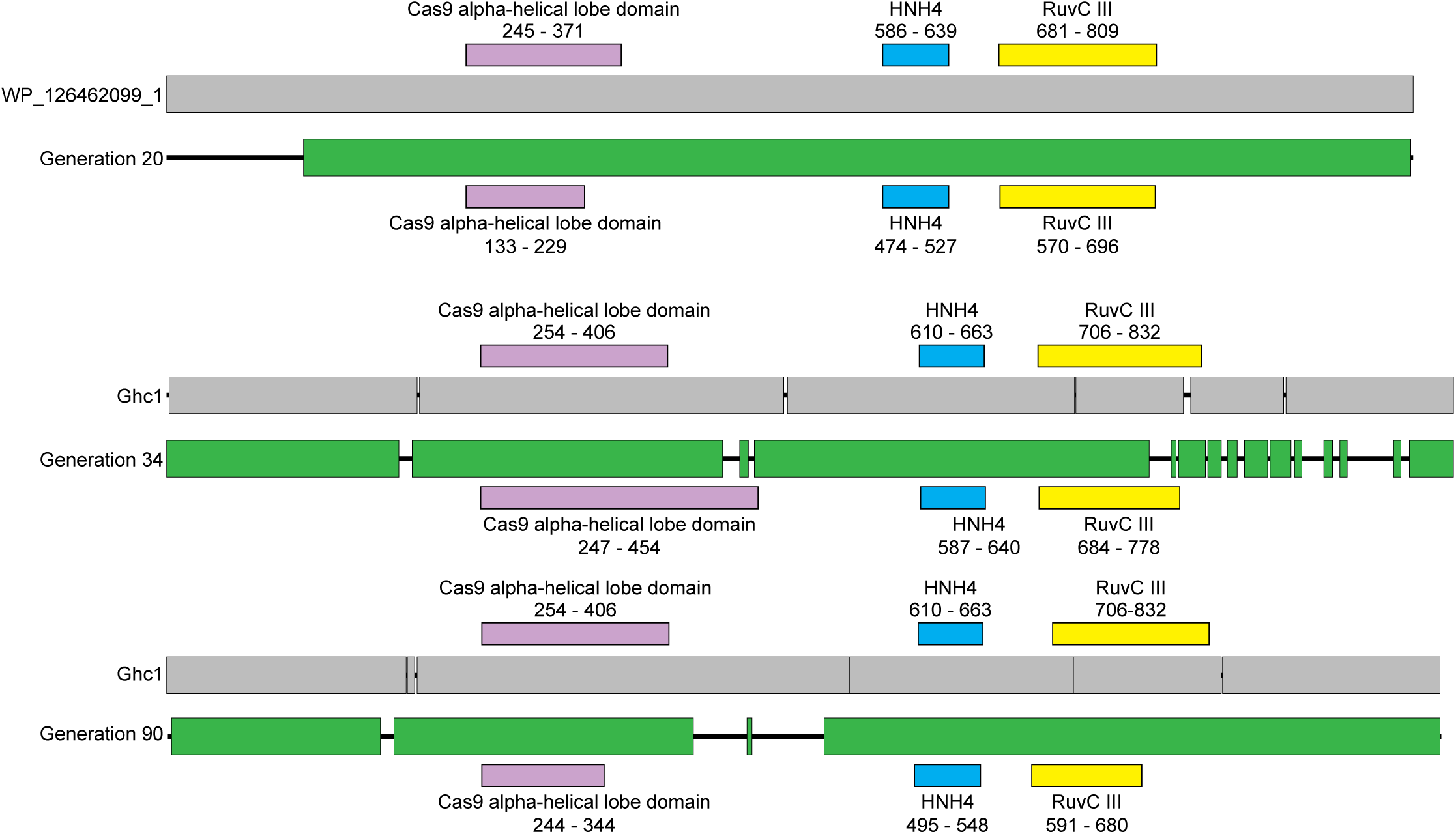
Alignments for shortened Cas9s from Figure 5 relative to the closest natural Cas9 in sequence space.

**Figure S15:**
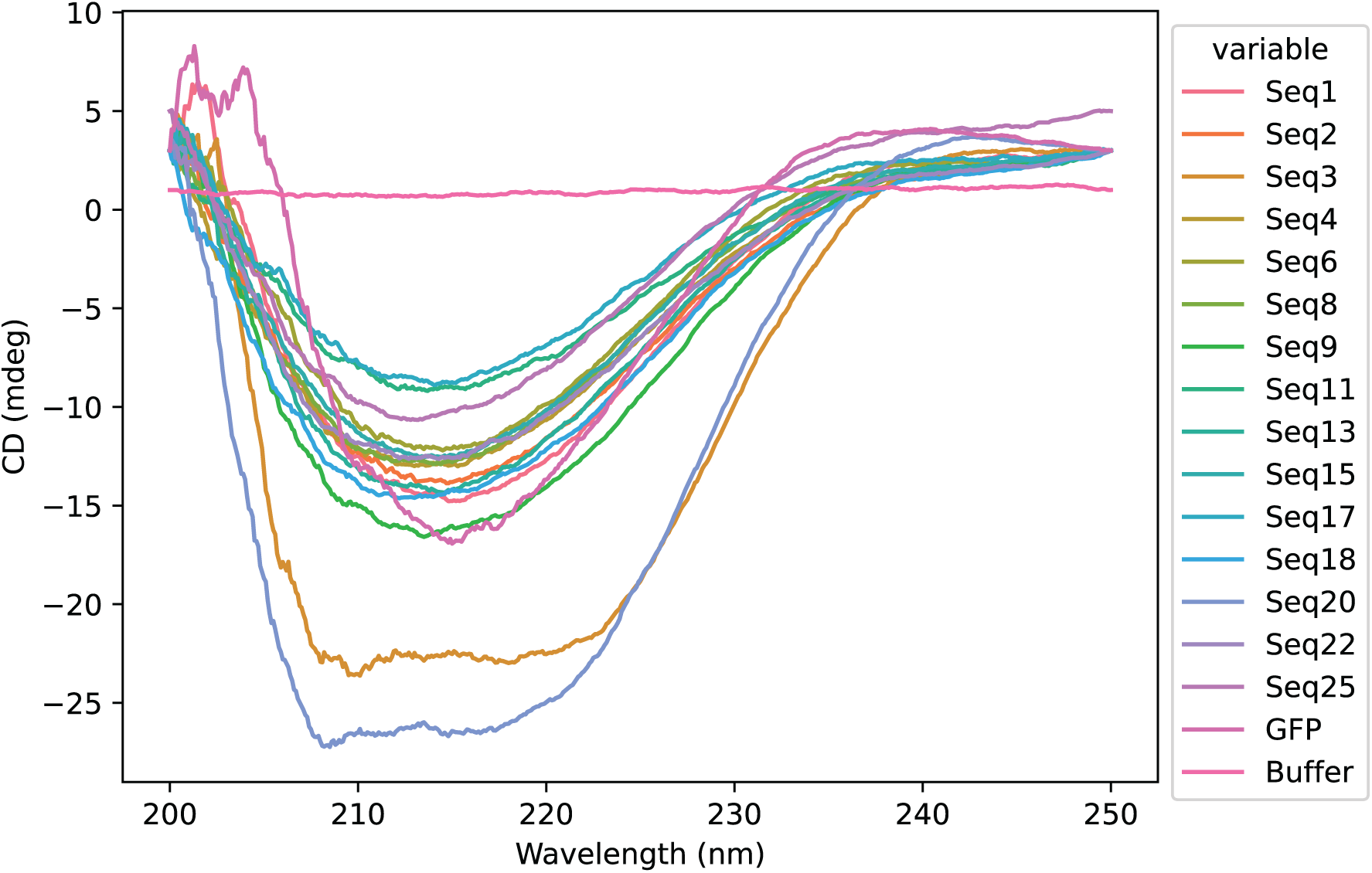
Circular dichroism spectra for sequences generated by Dayhoff-3b-UR90, manually-curated for *in vitro* validation, and successfully expressed in *E. coli*.

